# PML1-Mediated Feedforward Loop Through PI3K and MAPK Axes Drives Endocrine Resistance

**DOI:** 10.1101/2025.05.15.653365

**Authors:** Han Wang, Zixi Yun, Yuan Cao, Xinyue Li, Zhenghao Liu, Chun-Peng Pai, Chen Wu, Jiangan Yue, Gina Lin, Jackie Cai, Belinda Willard, Sichun Yang, Ruth A. Keri, William P. Schiemann, J. Alan Diehl, Hung-Ying Kao

**Author notes:** The authors declare no potential conflicts of interest. To whom correspondence should be addressed: Hung-Ying Kao, Department of Biochemistry, Case Western Reserve University and The Comprehensive Cancer Center of Case Western Reserve University, 10900 Euclid Avenue, Cleveland, OH 44106-4935, USA.

## Abstract

Treatment of estrogen receptor-positive (ER+) breast cancer is significantly hindered by endocrine resistance. We identified PML1 as a key therapeutic entity that can be targeted to overcome resistance. Endocrine-resistant breast cancer cells share three key characteristics: elevated PML1 protein levels, enhanced activity of PI3K, MAPK, or both signaling pathways, and reduced ER activity. We developed a PML1 gene signature that predicts poor prognosis and correlates strongly with PI3K, MAPK, and endocrine resistance gene signatures, as evident by cellular studies, scRNA-seq analysis, and spatial transcriptomics of endocrine therapy-treated tumors. This signature is present in endocrine-resistant breast cancer cells harboring the Y537S ER mutation. We consistently demonstrate high PML1 protein levels across cells resistant to various treatments, including 4-hydroxytamoxifen, fulvestrant, elacestrant, and CDK4/6 inhibitors. Furthermore, treatments with these therapeutic agents or knockdown of *ESR1* mRNA also increase PML1 protein levels. Mechanistically, we show that ER inhibition through fulvestrant treatment activates PI3K and MAPK signaling, which enhance PML1 protein stability and synthesis. PML1 then drives a feedforward loop by stimulating the expression of cytokine and growth factor mRNAs, including *CCL5* and *HBEGF*, further amplifying PI3K and MAPK signaling. Consequently, in endocrine-resistant cells, endocrine therapies, while inactivating ER, paradoxically reinforce this loop through increased PI3K/MAPK activation and PML1 protein accumulation, ultimately compromising therapeutic efficacy. Finally, we demonstrated that arsenic trioxide, an FDA-approved, PML-reducing drug, effectively disrupts this feedforward loop, offering a promising strategy for treating resistant metastatic breast cancer.

**STATEMENT OF SIGNIFICANCE:** Endocrine resistance remains a major obstacle in treating estrogen receptor-positive metastatic breast cancer. Our study identifies PML1 as a central mediator of this resistance, revealing how it maintains a self-reinforcing signaling network through PI3K and MAPK pathways by enhancing the production of cytokines and growth factors. The clinical significance of our findings is threefold: we establish PML1 as a biomarker for therapy resistance, demonstrate its mechanistic role in treatment failure, and show that FDA-approved arsenic trioxide can disrupt PML1-driven resistance. These insights provide a direct path to clinical translation, as combining arsenic trioxide with existing therapies could benefit patients with limited treatment options.

## INTRODUCTION

Estrogen receptor alpha (ER) is a ligand-dependent transcription factor that plays a central role in breast cancer pathogenesis, driving approximately 70% of cases. The standard of care for ER-positive (ER+) breast cancer includes endocrine therapies, including aromatase inhibitors (AIs), selective ER modulators (SERMs), and selective ER downregulators (SERDs) (1–3). While these treatments initially show efficacy, *de novo* and acquired resistance pose significant clinical challenges, particularly in metastatic disease (2,4).

Recent advances in endocrine therapy have transformed the treatment landscape for metastatic breast cancer (MBC), though therapeutic resistance remains a significant clinical challenge. The pairing of fulvestrant (Fulv, a first-generation SERD) with cyclin-dependent kinase 4/6 inhibitors (CDK4/6i) significantly extends progression-free survival in patients with MBC (5–9). More recently, elacestrant (ELA), an oral SERD with pharmacological properties superior to earlier generations, has received FDA approval for second-line treatments and shows promising antitumor activity against Y537S-mutant tumors (10–13). However, patients with CDK4/6i and ELA may experience resistant tumors (14–18). Thus, understanding resistance mechanisms to CDK4/6i and ELA remains crucial for further therapeutic advancement.

Targeted metastatic breast cancer (MBC) sequencing has revealed mutations linked to resistance(8,19–27), and ER mutations, particularly Y537S and D538G, represent a substantial fraction of resistant MBC cases. Resistance also frequently emerges through dysregulation of key mitogenic and survival signaling cascades, specifically the PI3K/Akt/mTOR and Raf/MAPK signaling pathways. Despite the identification of pathways, their underlying molecular mechanisms remain incompletely understood.

We previously identified PML1 as the predominant PML isoform across breast cancer subtypes, and its elevated expression correlates with poor prognosis in ER+ breast tumors. Unlike PML4, which acts as a tumor suppressor, PML1 exhibits oncogenic properties in ER+ breast cancer by upregulating genes associated with breast cancer stem cells (BCSCs) (28). Notably, PML1 overexpression promotes Fulv resistance, and the *PML* gene is amplified across all metastatic breast cancer (MBC) subtypes, including a significant proportion of ER+ cases. These observations suggest a potential role for PML1 in therapy resistance and disease progression.

In this study, we report elevated PML1 protein levels in endocrine-resistant cells. We identified an inverse relationship, in which PML1 levels increase as ER signaling is suppressed via endocrine therapies, CDK4/6 inhibition, or reduction of ER protein by knockdown. This pattern, further supported by our clinical correlates and cellular analyses demonstrating increased PML1 expression in more aggressive breast cancers, such as triple-negative breast cancer (TNBC), suggests that PML1 upregulation may be an adaptive response to treatment and plays a key role in maintaining cancer cell survival during therapy.

## RESULTS

### A PML1 Gene Signature Correlates With PI3K, MAPK, And Endocrine Resistance Signaling

Following the identification of PML1 as the predominant isoform in ER+ breast cancer cells (28), we examined its functional role through comprehensive multi-omics analysis and cellular studies. We analyzed chromatin immunoprecipitation followed by sequencing (ChIP-seq) data and found that PML1/ER overlapped gene promoters are enriched in EGFR, ERBB2, and PI3K-Akt signaling pathways (Fig. 1A). Moreover, PML-bound promoters are also enriched in EGFR/ERBB2/RAF/MAPK and PI3K/Akt/mTOR pathways (Fig. 1B, Table S1). Notably, many PML1/ER overlapped promoters are associated with endocrine resistance. Functionally, *PML* depletion reduced the phosphorylation of EGFR, ERK, mTOR, and p70 (p70S6K) (Fig. 1C). Conversely, PML1 overexpression enhanced their phosphorylation, increased Fulv resistance, enhanced colony formation capacity, and supported estrogen-independent growth (Fig. S1A-C). These results demonstrated PML1’s role in activating PI3K and MAPK signaling pathways. RNA-seq analysis in *Pml* knockout mouse embryonic fibroblasts (MEFs) (29) further confirmed PML1’s role in regulating MAPK, EGFR, RAF, and ERBB2 pathway activation (Fig. S1D).

**Figure 1.**
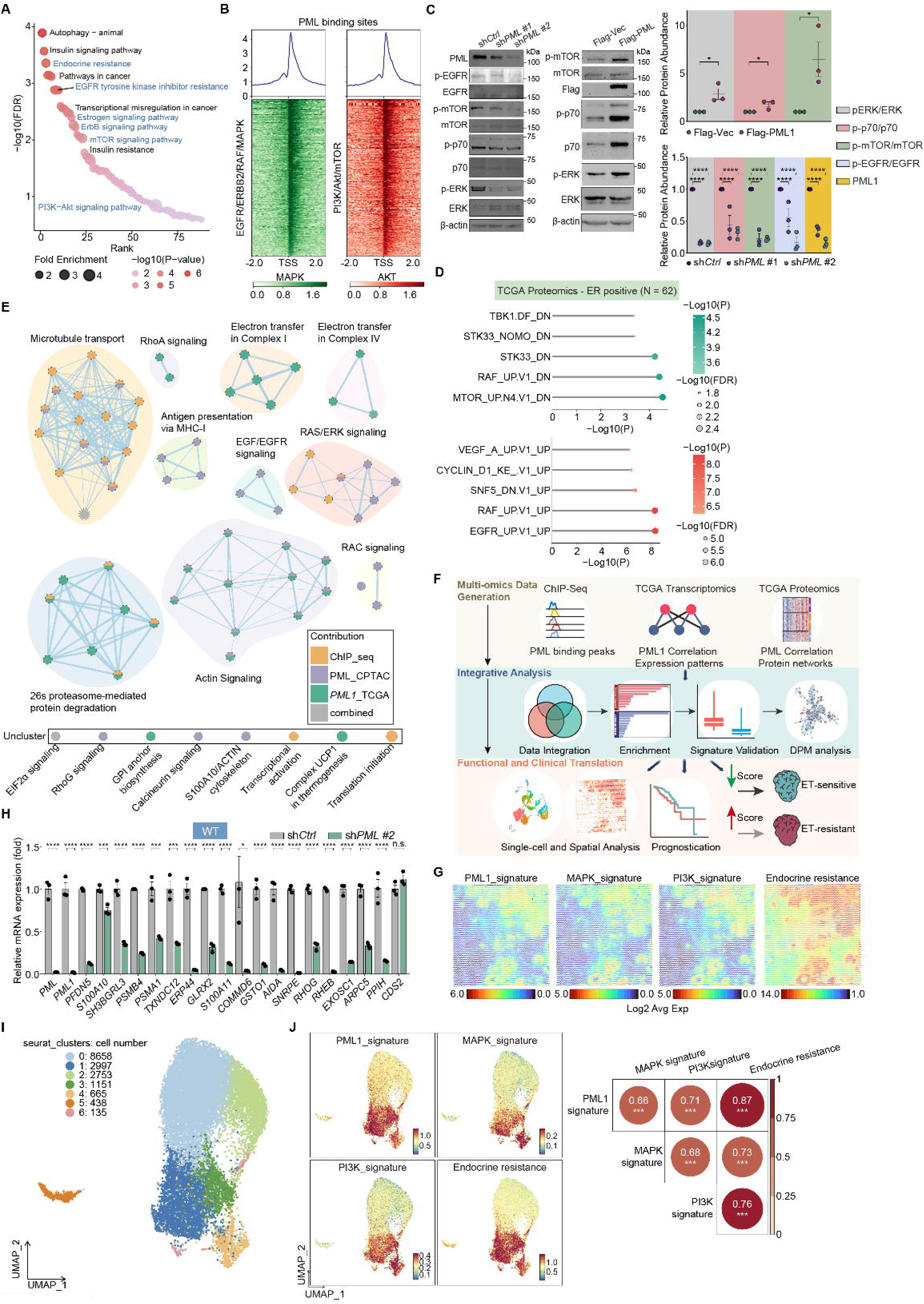
PML1 activates PI3K and MAPK signaling pathways. ***A,*** A pathway enrichment analysis plot of MCF-7 ChIP-seq data reveals PML/ER overlapped binding on promoter regions of genes involved in endocrine resistance, EGFR, estrogen signaling, ERBB2, mTOR, and mTOR signaling pathways. ***B,*** Heat maps depicting PML-binding sites on genes in EGFR/ERBB2/RAF/PI3K and MAPK/Akt/mTOR pathways. ***C,*** *Left*, Western blot analyses demonstrate PML1’s role in MEK and mTOR signaling. *PML* knockdown reduces pathway activity, while PML1 overexpression enhances signaling through these pathways, as shown in stable cell lines expressing FLAG-vector (F-vec) or FLAG-PML1 (F-PML1). *Right*, quantification of Western blots from two repeats. ***D,*** A gene set enrichment analysis of proteomic analysis of ER+ breast tumors (TCGA, N=62) demonstrates a strong correlation between PML protein abundance and pathway activation. Protein sets showing strong positive (top) or negative (bottom) correlation with PML (threshold |r| > 0.25) were evaluated using oncogenic signature gene sets. ***E,*** Integrative Active Pathway analysis from ChIP-seq, PML_TCGA, PML1_TCGA, and combined analyses (node coloring) identifies significant biological processes, including EGF/EGFR signaling, through multi-omics data integration. At the bottom, an "Uncluster" section shows additional pathways that were not grouped into the main clusters. ***F,*** Schematic workflow for generating the PML1 gene signature using multi-omics data integration. Integration of ChIP-seq (PML-bound genes), proteomics (proteins positively correlating with PML abundance), and RNA-seq (transcripts correlating with *PML1* expression in ER+ breast tumors) identifies high-confidence PML1 direct target genes. ***G,*** Spatial transcriptomic analysis of a luminal B breast tumor stage II reveals four heatmaps representing PML1, PI3K, MAPK, and endocrine resistance with scores. ***H,*** RT-qPCR validation confirms differential expression of PML1 signature genes following *PML* knockdown, with *CDS2* as a non-target control gene. ***I-J,*** UMAP clustering of scRNA-seq analyses of an E2-accelerating Pdx breast tumor. (***I***) reveals a strong correlation between PML1, PI3K, MAPK, and endocrine therapy-resistance gene signatures (***J***).

Analysis of the ER+ breast tumor TCGA dataset identified proteins significantly correlated with PML abundance (|r| > 0.25) (Table S1). Enrichment analysis of oncogenic signatures among these proteins revealed that the RAF and EGFR activation signatures showed the strongest positive correlation with PML protein levels (Fig. 1D, Table S1). Conversely, mTOR and RAF inhibition signatures exhibited the strongest negative correlation. Active Pathway analysis (30) of PML1, integrating multiple data types, revealed the involvement of significant biological processes, including EGF/EGFR and RAS/ERK signaling (Fig. 1E).

To validate these findings in clinical samples, we developed a 46-gene PML1 gene signature by integrating ChIP-seq, TCGA proteomics, and TCGA RNA-seq data (Fig. 1F, S2-5, Table S1). Interestingly, in addition to MCF-7 cells, PML engages these gene promoters in GM12878 and K562 cells (Fig. S6). Spatial transcriptomics analyses of luminal breast tumors (31) demonstrated significant overlap between PML1, PI3K (32), MAPK (33), and endocrine resistance gene signatures (34)(Fig. 1G & S7, Table S2). We further confirmed considerable downregulation of these signature genes and BCSC-associated genes following total *PML* or *PML1*-specific knockdown via RT-qPCR validation (Fig. 1H & S8). UMAP clustering of single-cell RNA-seq (scRNA-seq) data from ER+ Pdx breast tumor (35)(Fig. 1I & S9) revealed strong correlations between PML1 gene signatures and PI3K, MAPK, and endocrine resistance signatures across 7 distinct epithelial cell populations (Fig. 1J). These findings establish a mechanistic link between PML1 and PI3K and MAPK pathway activation, supporting the role of PML1 in endocrine therapy resistance.

### PML1 Protein Levels Are Elevated In Endocrine Therapy-Resistant Cells, Associated With Fulv Resistance, And Inversely Correlated With ER Protein Abundance

The observation that PML1 promotes Fulv resistance in ER+ breast tumors (28) prompted us to investigate PML1 protein expression across three independent models of Fulv-resistant cells: *ESR1* mutation-knock-in (KI) cells (36,37), cells grew independently of estrogen (E2) (38,39), and Fulv-resistant MCF-7 cell variants developed in our laboratory. Cells harboring the prevalent resistance-conferring *ESR1* mutations (Y537S and D538G), ^Y537S^ER- and ^D538G^ER-KI MCF-7 and T47D cells, exhibited significantly higher PML1 protein levels compared to ^WT^ER-KI controls (Fig. 2A). Immunofluorescence microscopy confirmed this pattern, with ^Y537S^ER-KI cells showing the highest PML abundance, followed by ^D538G^ER and ^WT^ER-KI MCF-7 cells (Fig. 2B).

**Figure 2.**
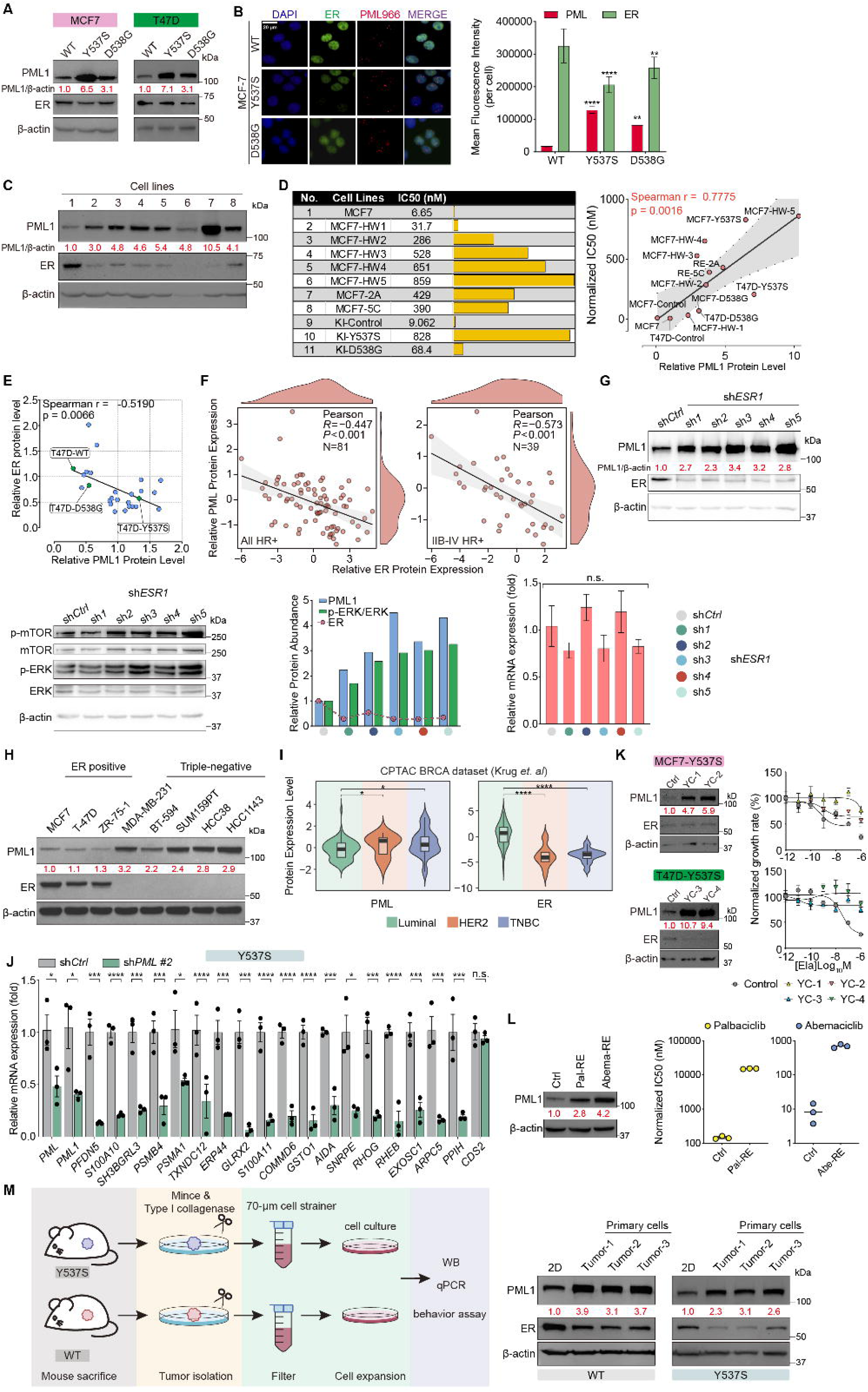
PML1 protein levels are associated with endocrine resistance and inversely correlate with ER levels. ***A,*** PML1 protein levels are elevated in ^Y537S^ER- and ^D538G^ER-KI cells compared to ^WT^ER-KI MCF-7 and T47D cells. ***B,*** Representative (*left*) images and quantification analyses (*right*) reveal elevated PML protein levels in ^Y537S^ER- and ^D538G^ER-KI cells compared to ^WT^ER-KI MCF-7 cells. ***C,*** PML1 protein levels are significantly higher in Fulv-resistant MCF-7 cell variants than in parental cells. ***D,*** *Left*, IC_50_ values across multiple Fulv-resistant MCF-7 cell variants demonstrate varying degrees of acquired resistance. *Right*, Correlation analysis shows a strong positive relationship between PML1 protein levels and Fulv resistance. ***E,*** PML1 and ER protein abundance are inversely correlated in MCF-7 and T47D cell variants. ***F,*** Quantitative analysis of clinical samples reveals the inverse correlation between PML and ER protein levels in all-stage (*left*) and advanced (*right*) ER+ breast cancer cohorts. ***G,*** Increased PML1 protein levels coincide with Akt and MEK activation in *ESR1* knockdown MCF-7 cells, shown by elevated phospho-ERK and mTOR levels (*left*). Quantitative analysis reveals differences in ER, PML1, and pERK levels between control and *ESR1* knockdown cells (*middle*). *ESR1* knockdown does not affect *PML* mRNA expression (*right*). ***H,*** PML1 protein expression is significantly higher in TNBC cells than in ER+ breast cancer cells. ***I,*** Analysis of the CPTAC data reveals elevated PML protein levels in HER2 (N=14) and TNBC (N=29) tumors compared to luminal (N=74) tumors. An inverse relationship between PML and ER protein levels was observed across breast cancer subtypes. ***J,*** The expression of the PML1 signature genes is significantly reduced in *PML* knockdown ^Y537S^ER-KI MCF-7 cells. ***K,*** *Left*, PML1 protein abundance is upregulated in ELA-resistant ^Y537S^ER-KI MCF-7 (*top*) and T47D (*bottom*) cells. *Right,* ELA-resistant cells exhibit higher IC50. ***L,*** *Top*, PML1 protein abundance is upregulated in CDK4/6i-resistant MCF-7 cells. *Bottom*, Pal- and Abema-resistant cells exhibit higher IC50. ***M,*** PML1 protein levels are elevated in xenografted MCF-7 tumors compared to cultured cells, whereas ER protein levels show the opposite pattern.

The MCF-7 2A and 5C variants were isolated based on their ability to grow independently of E2, with the 2A variant also showing proliferation in the presence of 4-hydroxytamoxifen (4-OHT) (38). We established additional Fulv-resistant variants (HW1-5) by selecting cells capable of growing in various Fulv concentrations. Across all resistant cell variants, we observed at least threefold higher PML1 protein levels than parental cells (Fig. 2C), and knockdown of *PML* significantly reduced Fulv IC50 (Fig. S10A). We observed a positive correlation with Fulv IC_50_ and PML1 protein levels (Fig. 2D). Intriguingly, we identified an inverse relationship between PML1 and ER protein abundance (Fig. 2E). This correlation was validated in proteomic data from ER+ breast tumors (Fig. 2F). Additionally, *ESR1* knockdown increased PML1 protein levels while simultaneously activating Akt and MEK signaling pathways (Fig. 2G), reinforcing the connection between PML1 abundance and PI3K and MAPK pathway activation. However, *ESR1* knockdown had no or little effect on *PML* mRNA levels. Knockdown of *ESR1* in ^Y537S^ER-KI cells also increased PML1 protein levels (Fig. S10B). Moreover, we observed significantly higher PML1 protein levels in triple-negative breast cancer (TNBC) cell lines than in ER+ breast cancer cells (Fig. 2H). Analyses of proteomics data from breast tumor samples further supported these findings, demonstrating elevated PML protein levels in TNBC tumors relative to ER+ breast tumors (Fig. 2I). Collectively, these results establish an inverse correlation between PML1 and ER protein levels in both cell lines and clinical samples. Moreover, the depletion of total *PML* by knockdown significantly reduced PML1 signature gene expression in ^Y537S^ER-KI cells (Fig. 2J).

We extended our study to examine resistance to other therapeutic agents by establishing MCF-7 cell variants resistant to elacestrant (ELA) and CDK4/6 inhibitors, palbociclib (Pal), and abemaciclib (Abema). ELA is a SERD and has recently been FDA-approved as a second-line treatment, particularly for patients with the Y537S mutation. Our results showed that PML1 protein levels and the IC50 were significantly elevated in these resistant variants (Fig. 2K-L) compared to parental cells. Loss of *PML* by knockdown reduced ELA IC50 (Fig. S10C). We also observed elevated PML1 protein levels (Fig. 2M), BCSC-associated gene expression, and cancer cell functional characteristics (Fig. S10D-G) in primary tumor cells compared to those grown in 2D culture.

These findings establish that (1) PML1 positively regulates PI3K and MAPK signaling activation, (2) PML1 protein levels are consistently elevated in therapy-resistant cells, (3) the PML1 gene signature is strongly associated with PI3K and MAPK gene signatures in ER+ breast tumors, and (4) PML1 levels generally show an inverse correlation with ER protein abundance. These observations reveal an intriguing interplay between PML1, ER protein levels, and therapy resistance in ER+ breast cancer, highlight the association between PML1 protein levels and PI3K/MAPK pathway activation, and suggest a role of PML1 in promoting endocrine resistance.

### Fulv Treatment Induces PML1 Protein Accumulation And PI3K/MAPK Pathway Activation

Given the inverse correlation between PML1 and ER protein levels, we investigated the effects of Fulv on PML1 protein levels. Western blot analysis demonstrated time-dependent PML1 protein accumulation and PI3K and MAPK signaling activation following Fulv treatment in all three KI cell variants (Fig. 3A). RNA-seq analysis revealed Fulv-induced activation of EGFR, ERBB2, Raf, and MEK signaling pathways across all three *ESR1* KI cell lines (Fig. 3B). Specifically, Fulv triggered MEK-ERK and mTOR-p70 activation, with increased ERK and p70 phosphorylation occurring within the first hour preceding PML1 protein accumulation (Fig. S11A). Immunofluorescence microscopy confirmed increased PML staining intensity and reduced ER staining following Fulv treatment, revealing an inverse relationship between PML and ER staining intensity (Fig. 3C). We also demonstrated that Fulv induces PML1 protein expression in T47D KI cell variants (Fig. 3D) and in-house Fulv-resistant cells (Fig. S11B).

**Figure 3.**
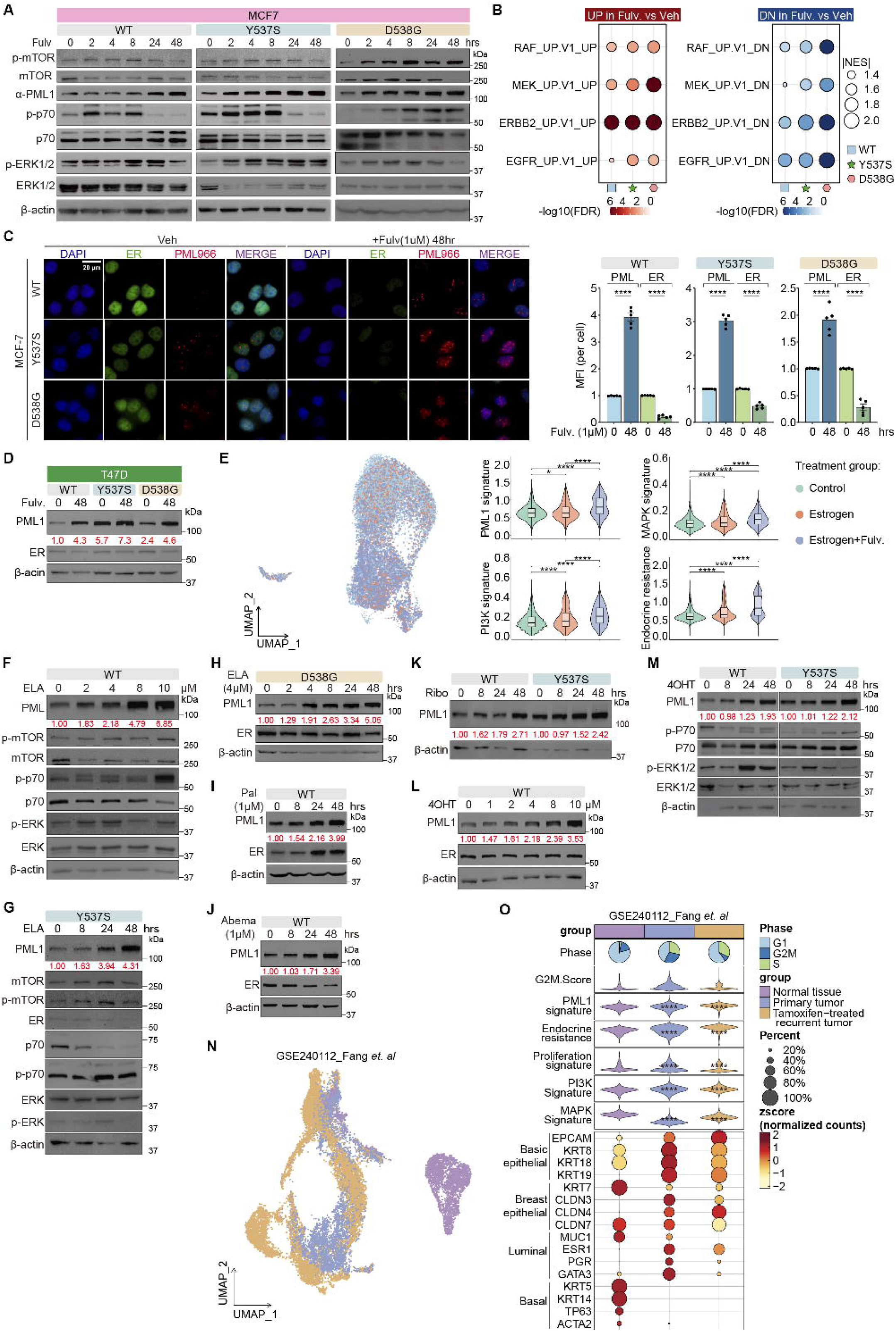
Fulv, ELA, CDK4/6i, and 4-OHT induce PML1 protein accumulation. ***A,*** Fulv treatment leads to activation of MEK-ERK and Akt-mTOR signaling pathways and PML1 protein accumulation. ***B,*** Dot plots based on RNA-seq analyses show Fulv-induced activation of RAF, MEK, ERBB2, and EGFR signaling cascades in ^WT^ER-KI, ^Y537S^ER-KI, and ^D538G^ER-KI MCF-7 cell variants following 48-hour Fulv (1 μM) treatment. ***C,*** *Left*, Representative images and quantification analyses of immunofluorescence microscopy in MCF-7 cells show Fulv-induced increases in PML protein abundance. *Right*, PML and ER intensity were quantified based on six fields over 100 cells for each sample. ***D,*** Fulv induces PML1 protein expression in ^WT^ER-, ^Y537S^ER-, and ^D538G^ER-KI T47D cells. ***E,*** *Top*, A UMAP integrating scRNA-seq data from a Pdx model of luminal B breast cancer shows the distribution of different cell clusters of treatments. *Bottom*, Estrogen and Fulv enhance PML1, PI3K, MAPK, and endocrine resistance gene signatures in E2-accelerating Pdx breast tumors. ***F-H***, ELA treatment induces PML1 protein levels in a dose-(***F***) and time-dependent manner in ^Y537S^ER-(***G***) and ^D538G^ER-KI (***H***) MCF-7 cells. ***I-K,*** Pal (***I***), Abema (***J***), and Ribo (***K***) treatments upregulate PML1 protein levels. ***L-M,*** 4-OHT treatment increases PML1 protein abundance in a dose-(***L***) and time-dependent (***M***) manner (2 μM). ***N,*** UMAP of scRNA-seq analyses of a luminal A tumor shows cell distribution of normal tissues, primary ER+ breast tumors, and TAM-treated recurrent tumors. ***O***, A comprehensive scRNA-seq analysis of TAM-resistant ER+ breast tumor that compares molecular and cellular characteristics across the progression from normal breast tissue to primary and tamoxifen-resistant recurrent tumors. The analysis indicates that PML1, PI3K, MAPK, and endocrine resistance gene signatures are upregulated in TAM-treated tumor tissues compared to primary tumors. Marker genes for different cell types are shown.

The effect of Fulv on the PML1 gene signature was further assessed by analyzing scRNA-seq of a Pdx model of luminal breast tumors (34), comparing tumors treated with vehicle control, E2, or E2 plus Fulv, which revealed Fulv-induced enhancement of the PML1 gene signature and activation of PI3K and MAPK signaling pathways across 7 distinct cell populations (Fig. 3E & S9).

Investigation of ELA showed similar time- and dose-dependent (Fig. 3F-H) increases in PML1 protein levels. Given recent FDA approval of the combined treatments of CDK4/6 inhibitors (CDK4/6i) with Fulv in MBCs (16,17,40), we examined the effects of Pal, Abema, and ribociclib (Ribo) on PML1 protein levels. All three CDK4/6 inhibitors induced PML1 protein accumulation (Fig. 3I-K). Additionally, 4-OHT induced PML1 protein accumulation in both dose-(Fig. 3L) and time-dependent (Fig. 3M) manners. scRNA-seq analysis further supported the conclusion that PML1, PI3K, MAPK, and endocrine resistance gene signatures are elevated in tamoxifen (TAM)-resistant tumors (41) compared to primary tumors across 14 cell clusters (Fig. 3N-O & S12).

These results demonstrate that multiple classes of endocrine therapies (Fulv, ELA, CDK4/6i, and 4-OHT) induce PML1 protein accumulation and PML1, PI3K, MAPK, and endocrine resistance gene signatures. This pattern suggests a central role for PML1 in therapeutic resistance mechanisms.

### PML Depletion Or Inhibition Enhances Fulv Efficacy In ER+ Breast Cancer

To investigate the functional significance of elevated PML1 in Fulv resistance, we measured the impact of *PML* depletion on Fulv sensitivity. *PML* knockdown significantly enhanced Fulv sensitivity in ^Y537S^ER-KI cells (Fig. 4A). Arsenic trioxide (ATO), which induces PML protein degradation and inhibits mTOR (Fig. 4B), demonstrated synergistic effects with Fulv in ^Y537S^ER-KI cells (Fig. 4C-D), as quantified by the Chou-Talalay method (42) and Fulv’s IC_50_ (Fig. 4E). Similar synergistic effects were observed in ^WT^ER- and ^D538G^ER-KI cells (Fig. S13A-E).

**Figure 4.**
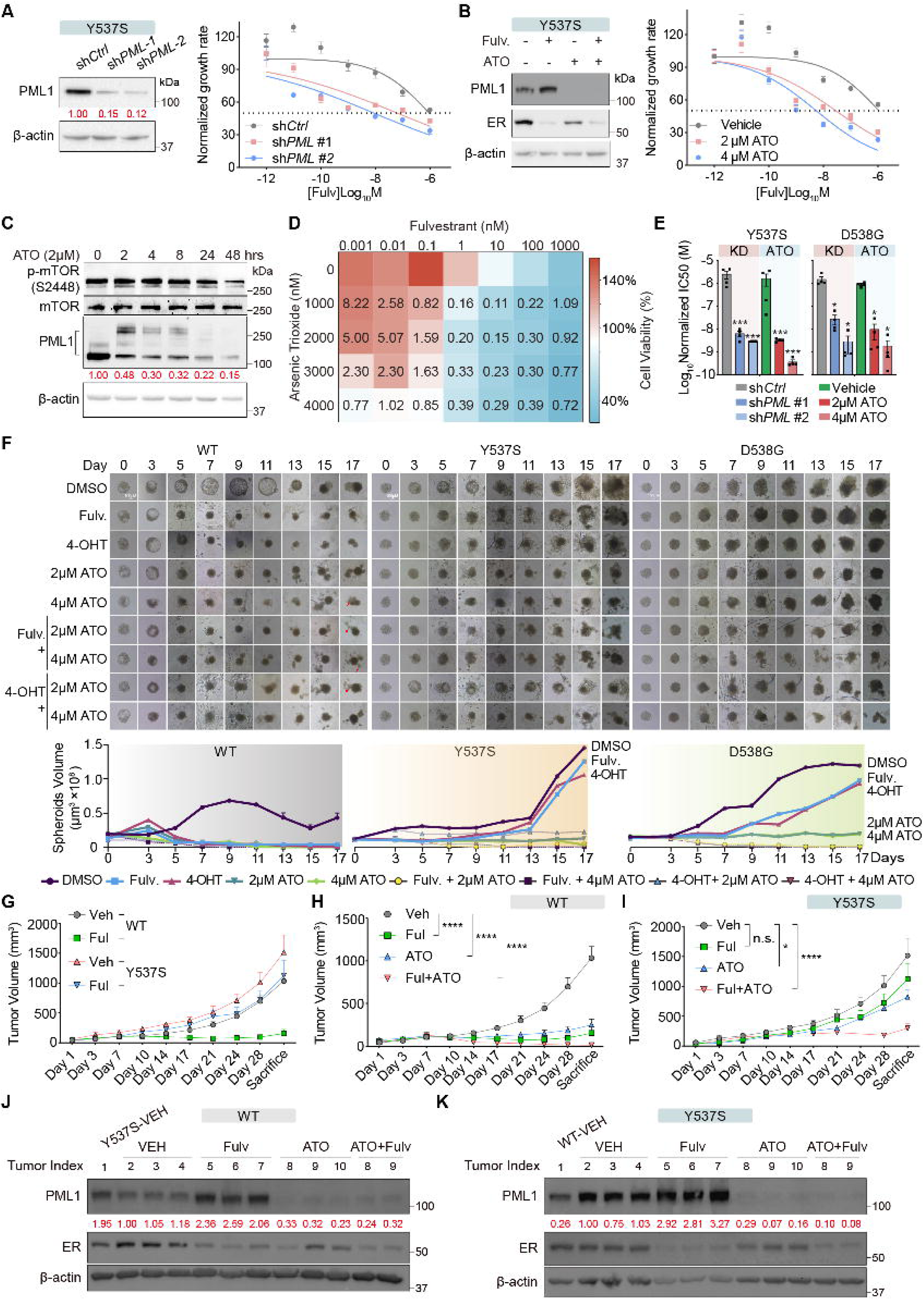
ATO treatment enhances Fulv’s anti-cancer efficacy *in vitro* and in orthotopic xenograft models. ***A,*** Cell viability assays demonstrating enhanced Fulv sensitivity in ^Y537S^ER-KI cells following *PML* knockdown. ***B,*** Western blot analysis showing ATO-induced PML1 degradation and reduced mTOR activation. ***C,*** Western blot demonstrating Fulv (1 μM) and ATO (2 μM) effects on PML1 and ER protein levels in ^Y537S^ER-KI. Sumoylated PML1 species migrate at a molecular weight of > 150 Kd. ***D,*** Drug synergy analysis of Fulv/ATO combinations using the Chou-Talalay method with Combination Index (CI) values. ***E,*** *PML* knockdown or ATO significantly reduced Fulv’s IC_50_ in ^Y537S^ER- and ^D538G^ER-KI MCF-7 cells. ***F,*** The effects of vehicle, 4-OHT, Fulv, ATO, or their combinations on the growth of tumor spheroids. Spheroid (top) volume (bottom) was quantified by imaging analysis. ***G,*** Xenografted tumors derived from ^Y537S^ER-KI cells demonstrated resistance to Fulv treatment. ***H-I.*** ATO enhances Fulv-mediated tumor growth inhibition in orthotopically xenografted ^WT^ER-(***H***) and ^Y537S^ER-KI (***I***) MCF-7 cells. Tumor volume measurements demonstrate an enhanced anti-tumor effect of the Fulv/ATO combination versus single agents (N > 10, two-way ANOVA with Tukey’s multiple comparisons test). ***J-K,*** Western blot analyses of xenograft tumors showing Fulv- and ATO-mediated reduction of ER and PML1 proteins in ^WT^ER- (***J***) and ^Y537S^ER-KI (***K***) MCF-7 cells.

Next, we examined the effects of Fulv, ATO, or their combination on the growth of tumor spheroids in the KI cells. Intriguingly, we found that PML1 protein levels in tumor spheroids are elevated compared to cultured 2D cells (Fig. S13F). While Fulv, 4-OHT, and ATO alone effectively reduced spheroid sizes in ^WT^ER-KI cells, they had little or no effect on the growth of tumor spheroid in ^Y537S^ER- or ^D538G^ER-KI cells (Fig. 4F). However, combining ATO with either Fulv or 4-OHT significantly reduced spheroid sizes in both ^Y537S^ER- or ^D538G^ER-KI cells, consistent with the results of cellular studies.

In orthotopic xenograft studies, tumors from ^Y537S^ER-KI cells grew larger than those from ^WT^ER-KI cells (Fig. 4G). While Fulv and ATO significantly reduced tumor growth in ^WT^ER-KI cells alone or in combination (Fig. 4G-H, S13G-H), they showed limited efficacy as single agents against ^Y537S^ER-KI tumors. Importantly, ATO significantly enhanced Fulv’s anticancer activity against ^Y537S^ER-KI tumors. Moreover, analyses of tumor samples confirmed our cellular findings: Fulv treatment induces PML1 and reduces ER protein abundance. While ATO treatment substantially reduces PML1 protein levels, the Fulv-ATO combination eliminated ER and PML1 protein expression (Fig. 4J-K).

### The *PML* Gene Is Frequently Amplified In Metastatic Breast Cancer (MBC) Cases

Our findings that PML1 promotes Fulv resistance and is elevated in resistant cells prompted us to investigate *PML* gene alterations in ER+ MBCs. Whole genome sequencing (43) analyses revealed *PML* gene amplification in 14% of ER+ MBC cases (Fig. 5A) and 20% of all MBC cases (Fig. S14A).

**Figure 5.**
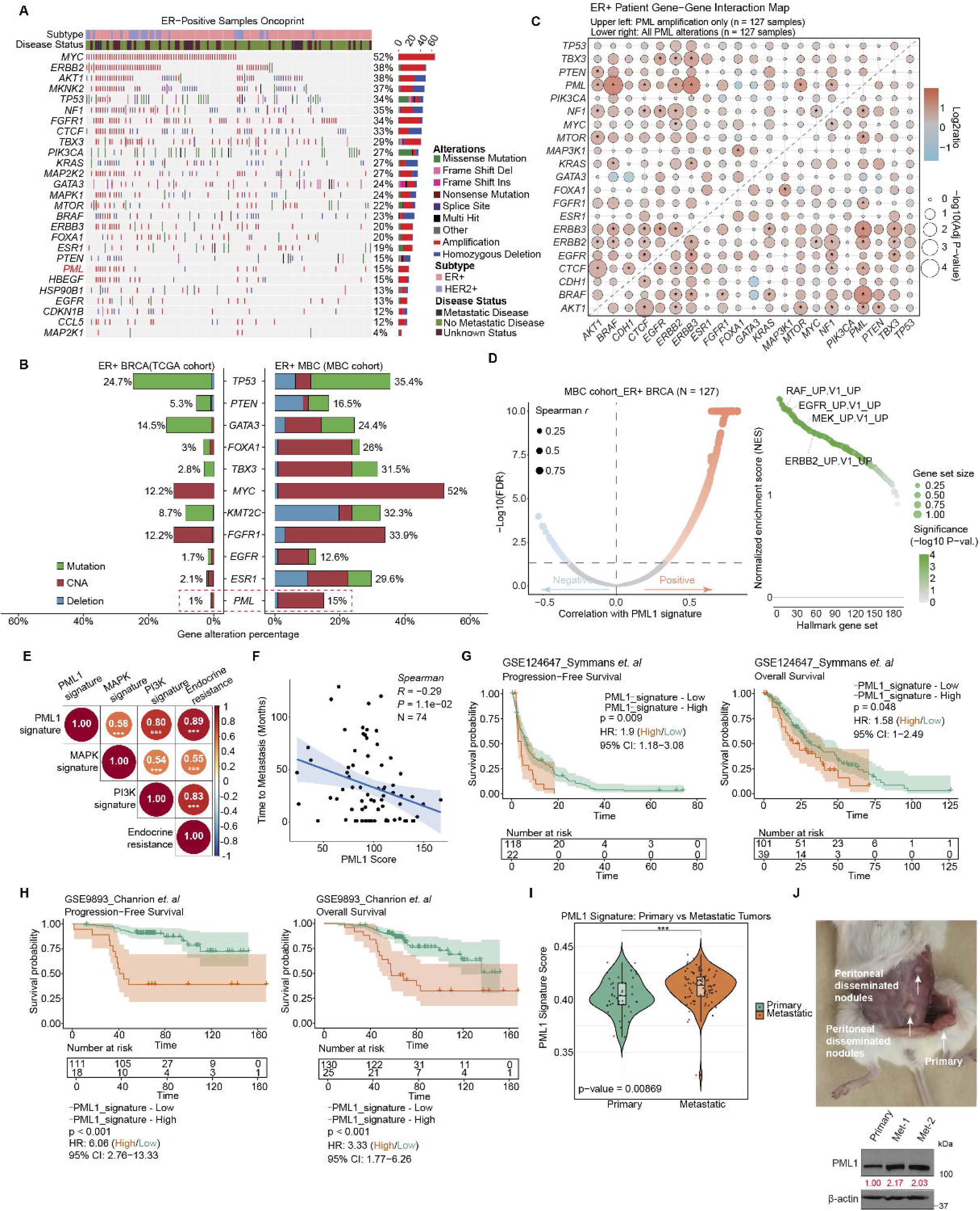
*PML* gene amplification and alterations in ER+ MBC. ***A,*** Oncoplots of mutations and copy number alterations (CNAs) showing *PML* gene amplification in ER+ MBC tumors (N=127). ***B,*** Comparative analysis of gene mutation and amplification frequencies between ER+ MBC and TCGA primary tumors reveals significantly higher rates of *PML* gene amplification in ER+ MBC. ***C,*** A gene-gene interaction map for ER+ MBC reveals significant co-occurrence between *PML* amplification/alterations and *AKT1*, *BRAF*, *ERBB2*, *ERBB3*, *MTOR*, and *NF1*. The asterisk indicates *p*-value < 0.05. ***D,*** *Left*, Volcano plot for a cohort of ER+ MBC patients. The x-axis shows correlation values between the PML1 gene signature and other mRNAs, and the y-axis shows statistical significance as - Log10(FDR). *Right*, Oncogenic gene set enrichment analyses show that the PML1 gene signature is most significantly enriched in RAF, EGFR, MEK, and ERBB2 signaling in ER+ MBC. ***E,*** Significant associations of PML1 gene signature with PI3K, MAPK, and endocrine resistance gene signatures in ER+ MBC. ***F,*** PML1 gene signature scores inversely correlate with time to metastasis in ER+ MBC patients. ***G-H,*** High PML1 gene signature scores correlate with progression-free survival and overall survival in two cohorts, GSE124647 (***G***) and GSE9893 (***H***). ***I,*** PML1 gene signature scores were significantly elevated in metastatic breast cancer (MBC) compared to primary tumors (n=42). ***J,*** Disseminated nodules derived from ^Y537S^ER1-KI cell xenografts exhibited significantly increased PML1 protein expression relative to primary tumor sites.

Analysis of TCGA data revealed significantly higher *PML* amplification frequency in ER+ and total MBC cases than in primary tumors (Fig. 5B & S14B), suggesting a potentially crucial role for the *PML* gene in metastatic progression. Genomic analyses demonstrated mutually exclusive patterns between *TP53* and *ESR1* gene mutations in total MBC cases (Fig. S14C), consistent with recent literature (44). Notably, *PML* gene alterations and amplifications significantly co-occurred with gene alterations in *AKT1*, *BRAF*, *CTCF*, *ERBB2*, *ERBB3*, *MTOR*, and *NF1* in ER+ and total MBC cases, suggesting functional interactions (Fig. 5C & S14C).

RNA-seq analyses of ER+ (Fig. 5D) and total MBC cases (S14D) revealed correlations between the PML1 gene signature and total mRNA expression. Gene set enrichment analyses revealed strong associations between the PML1 gene signature and RAF, EGFR, ERBB2, and MEK. We further validated the correlations between PML1 gene signature and PI3K, MAPK, and endocrine resistance gene signatures in ER+ and total MBC cases (Fig. 5E & S14E). Additionally, patients with elevated PML1 gene signatures exhibited reduced time to metastasis, progression-free survival, and overall survival across two MBC clinical datasets, as well as increased metastatic potential (Fig. 5F-I & S15A-D). In xenograft models, disseminated tumors harboring the Y537S mutation displayed increased PML1 protein expression relative to primary tumors (Fig. 5J), further supporting the relationship between metastatic progression and PML1 abundance.

### MEK/ERK and AKT/mTOR signaling enhance PML1 protein accumulation in ^Y537S^ER-KI cells and mediate PML1 protein induction by Fulv treatment

While Fulv significantly increased PML1 protein abundance, *PML1*, *PML2*, and *PML4* mRNA remained unchanged (Fig. S16A-B). Similarly, ^Y537S^ER- and ^D538G^ER-KI variants exhibited higher PML1 protein levels than ^WT^ER-KI cells without corresponding differences in *PML1*, *PML2*, or *PML4* mRNA levels. These observations suggest post-transcriptional regulation as the underlying mechanism that regulates PML1 protein abundance. To investigate this, we conducted cycloheximide chase experiments to assess protein stability across different cell variants and investigate this mechanism. ^Y537S^ER-KI cells exhibited a PML1 protein half-life exceeding 48 hours, compared to 24 hours in ^WT^ER-KI cells and slightly less than 24 hours in ^D538G^ER-KI MCF-7 (Fig. 6A). Similarly, we observed extended PML1 protein stability in our in-house Fulv-resistant cells (Fig. S16C) and in ^Y537S^ER-KI T47D cells (Fig. S16D).

**Figure 6.**
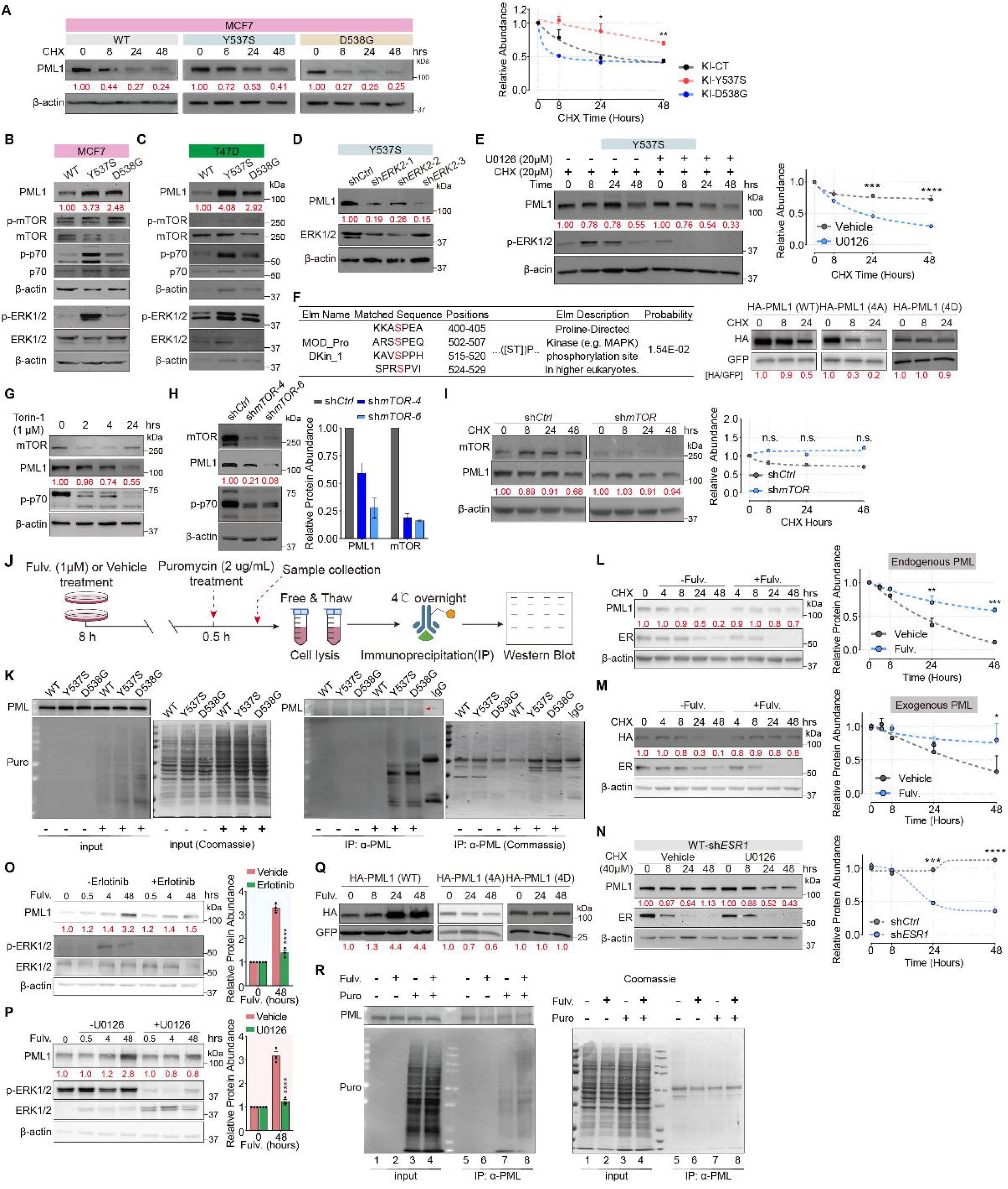
*ESR1* mutations and Fulv treatment regulate PML1 protein stability and synthesis. ***A,*** Protein stability assays demonstrate extended PML1 protein in ^Y537S^ER-KI cells versus ^D538G^ER- and ^WT^ER-KI cells. ***B-C,*** Western blots show enhanced mTOR and MEK pathway activation in ^Y537S^ER-KI MCF-7 (***B***) and T47D (***C***) cells compared to ^WT^ER-KI cells. ***D-E***, ***E-F,*** ERK pathway inhibition using shRNA-mediated knockdown (***D***) or MEK inhibitor U0126 (***E***) reduces PML1 protein levels and half-life in ^Y537S^ER-KI MCF-7 cells. ***F,*** *Left*, Putative ERK phosphorylation sites. *Right,* Half-life analysis shows that the PML1 (4D) variant has the most extended protein stability among PML1 variants. ***G-I,*** In ^Y537S^ER-KI cells, mTOR inhibition by Torin-1 treatment (***G***) or knockdown (***H***) decreases PML1 protein levels without affecting protein half-life (***I***). ***J*,** Schematic of the SUnSET protein synthesis analysis workflow. ***K,*** SUnSET assays reveal increased PML1 protein synthesis in ^Y537S^ER- and ^D538G^ER-KI cells versus ^WT^ER-KI cells. Cells were treated without (lanes 1-3) or with (lanes 4-6) puromycin (2 μg/ml, 30 min). Lysates were analyzed by Western blot using anti-PML or anti-Puro antibodies (left), with Coomassie staining (right). Cell lysates were immunoprecipitated with PML antibodies, followed by Western blots with anti-PML (top) and anti-Puromycin antibodies (bottom). Control antibody IP from ^Y537S^ER-KI cell lysates was a negative control (lane 7). ***L-N,*** Fulv (1 μM) extends the half-life of both endogenous PML1 in MCF-7 cells (***L***) and exogenous HA-tagged PML1 in stable-expressing cells **(*M*)**. ***N***, ERK inhibition significantly shortens *ESR1* knockdown-induced extended half-life. ***O-P*,** Treatments of MEK inhibitor U0126 (20 μM) (***O***) or EGFR inhibitor erlotinib (1 μM) (***P***) partially reverse Fulv-induced PML1 protein accumulation. ***Q,*** Western blots demonstrate Fulv (1 μM) increases wild-type PML1 levels, not PML1 (4A), or PML1 (4D) variants. ***R,*** Fulv treatment enhances PML1 protein synthesis in ^Y537S^ER-KI cells.

Given the established connection between PML1 protein levels and PI3K/MAPK pathways, we examined pathway activity in KI cells. ^Y537S^ER-KI MCF-7 and T47D cells showed significantly elevated MEK and Akt-mTOR activity (Fig. 6B-C). Analysis of independent RNA-seq datasets revealed an increase in PML1, PI3K, MAPK, and endocrine resistance gene signatures in response to Fulv treatment (Fig. S16E). While ^D538G^ER-KI cells displayed elevated mTOR activity like ^Y537S^ER-KI cells, their ERK activation was modestly higher than in ^WT^ER-KI cells. Investigation of ERK2’s role revealed that its depletion significantly reduced PML1 protein abundance in ^Y537S^ER-KI MCF-7 cells (Fig. 6D), and ERK2 inactivation by U0126 significantly decreased PML1 protein half-life (Fig. 6E). However, U0126 showed minimal effects on PML1 protein half-lives in ^WT^ER- or ^D538G^ER-KI MCF-7 cells (Fig. S16F).

Analysis of patient proteomic data (CPTAC) revealed positive correlations between PML protein abundance and phospho-peptides at S403, S505, S518, and S527/S530 (Fig. S17). Proteomics analyses of stably expressed FLAG-PML1 further validated these phospho-peptides in MCF-7 cells (Fig. S18). We thus hypothesized that activated ERK mediates PML1 phosphorylation at these residues, leading to protein stabilization and accumulation. To test, we generated PML1 mutants by substituting serine residues with either alanine (A) or aspartate (D) and assessed their half-lives. The PML1 (4D) phosphomimetic mutant showed the highest stability, while the PML1 (4A) phospho-deficient mutant exhibited the lowest stability (Fig. 6F).

We further established the role of the Akt-mTOR pathway in regulating PML1 protein abundance by employing pharmacological (Torin-1) and genetic (*MTOR* knockdown) approaches. Both interventions significantly reduced PML1 protein abundance in ^Y537S^ER-KI cells (Fig. 6G-H). However, *MTOR* knockdown did not decrease PML1’s half-life (Fig. 6I). Interestingly, Torin-1’s effects were cell variant-type specific: minor impact in ^WT^ER-KI cells but decreased PML1 protein levels in ^D538G^ER-KI cells (Fig. S19A-B), similar to its effects on PML1 in ^Y537S^ER-KI cells.

The observation that ^D538G^ER-KI cells maintain higher PML1 protein levels than ^WT^ER-KI cells despite similar mRNA levels and protein stability, combined with elevated mTOR activity in ER mutant cells and its induction by Fulv, led us to investigate potential differences in PML1 protein synthesis rates among these cell lines. Using the SUnSET (surface sensing of translation) technique (Fig. 6J), we quantified higher synthesis rates of selected proteins in ^Y537S^ER- and ^D538G^ER-KI cells compared to ^WT^ER-KI cells, as demonstrated by stronger puromycin-labeled peptide signals in ^Y537S^ER- and ^D538G^ER-KI whole cell lysates (Fig. 6K, input, lanes 5 & 6) versus ^WT^ER-KI cells (lane 4). Immunoprecipitation with anti-PML antibodies revealed more intense puromycin-labeled signals in lysates from ^Y537S^ER- and ^D538G^ER-KI cells (Fig. 6K, IP, lanes 5 and 6) compared to ^WT^ER-KI cells (lane 4), indicating higher synthesis rates of PML1 protein and/or its associated proteins in the mutant cells.

To elucidate the mechanism underlying Fulv-induced PML1 protein accumulation, we next examined the effects of Fulv treatment on protein stability and pathway involvement. Fulv treatment significantly extended half-lives of both endogenous and exogenous PML1 proteins: in parental MCF-7 and T47D cells, endogenous PML1 half-life increased from 24 hours to beyond 48 hours (Fig. 6L, S19C), while in HA-PML1 stably expressing cells, exogenous PML1 half-life extended beyond 48 hours (Fig. 6M). Moreover, treatment with U0126 significantly reduced *ESR1* knockdown-induced extended PML1 protein half-life (Fig. 6N).

The early Fulv-induced activation of ERK and p70 precedes the accumulation of PML1 protein (Fig. S11A). To investigate the connection between Fulv-induced PML1 accumulation and pathway activation, we examined the effects of MEK, mTOR, and EGFR inhibitors. Inhibition of MEK, EGFR, or mTOR activity caused a significant reduction in Fulv-induced PML1 protein accumulation (Fig. 6O-P & S19D-K). Additionally, treatments with U0126 partially dampened *ESR1* knockdown-induced PML1 protein accumulation in the ^Y537S^ER-KI cell variant (Fig. S19L). These results suggest that while EGFR- and MEK-dependent pathways contribute to Fulv-induced PML1 accumulation, additional mechanisms likely play important roles. Furthermore, Fulv increased exogenous PML1 (WT) protein abundance but not the 4A nor 4D mutants, suggesting that Fulv-induced ERK activation is critical for its PML1-elevation activity (Fig. 6Q). Additionally, Fulv treatment increased PML1 protein synthesis rates in ^Y537S^ER-KI cells (Fig. 6R). These findings collectively indicate that Fulv-induced PML1 protein accumulation is mediated through multiple mechanisms: extended protein half-life, reduced degradation, partial dependence on EGFR and MEK signaling pathways, and likely involvement of additional yet unidentified pathways.

#### PML1 Induces CCL5 And HB-EGF Expression And Secretion, Contributing To Fulv Resistance

Building on our previous work establishing PML1 as a transcriptional coactivator in enhancing breast cancer stem cell (BCSC) gene expression (28), we explored whether PML1 might upregulate growth factors and cytokines to activate receptor tyrosine kinases and induce PI3K/MAPK activation. We first examined the effects of conditioned medium (CM) from cells with varying PML1 expression levels on Fulv sensitivity. CM from PML1-overexpressing (OE) cells activated MEK and mTOR pathways (Fig. 7A) and promoted Fulv resistance in both ^WT^ER- and ^Y537S^ER-KI cells (Fig. 7B, Fig. S20A-B), suggesting PML1-induced secretion of resistance-promoting factors. CM from in-house Fulv-resistant cells, ^Y537S^ER- and ^D538G^ER-KI cells, which express higher PML1 protein levels, enhanced Fulv resistance (Fig. 7B & S20C-D).

**Figure 7.**
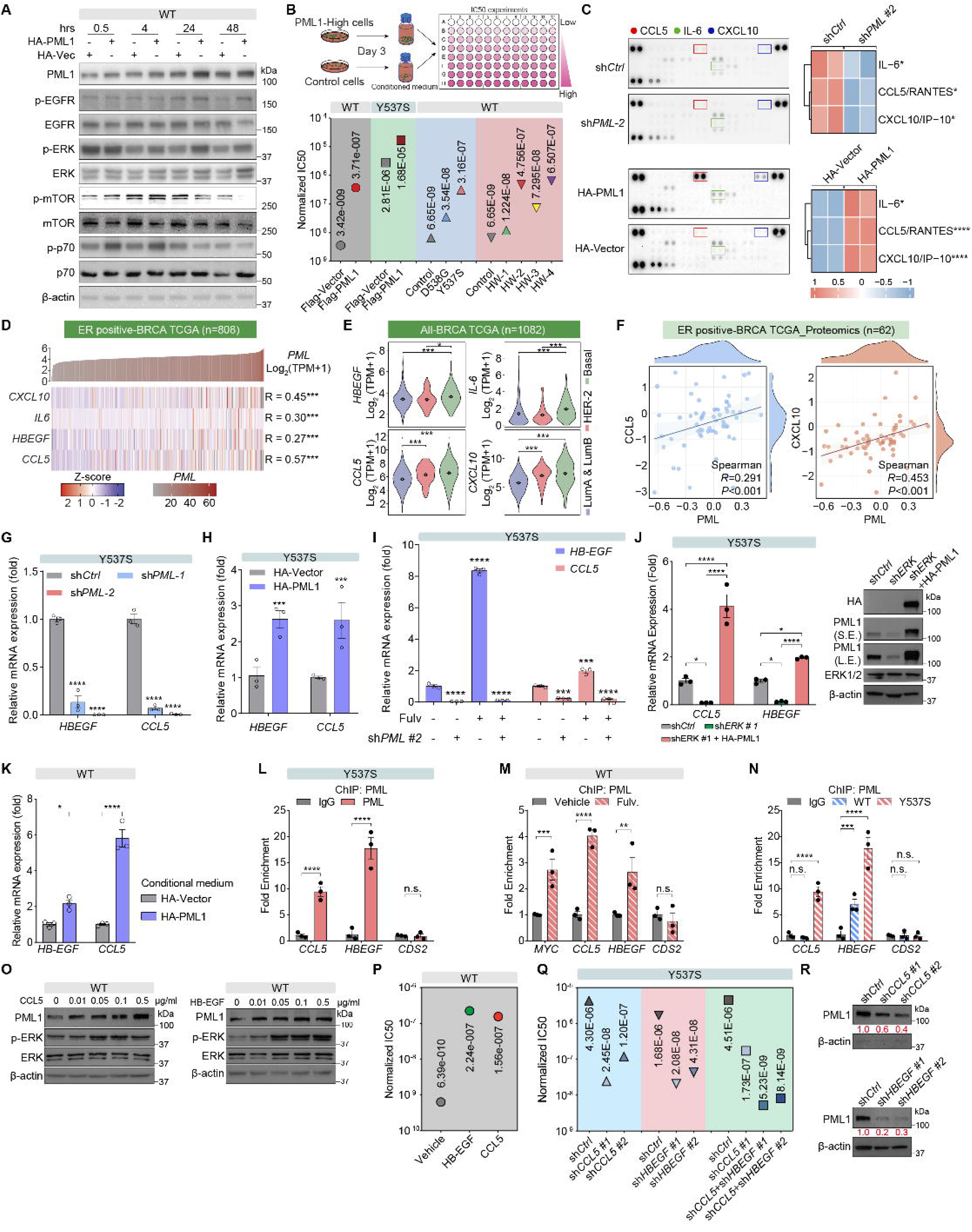
PML1 establishes a feed-forward loop through CCL5 and HB-EGF to activate PI3K and MAPK signaling pathways. ***A,*** Conditioned media (CM) from PML1-overexpressing cells, PML1 (OE), activates MEK-ERK and Akt-mTOR signaling pathways. ***B***, *Top*, a schematic of CM assays. *Bottom*, CM from PML1-overexpressing cells, PML1 (OE) confers Fulv resistance in both ^WT^ER- and ^Y537S^ER-KI MCF-7, and CM from ^Y537S^ER- and ^D538G^ER-KI and in-house Fulv-resistant MCF-7 cell variants and confer higher Fulv IC_50_ than CM from ^WT^ER-KI control cells. ***C,*** Cytokine array screening reveals PML1-dependent regulation of secreted CCL5, IL-6, and CXCL10 (left). Quantification of the secreted cytokine profile is presented as a heat map (right panel). ***D,*** RNA-seq analyses of ER+ breast tumor (TCGA) data demonstrate a positive correlation between *PML1* expression and *CXCL10*, *IL6*, *HBEGF*, and *CCL5* mRNA levels. ***E,*** Analyses ER+ breast tumor (TCGA) data reveal positive correlations between protein levels of PML and CXCL10 (*left*) and PML and CCL5 (*right*). ***F,*** Analyses ER+ breast tumor (TCGA) reveals significant elevation of CCL5 and CXCL10 cytokines in TNBC compared to luminal breast tumors. ***G-I***, Expression analysis demonstrates *CCL5* and *HBEGF* mRNA levels following *PML* knockdown in ^Y537S^ER-KI MCF-7 cells (***G***), effects of PML1 overexpression on *CCL5* and *HBEGF* expression in ^WT^ER-KI cells (***H***), and PML1-dependent induction of *CCL5* and *HB-EGF* expression in response to Fulv treatment (***I***). ***J,*** *Left*, PML1 rescues *CCL5* and *HBEGF* mRNA expression in ERK2 knockdown cells. *Right*, Western blot of the indicated proteins. ***K,*** CM of PML1-overexpressing cells induces *CCL5* and *HBEGF* mRNA levels. ***L-N,*** ChIP assays reveal PML binding to *CCL5* and *HBEGF* promoters in ^Y537S^ER-KI MCF-7 cells (***K***), increased PML recruitment to *CCL5* and *HBEGF* promoters following 24-hour Fulv treatment in ^WT^ER-KI cells (***L***), and increased PML occupancy at these promoters in ^Y537S^ER-KI cells compared to ^WT^ER-KI cells (***M***). The *CDS2* promoter serves as a negative control, showing no PML binding. ***O-P,*** Treatment with recombinant CCL5 or HB-EGF enhances PML1 protein accumulation (***O***) while inducing Fulv resistance in MCF-7 cells (***P***). ***Q-R,*** Knockdown of *CCL5* or *HBEGF* enhances sensitivity to Fulv (1 μM) (***Q***) and reduces PML1 protein levels in ^Y537S^ER-KI MCF-7 cells (***R***).

To identify PML1-induced secreted molecules, we conducted cytokine array analyses, which revealed significantly reduced CCL5 (RANTES) levels in CM from *PML*-knockdown ^WT^ER-(Fig. 7C, top panel & S20E) and ^Y537S^ER-KI (Fig. S20F) MCF-7 cells compared to controls, with this reduction reversed in CM from PML1 (OE) cells (Fig. 7C, bottom panel). We observed a consistent effect of CM from *PML* knockdown ^Y537S^ER-KI cells on *IL-6* and *CXCL10* levels (Fig. 7C, S20E-F).

Analyses of ER+ breast tumor RNA-seq data showed positive correlations between *PML* and *CCL5*, *HBEGF*, *IL6*, and *CXCL10* mRNAs in ER+ (Fig. 7D) and total breast tumors (Fig. S20G). Moreover, like the PML1 gene signature, *CCL5*, *HBEGF*, *IL6,* and *CXCL10* mRNA levels are significantly higher in TNBC than in luminal breast cancer (Fig. 7E). We also observed strong correlations between PML and CCL5 or CXCL10 at the protein levels in ER+ breast tumors (Fig. 7F). No information on IL-6 or HBEGF protein abundance is available. Mechanistic studies showed that *PML* knockdown reduced *CCL5* and *HB-EGF* mRNA levels in MCF-7 and T47D cells (Fig. 7G, Fig. S20H-I), while PML1 overexpression increased their mRNA levels (Fig. 7H). The expression levels of *CCL5*, *HBEGF*, *IL6,* and *CXCL10* mRNAs also correlate with the PML1 gene signature in ER+ and total MBC (Fig. S20J). Independent CPTAC proteomics data analysis revealed positive correlations between PML and CXCL10, as well as between PML and CCL5 (Fig. S20K). Additionally, significantly higher levels of CCL5 and CXCL10 were observed in TNBC compared to luminal breast cancer (Fig. S20L), further supporting the conclusions derived from TCGA data analysis. We showed that PML was essential for Fulv-induced *CCL5* and *HB-EGF* expression (Fig. 7I), and PML1 rescued *CCL5* and *HBEGF* mRNA expression in *ERK2* knockdown cells (Fig. 7J). Cells treated with CM derived from PML1-overexpressing cells express higher *CCL5* and *HBEGF* mRNAs than those treated with CM from control cells (Fig. 7K). ChIP-qPCR analysis demonstrated PML binding to *CCL5* and *HBEGF* promoters (Fig. 7L). Fulv treatment enhanced PML recruitment to these promoters (Fig. 7M & S20M), with ^Y537S^ER-KI cells showing more PML binding than ^WT^ER-KI cells (Fig. 7N).

In functional studies, recombinant CCL5 and HB-EGF enhanced PML1 protein levels and ERK activation (Fig. 7O) and promoted Fulv resistance (Fig. 7P). Conversely, *CCL5* or *HBEGF* knockdown enhanced Fulv’s anticancer activity in ^Y537S^ER-KI cells (Fig. 7Q) and reduced PML1 protein levels (Fig. 7R). These results indicate that elevated CCL5 or HB-EGF are necessary and sufficient to promote Fulv resistance. Moreover, *HBEGF* and *CCL5* genes are amplified in 12% and 9% ER+ MBCs, respectively (Fig. 5A).

These findings establish a novel feedback mechanism in which PML1 upregulates the expression of cytokines and growth factors, including *CCL5* and *HBEGF*, which trigger the PI3K and MAPK signaling pathways. This increases PML1 protein accumulation by enhancing protein stability and synthesis, thereby promoting Fulv resistance. This PML1-mediated secretion of pro-survival factors is a previously unrecognized mechanism driving endocrine therapy resistance in breast cancer.

## DISCUSSION

Our study reveals three significant findings that may inform novel strategies for overcoming endocrine therapy resistance: (1) PML1 plays a previously unrecognized role in MBC pathogenesis through a complex feedforward regulatory circuit with PI3K/MAPK signaling, (2) PML1 elevation paradoxically occurs in response to standard therapies, and (3) targeting PML1 with arsenic trioxide (ATO) represents a promising therapeutic strategy.

Through multi-omics analysis, functional studies, scRNA-seq, and spatial transcriptomics, we established PML1 as a master mediator of endocrine therapy resistance in ER+ breast cancer. The PML1 gene signature strongly correlates with reduced progression-free survival, overall survival, and time to metastasis, while substantially overlapping with gene signatures related to PI3K, MAPK, and endocrine resistance.

Targeted whole-exome sequencing accounts for only approximately 40% of metastatic breast cancer cases (24), highlighting the significant role that genetic alterations in other genes or non-coding sequences likely play in MBC progression. The MBC project aimed to identify whole-genome alterations, validate previous findings, and expand the catalog of genes potentially involved in MBC development.

PML has long been considered a tumor suppressor (45,46). However, emerging evidence supports PML as an oncogene, particularly in breast cancer contexts (47–50). The significant gene amplification of *PML*, *CCL5*, and *HBEGF* observed in MBC cases reported by the MBC project and their role in enhancing PI3K and MAPK signaling pathway activity represent potentially targetable alterations that could inform novel therapeutic approaches for metastatic disease.

We uncovered a self-reinforcing regulatory network where genetic alterations such as *PML* gene amplification activate both PI3K and MAPK signaling pathways through enhanced expression of cytokines and growth factors, while hyperactivated PI3K/MAPK pathways (resulting from mutations, amplifications, or ER mutations such as Y537S and D538G) increase PML1 protein stability and synthesis. This feedforward loop contributes to *de novo* and acquired therapeutic resistance, providing a unifying framework for diverse resistance mechanisms.

Our analysis of ER variants reveals distinct mechanisms regulating PML1 protein levels: ^Y537S^ER-KI cells showed enhanced PML1 protein stability, accelerated synthesis, and the highest protein abundance; ^D538G^ER-KI cells exhibited elevated synthesis but lower turnover and intermediate abundance; and ^WT^ER-KI cells demonstrated the lowest synthesis rates and protein abundance. These differences elucidate how ER mutations may drive resistance through distinct effects on PML1 protein homeostasis.

A critical insight is the inverse relationship between ER and PML1: ER inhibition (through SERD treatment or knockdown) triggers PML1 accumulation. This relationship explains why standard treatments may paradoxically promote resistance. The effects of Fulv treatment are context-dependent: in ^WT^ER-KI cells, anti-tumor effects of ER degradation dominate despite increased signaling, inducing cell death; however, in cells with PML amplification, hyperactive signaling pathways, or *ESR1* mutations, Fulv further enhances these pathways while increasing PML1, driving resistance.

PML1’s influence extends beyond PI3K/MAPK signaling to regulate immune modulators (CCL5, IL6, CXCL10), enabling tumor modulation through endocrine and paracrine mechanisms. The observation that CDK4/6 inhibitor treatments induce PML1 accumulation aligns with the established complex crosstalk between CDK4/6 and PI3K/MAPK signaling pathways (51–54). This interaction is particularly significant in clinical settings, as the combined use of CDK4/6 inhibitors and Fulv therapy represents the current standard of care for ER+ MBC patients. The widespread elevation of PML1 protein levels in CDK4/6i-resistant ER+ breast tumors suggests its potential dual role as both a resistance mediator and a predictive biomarker. This finding has immediate translational relevance for patient care and treatment planning. Furthermore, as CDK4/6 inhibitor therapy expands into clinical trials across diverse cancer types, therapeutic strategies targeting the PML1-PI3K/MAPK axis may provide a novel approach to prevent or overcome drug resistance, potentially extending the clinical benefit of these critical targeted therapies.

Our findings reveal promising therapeutic strategies through two key observations: (1) arsenic trioxide effectively reduces PML1 protein levels and synergistically enhances Fulv efficacy in wild-type and ER-mutant tumors, and (2) PML1 reduction simultaneously suppresses both PI3K and MAPK signaling pathways. With ATO’s established safety profile in leukemia treatment, clinical trials of Fulv/ATO combinations can be expedited, particularly for patients with elevated PML1 levels.

While establishing PML1 as a central mediator of therapy resistance, several questions remain: the precise mechanisms by which ER inhibition triggers rapid PI3K/MAPK activation and PML1 accumulation; PML1’s complex interactions with immune cells; optimal dosing strategies for ATO/Fulv therapy; identification of biomarkers for patient stratification; and whether early intervention targeting the PML1-PI3K/MAPK axis can prevent CDK4/6i resistance.

## Methods

### Establishment and maintenance of drug-resistant cell lines

Drug-resistant cell lines were established through continuous exposure to increasing concentrations of various compounds. For fulvestrant resistance, we maintained cells in concentrations ranging from 200, 500, 800, and 2000 nM. Elacestrant resistance was developed using 500 nM and 1000 nM concentrations, while CDK4/6 inhibitor resistance (using palbociclib and abemaciclib) was established with 1000 nM treatment.

The resistance development process involved long-term maintenance in drug-containing media, during which initial growth inhibition and cell death were observed. Culture media was regularly refreshed while monitoring cell survival and proliferation. Cell capable of growing under high drug concentrations were selected over several months. After establishment, drug resistance was confirmed using CCK8 assays to determine IC50 values, with parental cells serving as controls. To maintain the stability of the resistant phenotype, passage numbers were limited (within 15 passages), and cells were continuously cultured in media containing maintenance doses of their respective drugs (fulvestrant, elacestrant, or CDK4/6 inhibitors) to preserve resistance characteristics.

### Transient Transfection

Cells in the logarithmic growth phase were seeded in 6-cm dishes. 8 μl Lipofectamine 2000 was diluted in 400 μl Opti-MEM and incubated at room temperature for 5 minutes. The plasmid DNA (4 μg) was then added to the Opti-MEM-Lipofectamine 2000 mixture, gently mixed, and incubated at room temperature for 20 minutes. The mixture was subsequently added to the cells.

### Lentivirus Production and Transduction

HEK293T cells within 10 passages, maintained in the logarithmic growth phase, were seeded in 10-cm culture dishes. After adhering to the plate, cells were transfected using Lipofectamine 2000 with packaging plasmids at the following ratios: target plasmid:PAX2:PMD2.G (5:1.25:3.75 μg). The cells were washed with PBS four hours post-transfection and refreshed with a complete medium. Viral supernatants were collected 48-, 72-, and 96-hours post-transfection and concentrated using Vivaspin concentrators (15R-maximum spin speed, Sartorius).

A reverse transduction method was employed for viral transduction. Target cells were trypsinized, neutralized with a complete medium, and centrifuged at 1,000 rpm for 5 minutes. A transduction mixture containing the virus, polybrene (at a final concentration of 10 μg/ml), and a complete medium was prepared. Cells were resuspended in this mixture and plated. After 24 hours, the medium was refreshed, and stable transductants were selected using puromycin (2 μg/ml).

### Whole cell lysate preparation and Western blotting

Cells were lysed using RIPA buffer (50 mM Tris pH 7.5, 150 mM NaCl, 1% NP-40, 0.5% Deoxycholic acid, 0.1% SDS). For 6cm dishes with 80% cell confluence, approximately 200 μl of RIPA buffer was used, while 10cm dishes with 80% confluence required approximately 600 μl. Lysis was performed on ice, with vortexing every 2 minutes and mixing using a 1000 μl pipette to ensure complete homogenization. After 15 minutes, samples were prepared immediately for western blot analysis or stored at -80°C. When retrieving samples from -80°C storage, samples were resuspended using a 1000 μl pipette to ensure thorough mixing before sample preparation for Western blottings. Protein samples were separated on 8% or 10% SDS-PAGE and transferred onto PVDF membranes (Bio-Rad, #1620177). Membranes were blocked with phosphatase-free, non-fat milk (LabScientific, #M-0842) for 1 hour at room temperature to minimize non-specific binding, followed by overnight incubation with primary antibodies at 4°C. The following day, membranes were incubated with corresponding horseradish peroxidase (HRP)-conjugated secondary antibodies for 1 hour at room temperature. After washing three times with 1x PBST buffer, chemiluminescent signals were detected with the ChemiDoc MP imaging system (Bio-Rad, #12003154).

### Cell Viability and Drug Sensitivity Analysis

Cell Growth Assay: 2,000 cells in the logarithmic growth phase were seeded into 96-well plates (Fisherbrand™, #FB012931). Twelve hours later, 10 µl of Cell Counting Kit-8 (CCK-8, Dojindo, #CK04-05) was added to each well, followed by 2 hours of incubation at 37°C. The absorbance at 450 nm was measured using a SpectraMax® M2 Multimode microplate reader (Molecular Devices, #89429-532) and recorded as the baseline value (Day 0). Cell ODs were recorded on Days 3, 5, and 7. Results are presented as Mean ± SEM from four technical replicates with three biological repeats.

### IC50 Determination

Cells were seeded as described above, allowed to attach for 24 hours, and treated with serial dilutions of the test compound. On Day 7, cell viability was assayed using the CCK-8 assay described above. The normalized IC50 values were calculated using GraphPad PRISM 9 software, fitting the data to a "log(inhibitor) vs. normalized response - Variable slope" algorithm.

### Colony Formation Assay

Five hundred cells in the logarithmic growth phase were seeded into 24-well plates, attached for 12 hours, and treated with the indicated compounds. Following ten days of incubation, the medium was removed, and cells were fixed with 4% paraformaldehyde at room temperature for 15 minutes. After washing with PBS to remove residual paraformaldehyde, colonies were stained with 0.2% crystal violet solution (Millipore Sigma, #C0775) at room temperature for 15 minutes. The plates were then gently washed with PBS to remove background staining. Colony counting was performed using ImageJ software, where colonies were defined as cell clusters containing 50 or more cells. Results are presented as Mean ± SEM base on two biological repeats, each with three technical replicates.

### 3D Spheroid Formation Assays

For 3D spheroid culture, 1,500 cells from 2D monolayer culture in the logarithmic growth phase were seeded into each well of CELLSTAR® 96-well Cell-Repellent U-bottom microplates (Greiner Bio-One, #07-000-612) in 200 µl culture medium. Plates were centrifuged at 1,000 RPM for 5 minutes to facilitate cell aggregation at the bottom of the wells. After 2-3 days of incubation, when the spheroids had formed, initial images were captured using a microscope (Eclipse Ts2R, Nikon) and designated as Day 1. For drug treatments, the medium was removed and replaced with 100 µl of fresh medium containing either test compounds or vehicle control. After each medium change, plates were centrifuged at 1,000 RPM for 5 minutes to maintain spheroid integrity. Spheroids were cultured for a total of 17 days. Spheroid volumes were calculated using ImageJ software, and growth curves were generated and analyzed using GraphPad PRISM software.

### Orthotopic xenograft

The animal experiments were conducted according to protocols approved by the Institutional Animal Care and Use Committee (IACUC) of Case Western Reserve University. Female NOD-SCID mice were maintained under pathogen-free conditions at the Athymic Animal Core Facility, Case Comprehensive Cancer Center. At approximately 6 weeks of age, mice underwent surgery for orthotopic tumor cell injection into mammary fat pads. MCF7-WT and Y537S cells (1×10^6^) were suspended in PBS, mixed with Matrigel (R&D Systems, #343300501) at a 1:1 ratio, and thoroughly homogenized. A vertical incision was made in the skin adjacent to the fourth pair of mammary glands, avoiding damage to the peritoneum. The mammary fat pad was carefully exposed through gentle tissue dissection. Forceps were used to lift the fat pad, and 100 μl of cell-Matrigel mixture was injected into each fat pad using a 25G needle. The fat pad was carefully repositioned to prevent cell leakage, and the procedure was repeated on the contralateral side. A 17β-estradiol pellet (1.7 mg, 60-day release; Innovative Research of America, #SE-121) was subcutaneously implanted.

Tumor dimensions were measured using digital calipers, and tumor volume was calculated using the formula V=W^2^×L/2, where W represents the width and L represents the length. When tumor volumes reached approximately 62.5 mm3, mice were randomized into four treatment groups: 1) Vehicle control; 2) Fulv (50 mg/kg, subcutaneous injection, twice weekly); 3) ATO (8 mg/kg, intraperitoneal injection, three times weekly); and 4) Combination therapy (Fulv 25 mg/kg, twice weekly; ATO 4 mg/kg, intraperitoneal injection, three times weekly). After 30 days of treatment, all mice were humanely euthanized, and tumors were harvested for subsequent analysis.

### Primary tumor cell extraction

Tumor tissue specimens were mechanically dissociated into 3-4 mm fragments using sterilized surgical scissors and rinsed with Hank’s Balanced Salt Solution (HBSS, Gibco™). Enzymatic digestion was performed by incubating tissue fragments in Collagenase Type I solution (200 U/ml in HBSS supplemented with 3 mM CaCl_, STEMCELL™ TECHNOLOGIES, #07902) at 37°C for 4 hours with gentle agitation. The resulting cell suspension was filtered through a sterile nylon mesh to separate single cells from undigested tissue fragments and collected in 50l conical tubes. The filtered cells were washed three times with HBSS via centrifugation at 1000 rpm for 5 minutes per wash. The cell pellet was resuspended in a complete growth medium following the final centrifugation and seeded onto 10 cm culture dishes for subsequent expansion and experimentation.

### *PML1*-specific knockdown shRNA construction

*PML1*-specific sequences corresponding to amino acids 620-882 were based on the reference sequence NM_033238.3 (Homo sapiens PML nuclear body scaffold (PML), transcript variant 1, mRNA) from NCBI. To design shRNAs targeting PML1-specific regions, we employed BLOCK-iT™ RNAi Designer (Thermo Fisher Scientific). We selected shRNA sequences with at least 3 nucleotide mismatches to unrelated genes to minimize off-target effects. Additional nucleotides were incorporated to form short hairpin structures and to introduce EcoRI and AgeI restriction sites, following the pLKO.1-TRC cloning vector protocol (Addgene, #10878).

For shRNA annealing, sense and antisense oligonucleotides were diluted to 20 μM and combined in an annealing reaction containing 2 μl each of sense and antisense oligos, 2 μl of 10× NEB buffer 2, and 14 μl of sterile ultrapure water. Annealing was performed using a T100 Thermal Cycler (Bio-Rad Laboratories Inc., USA) with the following temperature gradient: 5 min at 95°C, cooling to 70°C, followed by stepwise cooling at 2°C decrements for 5 min each until reaching room temperature.

The pLKO.1-TRC cloning vector was digested with EcoRI and AgeI restriction enzymes (NEB, #R3101; #R3552). With overnight incubation, annealed shRNAs were ligated into the digested vector using T4 DNA ligase (NEB). The transformation was performed using NEB® 5-alpha Competent E. coli (High Efficiency). After plasmid purification and sequence verification, lentiviral particles were produced in HEK293T cells and used to transduce breast cancer cell lines.

### RNA extraction and quantitative RT-PCR

Total RNA was extracted using the RNeasy Mini Kit (Qiagen, #74104) following the manufacturer’s instructions. One microgram (1,000 ng) of total RNA was reverse-transcribed using iScript supermix (Bio-Rad, #1708841). Quantitative PCR was performed on a Bio-Rad CFX Connect Real-Time PCR Detection System using iQ SYBR Green supermix (Bio-Rad, #1708882). We designed primers using NCBI Primer-BLAST with an exon-exon junction spanning strategy to avoid genomic DNA contamination. Primer sequences are listed in Table S3. Results are presented as Mean ± SEM from three technical replicates with two biological repeats.

### Primer Design for Quantitative PCR

Primers for qPCR were designed using NCBI’s Primer-BLAST tool (https://www.ncbi.nlm.nih.gov/tools/primer-blast/). The target gene was identified in the NCBI Gene database by searching for the gene name and selecting the entry for the appropriate species (Homo sapiens). Using the "Pick Primers" function within the Nucleotide record, parameters were set as follows: PCR product size range of 80-200 bp, and primers were required to span an exon-exon junction to prevent amplification of genomic DNA. From the resulting primer pairs, we selected those with GC content closest to 50% for both forward and reverse primers to ensure optimal annealing properties. Prior to qPCR experiments, all primers were validated by conventional PCR using Taq DNA polymerase (ThermoFisher Scientific), followed by electrophoresis on a 1% agarose gel to confirm the expected amplicon size, assess PCR efficiency (by comparison with established primer sets), and verify the specificity of amplification by the presence of a single band.

### SUnSET (surface sensing of translation)

Logarithmically growing cells were seeded in 10 cm culture dishes to achieve approximately 80% confluence. After 12 hours of attachment, cells were treated with 1 μM Fulvestrant or DMSO vehicle control. At 7.5 hours post-treatment, cells were exposed to 2 μg/mL puromycin or water control for 30 minutes. Cells were then lysed with 300 μl of S150 lysis buffer (50 mM Tris pH 7.5, 150 mM NaCl, 1 mM EDTA, 0.5% NP-40, 1X Protease and Phosphatase Inhibitor Cocktail (Roche, #11697498001; #4906845001), 125 U/mL Benzonase (Sigma-Aldrich, #E1014-5KU)). Cell lysis was enhanced using a freeze-thaw method: samples were flash-frozen on dry ice for 5 minutes, followed by thawing at 37°C for 1 minute with agitation. Lysates were centrifuged at 13,000 g for 30 minutes at 4°C, and supernatants were collected. A 20 μl aliquot was reserved as the IP input sample.

For immunoprecipitation, Pierce™ Protein G Agarose beads (Thermo Scientific™, #20397) were pre-incubated with 5 μg of anti-PML (rabbit) antibody or rabbit IgG control for 2 hours at 4°C. After washing the beads with S150 buffer to remove unbound antibodies, 20 μg of protein lysate was added, and the total volume was adjusted to 1,000 μl with S150 buffer. The samples were incubated overnight at 4°C with rotation. The next day, beads were collected by centrifugation at 3,000 RPM for 5 minutes and washed six times with S150 buffer. Immunoprecipitated proteins were eluted by adding 15 μl of water and 3 μl of 6X sample buffer.

Samples were analyzed by Western blot using anti-puromycin and anti-PML1 antibodies, with Coomassie blue staining performed to verify equal protein loading.

### Chromatin immunoprecipitation

Approximately 20 million cells were fixed with 1% formaldehyde (final concentration) and quenched with glycine (final concentration 125 mM). Nuclei were isolated using a fractionation method with differential lysis buffers. Nuclear lysis was performed using nuclei buffer containing 10 mM Tris-HCl (pH 8.0), 1 mM EDTA (pH 8.0), 0.5 mM EGTA, 100 mM NaCl, 0.1% Na-deoxycholate, and 0.5% Sarkosyl.

Chromatin was sheared using a Covaris E220 ultrasonicator with the following parameters: 140 peak intensity power, 10% duty factor, 200 cycles per burst, for 300 seconds per sample in AFA tubes (Covaris, #520135). After centrifugation at maximum speed, supernatants were collected, and 50 μl was reserved as input. Equal amounts of chromatin were used for immunoprecipitation (IP).

Chromatin samples were incubated with 5 μg of antibody overnight at 4°C. The following day, antibody-chromatin complexes were incubated with 50 μl Protein A/G PLUS-Agarose beads (Santa Cruz Biotechnology, sc-2003) for 2 hours at 4°C. Complexes were washed five times with ChIP wash buffer ( 50mM HEPES pH 7.5, 1mM EDTA, 0.7% Na-deoxycholate, 1% NP-40, 500 mM LiCl) followed by a final wash with modified TE buffer (10 mM Tris-HCl pH 8.0, 10 mM EDTA pH 8.0, 50 mM NaCl).

Chromatin was eluted using elution buffer containing 1× Proteinase K (elution buffer composition: 20mM Tris pH 8.0, 50mM NaCl, 5mM EDTA, 1%SDS). IP and input samples were incubated overnight at 65°C with shaking at 1300 rpm in a thermomixer. DNA was purified using a Qiagen PCR purification kit, and DNA concentration was determined using an Invitrogen Qubit 4 fluorometer. Primer sequences for qPCR are listed in Table S3.

### Immunofluorescence microscopy (IF)

Cell lines in the logarithmic growth phase were seeded onto sterile glass coverslips in 12-well plates, with 1 ml DMEM supplemented with 10% FBS or 1 ml RPMI supplemented with 10% FBS with 0.2 units/ml bovine insulin (Gibco #12585-014) per well, in triplicate. At 48 hours post-seeding, cells were treated with either DMSO (vehicle control) or 1 μM fulvestrant (Fulv). Following treatment, cells were washed with ice-cold PBS, fixed with 4% paraformaldehyde for 10 minutes, and permeabilized with 4% paraformaldehyde containing 0.1% Triton X-100 for 10 minutes. After six PBS washes, samples were blocked with freshly prepared 3% BSA for 1 hour, then incubated with primary antibodies against PML (#sc-966, Santa Cruz Biotechnology) and ER (#ab16660, Abcam) overnight at 4°C. Cells were washed six times with PBS-T buffer before incubation with secondary antibodies (Alexa Fluor 594 and Alexa Fluor 488, 1:1000 dilution in PBS, Invitrogen, #A32731) for 1 hour, followed by six washes with PBS-T. Coverslips were mounted onto microscope slides using a DAPI-containing mounting medium (Vectashield, #H-2000). Images were acquired using an Olympus BX-43 immunofluorescence microscope with a 40× objective, maintaining consistent imaging parameters across all samples.

### Immunofluorescence Image Quantification

Individual channels of each image were converted to 8-bit format. Regions of interest (ROIs) were selected under identical settings for each cell, and fluorescence intensity was measured. The analysis was performed on 100 cells per condition, and intensities were recorded and quantified.

### LC-MS and data analysis

Samples were analyzed by LC-MS using a Fusion Lumos Tribrid MS (ThermoScientific) equipped with a Dionex Ultimate 3000 nano UHPLC system, and a Dionex (25 cm x 75 µm id) Acclaim Pepmap C18, 2-μm, 100-Å reversed-phase capillary chromatography column. Peptide digests (5 μl) were injected onto the reverse phase column and eluted at a flow rate of 0.3 μl/min using mobile phase A (0.1% formic acid in H2O) and B (0.1% formic acid in acetonitrile). The gradient was held at 2% B for 5 minutes, %B was increased linearly to 35% in 105 minutes, increased linearly to 90% B in 10 minutes, and maintained at 90% B for 5 minutes. The mass spectrometer was operated in a data-dependent manner which involved full scan MS1 (350-1500 Da) acquisition in the Orbitrap MS at a resolution of 120000. This was followed by CID (1.6 Da isolation window) at 35% CE and ion trap detection. MS/MS spectra were acquired for 3 seconds. Dynamic exclusion was enabled where ions within 10 ppm were excluded for 60 seconds.

The raw data were analyzed by using all CID spectra collected in the experiment to search the human SwissProtKB database (downloaded on 3-23-2022, 26576 entries) and more specifically against the sequences of PML1 (P29590) and PML4 (P29590-5) with the program Sequest which is integrated in the Thermo Proteome Discoverer (V2.5) software package. Peptide and protein validation was performed using the percolator node with protein, peptide, and PSM thresholds at <1% FDR. The relative abundance of the positively identified proteins was determined using the extracted ion intensities (Minora Feature Detection node) with Retention time alignment. Only unique peptides were included in the quantitation.

### TCGA cohort analysis

#### TCGA-Transcriptomics

For transcriptomic analysis, we obtained TCGA Breast Cancer (BRCA) RNA-seq data (STAR-TPM) from the UCSC Xena GDC Hub along with TOIL RSEM isoform percentage data from UCSC Toil RNA-seq Recompute (55). We merged the two datasets based on patient identifiers to enable integrated analysis. PML isoform expression was quantified by calculating the percentage of transcripts containing exon 9, representing the PML1 isoform proportion. Correlation analyses between PML and other genes were performed using the Spearman rank correlation test. The correlation analysis was implemented using the cor.test function in R. For multiple testing corrections, we applied the Benjamini-Hochberg procedure (BH) using the p.adjust function with method="BH" to control the false discovery rate. Genes significantly correlated with PML (|r| > 0.25, adjusted p-value < 0.05) were identified and clustered using the mfuzz method to establish expression patterns. Pathway enrichment analysis of correlated genes was performed using the KEGG database through the ClusterGVis package.

#### TCGA-Proteomics

TCGA Breast Cancer (BRCA) proteomics data were obtained from the cBioPortal database for proteomic analysis. The dataset was filtered to remove proteins with more than 50% missing values across samples. Sample subtyping was performed based on the clinical annotation data, classifying samples into Luminal (LumA and LumB), HER2-enriched, and Triple-Negative Breast Cancer (TNBC) subtypes.

Similar to the transcriptomic analysis, correlation analyses between PML and other proteins were performed using Spearman’s correlation test, with Benjamini-Hochberg correction applied for multiple tests. Proteins significantly correlated with PML (|r| > 0.25, adjusted p-value < 0.05) were identified. We performed a correlation analysis for the luminal subtype to identify PML-associated proteins in hormone receptor-positive breast cancers. The significantly correlated proteins were visualized using hierarchical clustering and the ClusterGVis package, with samples ordered according to PML expression levels. The mfuzz clustering algorithm was used to identify distinct expression patterns among the correlated proteins, and we performed KEGG pathway enrichment analysis to identify biological processes associated with each cluster.

### Chromatin immunoprecipitation followed by sequencing (ChIP-seq) analysis

PML ChIP-seq datasets from three different cell lines (MCF-7, GM12878, and K562) were obtained from Gene Expression Omnibus (GEO) (GSM1010838, GSM1010771, and GSM1010722, respectively) (56,57) with corresponding Sequence Read Archive (SRA) accession numbers SRX190294, SRX190227, and SRX190178. All samples were generated using the same antibody (SC-71910).

Raw reads were assessed for quality using FastQC (v0.11.9), and adapter sequences were trimmed using Trim Galore (v0.6.7) with Cutadapt (v3.4). Filtered reads were aligned to the human reference genome using Bowtie2 (v2.4.4). Duplicate reads were marked using Picard MarkDuplicates (v2.27.4-SNAPSHOT), and mapping quality was evaluated with Samtools (v1.15.1). Peak calling was performed using MACS2 (v2.2.7.1) with a q-value threshold of 0.01, keeping only one tag at each location (--keep-dup 1) and an extension size of 146 bp (--extsize=146) without model building (--nomodel). Signal tracks for visualization were generated using BEDTools (v2.30.0) and converted to bigWig format using UCSC utilities (v377). Peaks were annotated with HOMER (v4.11) to identify their genomic distribution across promoters, exons, introns, and intergenic regions based on RefSeq gene annotations. Quality control metrics were calculated, including FriP (Fraction of Reads in Peaks), PBC (PCR Bottleneck Coefficient), and the overlap of peaks with DNase hypersensitive sites. DeepTools (v3.5.1) was used to generate heatmaps and profile plots, illustrating PML binding patterns relative to genomic features. Genes involved in pathways related to EGFR/ERBB2/RAF/MEK/ERK and PI3K/AKT/mTOR signaling were extracted from MSigDB (v2024.1), and the corresponding genomic regions were visualized using DeepTools to assess PML binding enrichment across these oncogenic signaling targets. We identified 10-fold- and 20-fold confident peaks for peak quality assessment based on MACS2 fold-change values. Consensus peaks across cell lines were determined using the MACS2 consensus module. Motif analysis and feature counts were performed to identify PML-associated DNA binding motifs and quantify reads over genomic features using SUBREAD featureCounts (v2.0.1). The resulting MACS2 peak files were further annotated using the R package ChIPseeker (v1.42.1) to characterize their genomic context. ChIPseeker’s annotatePeak function was used to assign peaks to genomic features, including promoters, untranslated regions (UTRs), exons, introns, and intergenic regions. Promoter regions were defined as sequences within 3kb upstream and 3kb downstream of transcription start sites.

### Activepathway analysis

To integrate multiple layers of genomic evidence and identify significantly enriched pathways, we employed the ActivePathways framework (version 2.0.5), an integrative multi-omics pathway enrichment method (58) (https://github.com/reimandlab/ActivePathways). This approach allowed us to combine ChIP-seq binding data with transcriptomic and proteomic correlation analyses to identify significantly enriched biological pathways associated with PML and PML1 signature expression patterns. Three types of data were integrated into our analysis: ChIP-seq peaks data from MCF7 cells with peaks annotated to identify promoter-associated binding events; PML protein expression correlation data from TCGA ER-positive breast cancer samples containing correlation coefficients and p-values between PML expression and genome-wide protein levels; and PML1 mRNA correlation data representing correlation coefficients and p-values between PML1 mRNA expression data and genome-wide gene expression in TCGA ER-positive breast cancer samples. We performed quality control steps for each dataset, including false discovery rate (FDR) correction using the Benjamini-Hochberg method. The datasets were filtered based on statistical significance thresholds (p-value < 0.05, FDR < 0.1) and correlation strength thresholds (|r| > 0.25) to focus on biologically meaningful associations.

We applied the directional p-value merging (DPM) method implemented in ActivePathways 2.0, which considers both statistical significance and the direction of effects. The analysis was conducted separately for positive and negative correlations to distinguish between activated and repressed pathways. The preprocessing workflow involved converting raw scores to statistical significance metrics, identifying gene-level summaries for each data type, segmenting correlations into positive and negative directions, creating unified p-value matrices with genes as rows and evidence sources as columns, and preparing corresponding direction matrices to encode the directionality of associations. For pathway annotations, we utilized the Reactome pathway database from MSigDB collections. Pathway size filters (5-1000 genes) were applied to focus on biologically interpretable gene sets. This configuration ensured pathways were only considered significant if they demonstrated coherent signals in both transcriptomic correlation datasets while accounting for potential ChIP-seq evidence.

The ActivePathways output was prepared for network visualization in Cytoscape using the EnrichmentMap plugin. The analysis automatically generated the necessary network files (nodes, edges, and enrichment results) with specific prefixes for positively and negatively correlated pathways. In Cytoscape, we imported these files using the EnrichmentMap App, an edge cutoff of 0.6 to connect pathways with substantial gene overlap, and the combined constant calculation method. The resulting pathway networks were visualized with node size proportional to gene set size and edge thickness representing the overlap between pathway gene sets. The AutoAnnotate plugin was applied to generate descriptive labels for pathway clusters based on word frequency in pathway names. This visualization approach allowed us to identify coordinated biological processes regulated by PML1 across multiple data types. The entire analytical workflow allowed for the systematic exploration of pathway enrichment patterns by integrating epigenomic binding data and transcriptomic and proteomic correlation profiles, providing a comprehensive view of PML-regulated biological processes in ER-positive breast cancer.

### scRNA-seq analysis

#### Tamoxifen-treated recurrent scRNA-seq dataset analysis

Single-cell RNA sequencing (scRNA-seq) data from GSE240112 (41) were analyzed to investigate transcriptional changes across normal breast tissue (NT), primary breast tumors (PT), and tamoxifen-treated recurrent tumors (RT). The dataset comprised eight samples: two normal tissue samples (GSM7681685_NT7, GSM7681686_NT8), three primary tumor samples (GSM7681687_PT1, GSM7681688_PT2, GSM7681689_PT5), and three tamoxifen-treated recurrent tumor samples (GSM7681690_RT3, GSM7681691_RT4, GSM7681692_RT6).

Raw data in 10X format were processed using the Seurat package (v4.0.0) in R. For each sample, expression matrices were generated using the ReadMtx function, and Seurat objects were created with minimal quality control thresholds (minimum 3 cells per gene and 200 features per cell). Sample-specific metadata, including group information (NT, PT, RT) was added to each object. Mitochondrial gene percentage was calculated for each cell using the PercentageFeatureSet function. Seurat objects were merged into a single dataset while preserving sample identities. Quality control filtering removed cells with fewer than 200 or more than 6,000 features and cells with mitochondrial gene content exceeding 20%. The filtered dataset underwent log normalization and identified 2,000 highly variable genes using the variance-stabilizing transformation method.

Data were scaled to remove technical variation, and principal component analysis (PCA) was performed. We employed Harmony integration (Korsunsky et al., 2019) with sample identities as batch variables to mitigate batch effects across samples. Uniform Manifold Approximation and Projection (UMAP) was used for dimensionality reduction, using the first 30 principal components before and after batch correction. Silhouette scores were calculated to assess the removal of batch effects quantitatively. Graph-based clustering was performed at a resolution of 0.5 using the FindNeighbors and FindClusters functions with Harmony-integrated dimensionality reduction. Marker genes for each cluster were identified using the FindAllMarkers function with a minimum log fold-change threshold of 0.25 and detection in at least 25% of cells within a cluster.

#### Patient-derived xenograft (PDX) scRNA-seq dataset analysis

Single-cell RNA sequencing data from patient-derived xenograft (PDX) breast cancer models under three treatment conditions (control, estrogen [E2], and estrogen plus fulvestrant [E2+Fulv.]) were obtained from the GEO dataset GSE213733 (35). Raw count matrices were processed using the Seurat package in R (v4.4.2).

Quality control was performed to remove low-quality cells and potential doublets. Cells with high mitochondrial content (>20%), extremely high feature counts (>6,000 genes), or extremely high UMI counts (>30,000) were excluded. Human and mouse genes were distinguished by their "hg19-" and "mm10-" prefixes. For downstream analyses, we focused primarily on human gene expression to better capture tumor cell-specific responses to treatments. After quality control, the data were normalized using Seurat’s standard log-normalization method. Variable features were identified among human genes using the FindVariableFeatures function with default parameters. Data were scaled, and principal component analysis (PCA) was performed using human genes. To correct batch effects across treatment groups while preserving biological variation, we applied Harmony integration using the IntegrateLayers function with the HarmonyIntegration method. Cells were clustered using the FindNeighbors and FindClusters functions (resolution = 0.5).

#### Gene Signature and Module Score Analysis

Four gene signatures were analyzed: (1) a 46-gene PML1 signature including genes (Table S1); (2) a MAPK pathway signature from MSigDB C6: oncogenic signature gene sets-MEK_UP.V1_UP comprising 195 genes (Table S1); and (3) a PI3K pathway signature from MSigDB Hallmark gene sets: HALLMARK_PI3K_AKT_MTOR_SIGNALING comprising of 105 genes (Table S1); an endocrine resistance signature from Kim *et. al* (PMID: 37747807) comprising of 307 genes. Additionally, we assessed epithelial cell identity using a 16-gene signature, including canonical markers (*EPCAM, KRT8, KRT18, KRT19*) and breast epithelium-specific markers (*KRT7, CLDN3, CLDN4, GATA3*).

Module scores for each signature were calculated using the AddModuleScore function in Seurat. The cell cycle phase was determined using the CellCycleScoring function with canonical S and G2/M phase gene sets. Epithelial cells were identified based on a threshold applied to the epithelial score.

Statistical analysis of module scores across sample groups was performed using pairwise Wilcoxon rank-sum tests with Bonferroni correction for multiple comparisons. For specific pathway comparisons across the three tissue types, pairwise t-tests with Benjamini-Hochberg correction were employed. Results were visualized using ggplot2 with boxplot and violin plot representations to highlight group score distributions. Cellular identities, gene expression patterns, and module scores were visualized using standard Seurat visualization functions (DimPlot, FeaturePlot) and the scRNAtoolVis package. All analyses were conducted in Windows R (v4.4.2).

### Spatial transcriptomics analysis

Spatial transcriptomics analysis was performed using the 10x Genomics Visium platform on breast cancer tissue sections. The preprocessed Seurat object containing spatial gene expression data was analyzed using the R package Seurat (v4.4.2) with the Space Transcriptomics Component (SCT) normalization. Multiple gene signatures were evaluated to characterize tumor heterogeneity and pathway activation. The calculation of these signatures followed the method used in the 10× Genomics coupe browser, based on log_ average expression. These included (1) a 46-gene PML1 signature, (2) a 170-gene MAPK pathway signature, (3) a 97-gene PI3K pathway signature, and (4) a 307-gene endocrine resistance signature. Signature genes are listed in Table S2.

We developed a classification approach based on canonical markers to distinguish tumors from stromal regions. We quantified a tumor signature comprising epithelial and proliferation markers (*EPCAM, KRT8, KRT18, KRT19, MKI67, PCNA, KRT7, CCNB1, TOP2A, CDK1*) and a stromal signature including fibroblast and extracellular matrix components (*COL1A1, COL1A2, VIM, DCN, LUM, FAP, PDGFRB, ACTA2, COL3A1, FN1*) for each spatial spot. The tumor proportion was calculated as the ratio of tumor signature expression to the sum of tumor and stromal signatures. Spatial spots with tumor proportion exceeding 0.2 were classified as tumor-enriched regions for subsequent tumor-specific analyses.

Gene signatures were visualized using SpatialFeaturePlot with customized color gradients optimized for tissue architecture interpretation. For tumor-specific pathway analyses, signature scores were calculated separately for tumor-enriched regions, and correlations between pathway activities were assessed using Pearson correlation coefficients. Correlation matrices and pairwise scatter plots were generated to evaluate relationships between PML1, MAPK, PI3K, and endocrine resistance signatures within the tumor microenvironment. Statistical analyses included correlation testing with p-value calculation for all pathway pairs and comparing correlation patterns between tumor-enriched and whole-tissue regions. Visualization was performed using ggplot2 and specialized correlation plotting packages (corrplot, GGally) with customized aesthetics to highlight significant relationships.

### RNA-seq analysis

RNA sequencing was performed using a comprehensive RNA-seq analysis pipeline in a high-performance computing environment. Quality control of raw sequencing reads was performed using FastQC (v0.12.1), and adapter sequences were trimmed using Trim Galore (v0.6.10) with Cutadapt (v4.9). Reads passing quality thresholds (minimum 10,000 reads per sample after trimming) were retained for downstream analysis. Read quality was assessed using FQ (v0.12.0) for linting and subsampling. High-quality reads were aligned to the GRCh38 human reference genome using STAR (v2.7.10a). The alignment was performed using genome indices and annotation files obtained from NCBI’s GRCh38 build. Following alignment, transcript quantification was performed using RSEM (v1.3.1) to estimate gene and transcript expression. Alignment files were processed using Samtools (v1.21) for sorting, indexing, and generating alignment statistics (flagstat, idxstats, and stats). Duplicate reads were marked using Picard MarkDuplicates (v3.1.1). For alignment quality assessment, RSeqC (v5.0.2) modules were implemented including bam_stat, inner_distance, infer_experiment, junction_annotation, junction_saturation, read_distribution, and read_duplication. These modules provided comprehensive metrics on mapping quality, strand specificity, and library complexity. Qualimap (v2.3) was used for additional RNA-seq-specific quality control. For visualization purposes, genome coverage was calculated using BEDTools (v2.31.1), and bigWig files were generated using UCSC utilities (bedClip v377, bedGraphToBigWig v469). Gene expression was also analyzed using StringTie (v2.2.3). Alternative quantification was performed using Salmon (v1.10.3) for transcript-level abundance estimation. For expression analysis, DESeq2 (v1.28.0, using R v4.0.3) variance stabilizing transformation was applied to normalize count data. The reference transcriptome was prepared using RSEM’s prepare-reference with STAR genome indices. Python scripts (v3.9.5) were utilized for GTF filtering and processing. All analysis steps were performed according to best practices for RNA-seq data processing, with all software configurations and parameters recorded for reproducibility.

We performed Gene Set Variation Analysis (GSVA) on RNA sequencing data from GSE89888, which includes transcriptome profiles of T47D and MCF7 breast cancer cell lines expressing either wild-type *ESR1* or *ESR1* with Y537S (YS) or D538G (DG) mutations. Log2-transformed CPM expression matrices were analyzed using four curated gene sets: MAPK signaling (190 genes), PI3K signaling (114 genes), PML1-associated genes (46 genes), and endocrine resistance (307 genes). GSVA enrichment scores were calculated. Differential pathway activity between mutants and wild-type samples was assessed using limma, with empirical Bayes moderation to compute statistics. We constructed design matrices with cell types (WT, YS, DG) as factors and established contrast matrices for YS vs. WT and DG vs. WT comparisons. Statistical significance was determined using adjusted p-values (Benjamini-Hochberg correction), with p<0.05 considered significant. Results were visualized using ggplot2 and pheatmap packages, where bubble plot point sizes represented GSVA scores and color intensity reflected statistical significance.

### MBC cohort analysis

#### Genomic Alteration Analysis in Metastatic Breast Cancer

We analyzed somatic mutations and copy number alterations in metastatic breast cancer samples using genomic data from the MBC (Metastatic Breast Cancer) cohort (43). We investigated potential gene-gene interactions between PML and other key breast cancer driver genes, including *ESR1, TP53, PIK3CA, PTEN, EGFR, ERBB2, KRAS, BRAF, GATA3, FOXA1, MYC, CTCF, MAP3K1, CDH1, AKT1, MTOR, NF1, FGFR1, ERBB3,* and *TBX3*.

Mutations were classified as missense mutations, nonsense mutations, frameshift insertions/deletions, splice site alterations, and multi-hit mutations (when multiple mutations occurred in the same gene in a single sample). Copy number alterations were categorized as either amplifications (≥2 copies) or homozygous deletions (≤-2 copies). Sample identifiers were standardized to ensure proper integration between datasets.

We generated two separate matrices to evaluate potential co-occurrence or mutual exclusivity between gene alterations: (1) a matrix including all PML alterations and (2) a matrix including only PML amplifications. We performed Fisher’s exact test on each gene pair for both matrices to assess significant associations. We created a contingency table for each gene pair and compared the observed co-occurrence frequency to the expected frequency under the null hypothesis of independence.

For each gene pair, we calculated the log_2_ratio between observed and expected co-occurrence frequencies to quantify the strength and direction of association. Positive log_2_ratios indicated co-occurrence, while negative values suggested mutual exclusivity. P-values were adjusted for multiple testing using the Benjamini-Hochberg false discovery rate (FDR) method, with adjusted p-values < 0.05 considered statistically significant.

We visualized results using a half-triangle bubble plot, with the upper left triangle representing all PML alterations and the lower right triangle representing PML amplifications only. Bubble size represented statistical significance (-log10 of adjusted p-value), and color indicated the log_2_ratio (blue for mutual exclusivity, red for co-occurrence). This dual visualization approach allowed for comparing interaction patterns of all PML alterations versus specifically PML amplifications, providing insights into the functional relationships between PML and other cancer-related genes in metastatic breast cancer.

#### Transcriptomics analysis

For the MBC cohort analysis, we employed a similar approach to the TCGA transcriptomics analysis, focusing on comparing gene expression signatures across different breast cancer subtypes. We examined the relationships between PML1 signature, MAPK pathway activation, PI3K signaling, and endocrine resistance gene expression patterns. Expression data was processed from RNA-seq datasets. We utilized the singscore R package to calculate signature scores for each sample based on pre-defined gene sets (Table S1). This approach was applied to both the entire breast cancer cohort and to a subset analysis focusing specifically on ER-positive breast cancer samples. Spearman correlation analysis was then performed to assess the correlations between the expression patterns of these gene signatures.

We implemented correlation analyses using Spearman methods to investigate relationships between the PML1 signature and other genes. The resulting correlation data included correlation coefficients, p-values, and false discovery rates (FDR) calculated using the Benjamini-Hochberg method for multiple testing correction. Results were sorted based on correlation strength and statistical significance to identify genes most strongly associated with the PML1 signature. For visualization of the genome-wide correlation results, we developed custom volcano plotting functions to display the relationship between correlation strength (x-axis) and statistical significance (y-axis, represented as -log10(FDR)). To contextualize the gene correlation results within known oncogenic pathways, we performed Gene Set Enrichment Analysis (GSEA) using the fgsea package. We utilized the MSigDB C6 oncogenic signatures collection to identify enriched pathways associated with the PML1 signature correlation pattern. The GSEA was configured with parameters including a minimum gene set size of 15, a maximum of 500, and 10,000 permutations to establish statistical significance. For significant pathway associations, enrichment plots were generated to visualize the distribution of pathway genes across the ranked correlation list, with corresponding normalized enrichment scores (NES) and adjusted p-values.

### CPTAC Proteomics Data Analysis and Gene Co-expression Clustering

#### Proteomics Dataset Processing

Breast cancer proteomics data were obtained from the Clinical Proteomic Tumor Analysis Consortium (CPTAC) database. The analysis focused on estrogen receptor (ER)-positive breast tumors, which were identified based on ER clinical status in accompanying clinical metadata. A total of 81 ER-positive samples were included in the analysis. Protein expression values were extracted from the complete proteome dataset, and gene symbols were used as identifiers. For proteins with multiple isoforms, expression values were consolidated by summing the intensities across all detected isoforms for each protein, resulting in a non-redundant gene-level expression matrix.

#### Correlation Analysis with PML

To identify proteins functionally related to PML in ER-positive breast cancers, we performed a systematic correlation analysis between PML and all other proteins in the dataset. We calculated Spearman rank correlation coefficients for each protein-PML pair to assess linear and non-linear relationships while minimizing the influence of outliers. Statistical significance was determined using correlation tests with adjustment for multiple hypothesis testing using the Benjamini-Hochberg false discovery rate (FDR) method. Proteins with absolute correlation coefficients greater than 0.25 and adjusted p-values less than 0.05 were considered significantly associated with PML. This approach identified proteins whose expression patterns were coordinated considerably with PML in ER-positive breast tumors.

#### Fuzzy Clustering Analysis

Significant PML-correlated proteins were further analyzed using fuzzy c-means clustering implemented in the ClusterGVis R package to identify distinct co-expression patterns. The optimal number of clusters was determined by evaluating cluster stability metrics using the getClusters function. Nine distinct co-expression clusters were identified and visualized. We performed functional enrichment analysis using the Kyoto Encyclopedia of Genes and Genomes (KEGG) pathway for each cluster. A PML1 signature gene set comprising 46 genes was specifically highlighted in the visualization to assess their distribution across the identified co-expression clusters. The expression matrix was ordered based on PML expression levels to reveal potential gradient effects.

#### Visualization and Statistical Analysis

Heatmaps and cluster visualizations were generated using the ClusterGVis package with customized parameters to highlight gene co-expression patterns. Expression patterns were displayed alongside functional enrichment results to facilitate biological interpretation. Statistical summaries of correlation analyses were visualized using ggplot2, with color coding to distinguish statistically significant associations (p < 0.05). All analyses were performed in R (version 4.0). The correlation between PML and signature genes was visualized as bar plots with significance thresholds indicated, providing a quantitative assessment of the PML-associated proteome in ER-positive breast cancer.

#### CPTAC-Phosphoproteomic Analysis

For phosphoproteomic analysis, we obtained CPTAC phosphoprotein quantification data from cBioPortal. We extracted PML phosphorylation sites from the dataset, focusing on site-specific phosphorylation patterns. To investigate the relationship between MAPK pathway activity and PML phosphorylation, we performed a multi-omics analysis integrating transcriptomic, proteomic, and phosphoproteomic data.

First, we calculated MAPK pathway activity scores from RNA-seq data using the GSVA package in R with a curated MAPK signature. The calculation utilized the gsvaParam and gsva functions to generate sample-specific enrichment scores that reflect MAPK pathway activation levels.

PML phosphorylation data were extracted and organized by phosphorylation site, with each site identified by its specific amino acid residue and position. We identified multiple PML phosphorylation sites in the dataset and transformed the data to enable correlation analysis with pathway activity scores. For integrative analysis, we identified the same samples between the MAPK activity score dataset and PML phosphorylation data, merging them into a unified analysis matrix.

Correlation analysis was performed between MAPK activity scores and each PML phosphorylation site using Pearson’s correlation test. We calculated correlation coefficients, p-values, and adjusted p-values for each phosphorylation site using the Benjamini-Hochberg method to control for multiple tests. Correlation results were ranked by absolute correlation strength to identify the most strongly associated phosphorylation sites.

Finally, this phosphoproteomic data was integrated with PML protein expression data to establish relationships between total PML protein levels, specific phosphorylation sites, and MAPK pathway activation. This three-component analysis framework provided insights into the regulatory mechanisms connecting signaling pathway activity with post-translational modifications of PML.

### Breast cancer clinical dataset analyses

#### Microarray Dataset Analysis

Two independent breast cancer microarray datasets (GSE9893 and GSE124647) (59,60) with survival data were downloaded from the Gene Expression Omnibus (GEO). Expression data were normalized using the limma normalizeBetweenArrays function. Multiple probes targeting the same gene were averaged for probe-to-gene mapping, and gene symbols indexed the resulting expression matrices.

#### PML1 Signature Score Calculation

A 46-gene PML1 signature was defined based on genes associated with PML1 isoform expression and cellular functions. We applied the singScore R package to calculate transcriptional signature scores for the PML1 gene set in each sample.

#### Survival Analysis

Kaplan-Meier survival analyses were performed for progression-free survival (PFS) and overall survival (OS). PFS time was defined as time to recurrence (local or distant metastases) or end of follow-up, with censoring for patients without recurrence. OS time was defined as the follow-up period for all patients. Event status was coded as 1 for recurrence (PFS), death (OS), and 0 for censoring.

Breast cancer patients were assigned into high and low PML1 signature groups based on the cut-off point estimated using the maximally selected rank statistic via the maxstat R package. Survival curves were compared using the log-rank test, and hazard ratios were calculated using Cox proportional hazards regression models. Results are presented with 95% confidence intervals and adjusted p-values; statistical significance was set at p < 0.05.

### Statistical analysis

Continuous variables were compared between the two groups using Student’s t-test for statistical analysis. For comparisons involving more than two groups, we applied a one-way analysis of variance (ANOVA), followed by posthoc tests. Specifically, Tukey’s test was used for multiple pairwise comparisons between all groups, while Dunnett’s test was employed when comparisons were only needed against a control group.

Statistical significance was defined as p < 0.05 and indicated in figures as follows: *p < 0.05, **p < 0.01, ***p < 0.005, and ****p < 0.0001. All statistical analyses were performed using R (version 4.4.2) or GraphPad Prism (version 9.0). For correlation analyses, Pearson’s or Spearman’s correlation coefficients were calculated as appropriate based on data distribution, with the same significance thresholds applied. When performing extensive statistical tests, multiple testing correction was applied using the Benjamini-Hochberg procedure to control the false discovery rate. Data are presented as mean ± SEM unless otherwise specified.

## Supplementary Figures

**Fig. S1.**
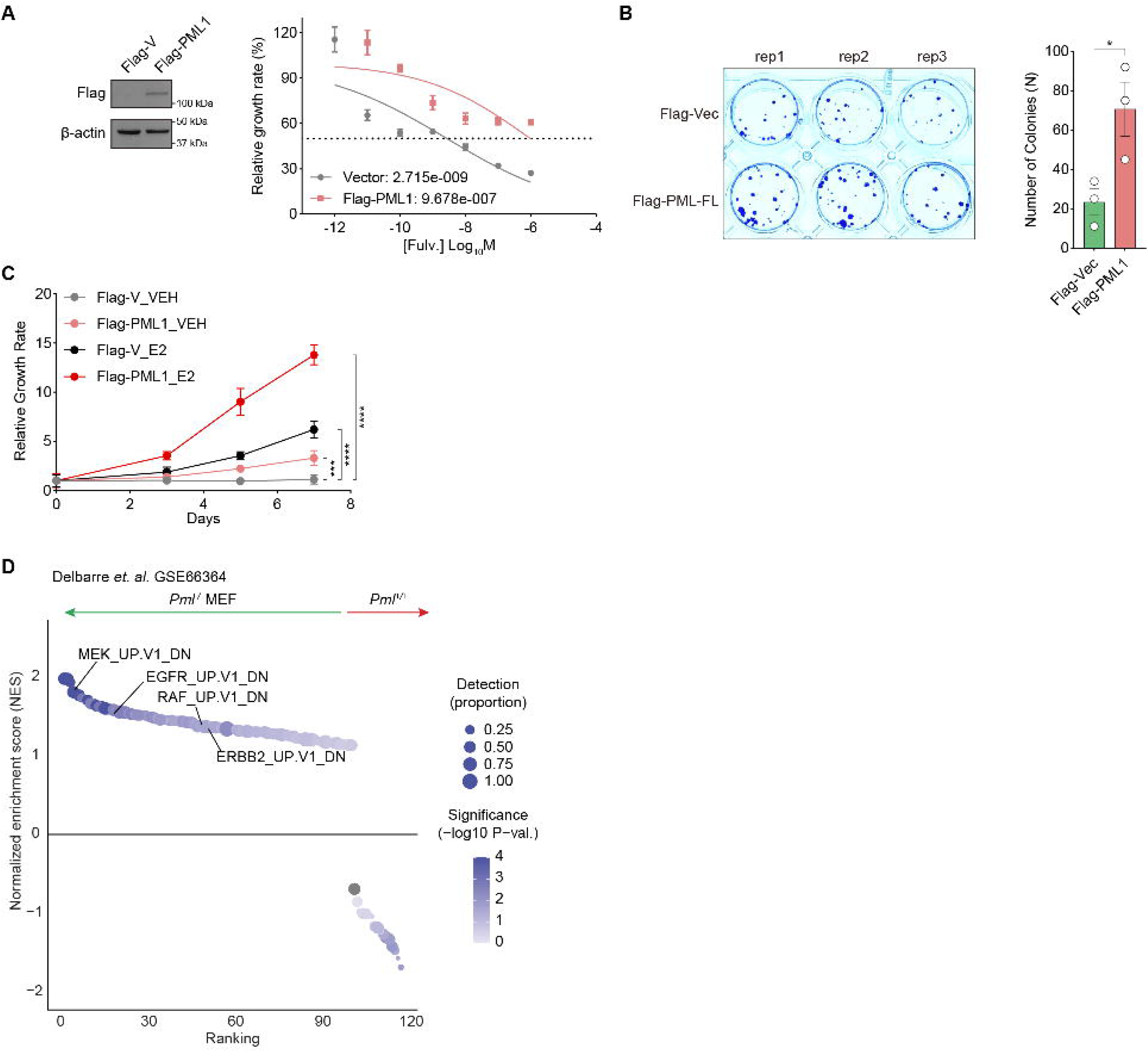
PML1 expression promotes endocrine resistance and oncogenic signaling. ***A-C***, Ectopic expression of Flag-PML1 induces Fulv resistance **(*A*),** enhances colony formation **(*B*),** and enables estrogen-independent growth **(*C*).** ***D*,** RNA-seq analysis of *Pml^+/+^* versus *Pml^-/-^*MEFs reveals enrichment of oncogenic signaling pathways, positively correlating with MEK, EGFR, RAF, and ERBB2 activation signatures.

**Fig. S2.**
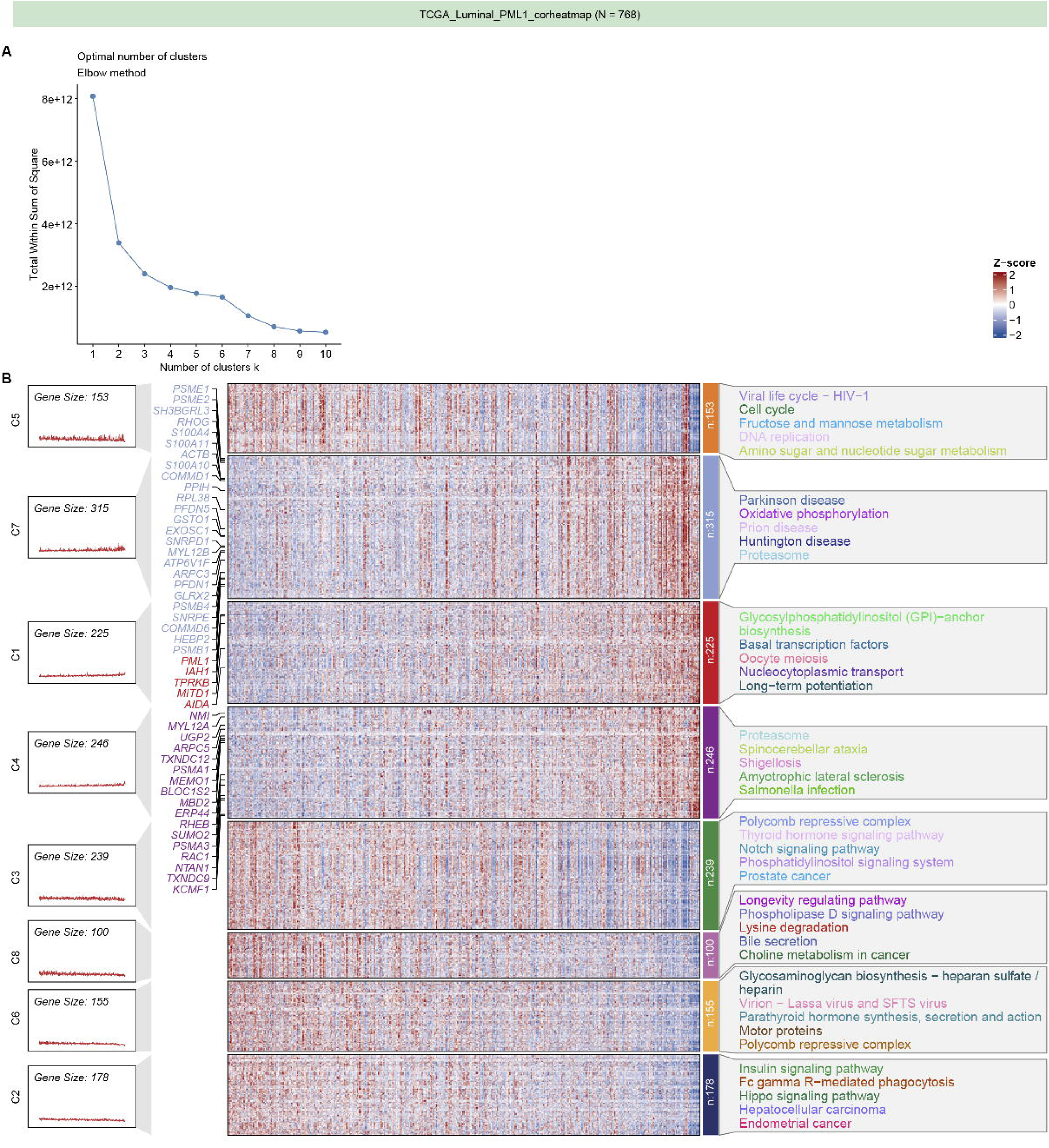
Transcriptomic analysis of TCGA ER+ breast tumors reveals PML1-associated gene networks in ER+ breast tumors. ***A,*** Elbow method plot showing optimal cluster number determination, with the Total Within Sum of Square (WSS) against the number of clusters (k). ***B,*** Heatmap of gene transcripts clustered by association with *PML1* mRNA expression. PML1 signature genes are highlighted within clusters. Associations are defined by a correlation coefficient (|r| > 0.25) and an FDR-adjusted *p*-value < 0.05. KEGG pathways for each cluster are indicated.

**Fig. S3.**
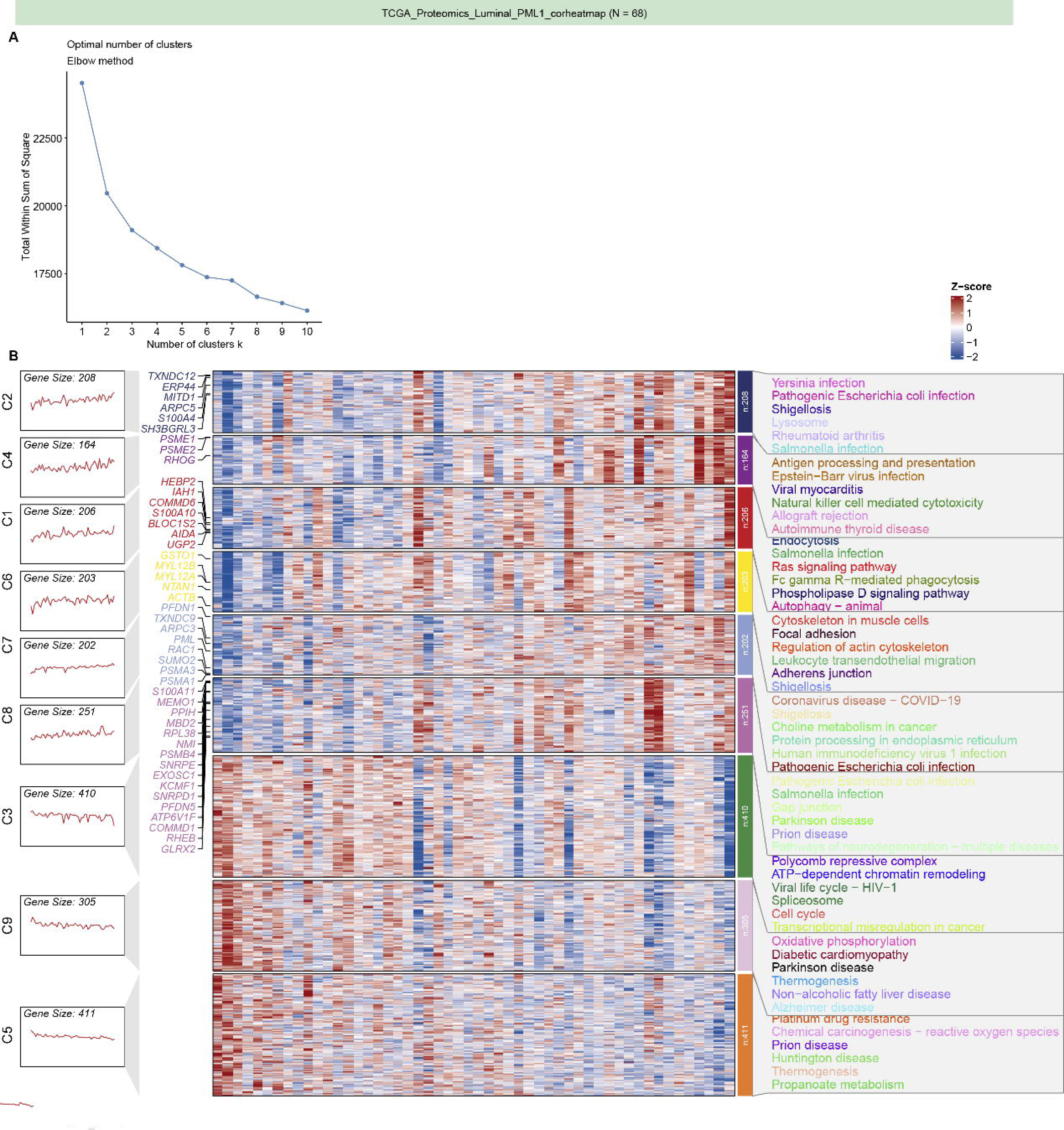
TCGA proteomics analysis of PML-associated proteins in ER+ breast tumors. ***A,*** Elbow method plot determining optimal cluster number by plotting Total Within Sum of Square (WSS) against number of clusters (k). ***B,*** Heatmap showing protein clusters associated with total PML protein abundance. PML1 signature genes are highlighted. Associations defined by correlation coefficient (|r| > 0.25) and FDR-adjusted *p*-value < 0.05. KEGG pathways for each cluster are indicated.

**Fig. S4.**
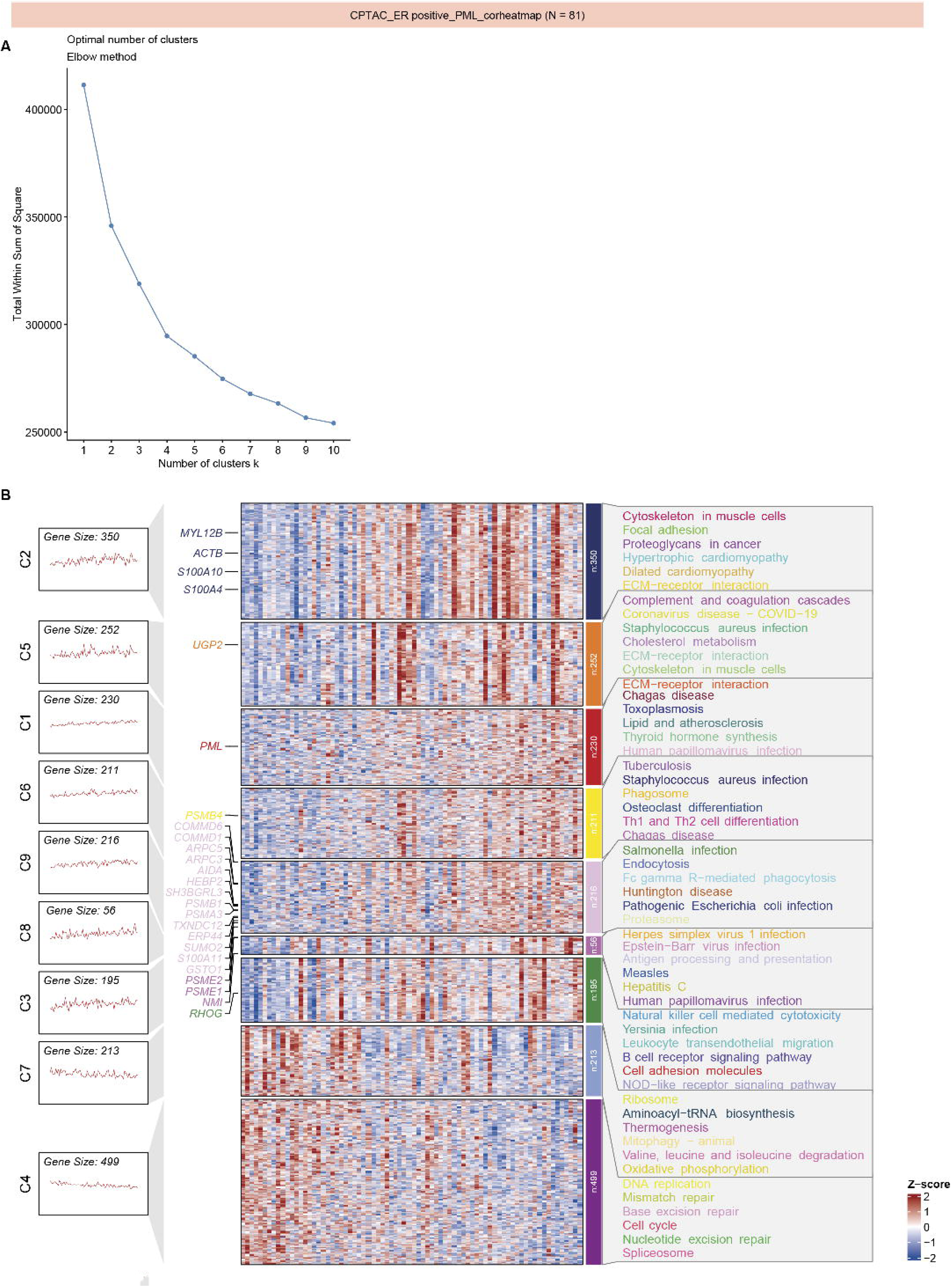
CPTAC proteomics analysis of ER+ breast tumors shows PML-associated protein networks in ER+ breast tumors. ***A,*** Elbow method plot showing optimal cluster number determination. ***B,*** Heatmap of protein clusters associated with total PML protein abundance. PML1 signature genes are highlighted. Associations defined by correlation coefficient (|r| > 0.25) and FDR-adjusted *p*-value < 0.05. KEGG pathways for each cluster are indicated.

**Fig. S5.**
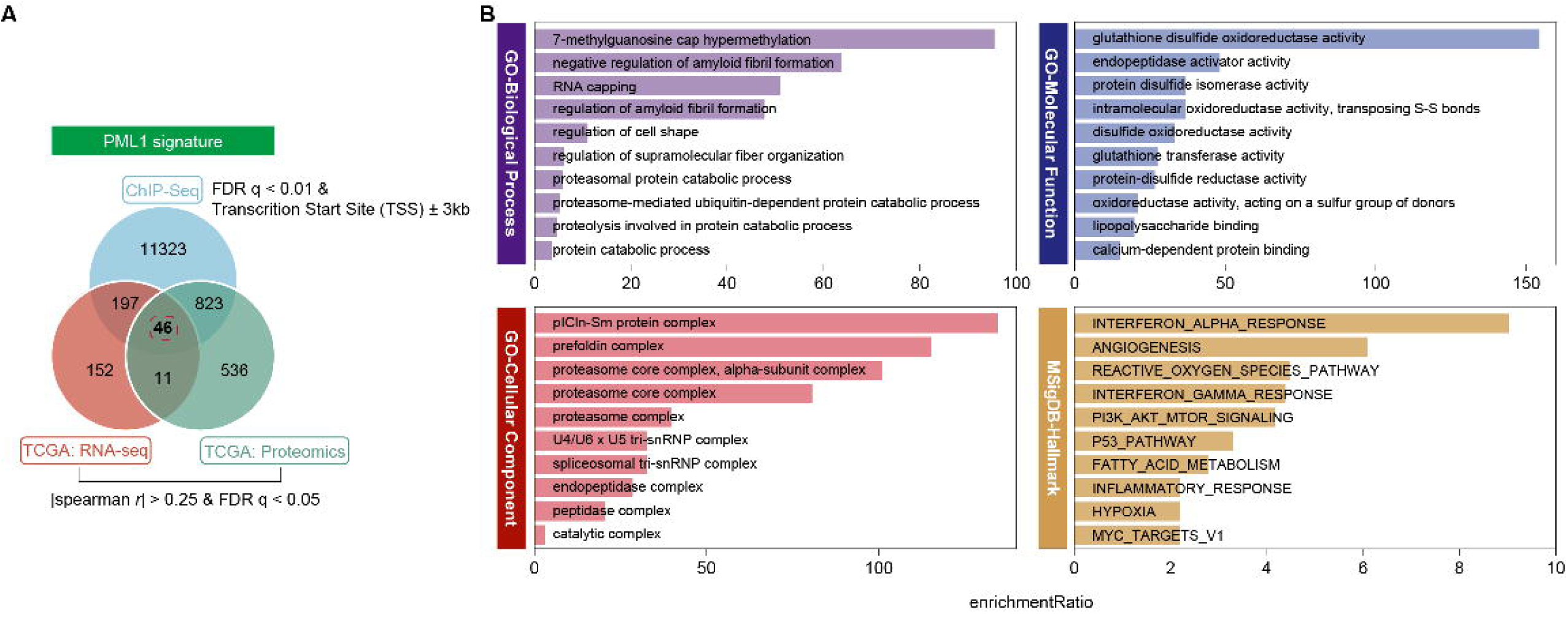
Multi-omics integration of ChIP-seq, RNA-seq, and proteomics analyses define the PML1 gene signature comprising 46 genes. ***A***, A Venn diagram showing the overlap of PML1-associated genes identified across ChIP-seq, TCGA RNA-seq, and TCGA proteomics datasets. ***B,*** Gene Ontology (GO) enrichment analysis of the PML1 gene signature categorized by Biological Processes, Molecular Function, Cellular Component, and Hallmark pathways.

**Fig. S6.**
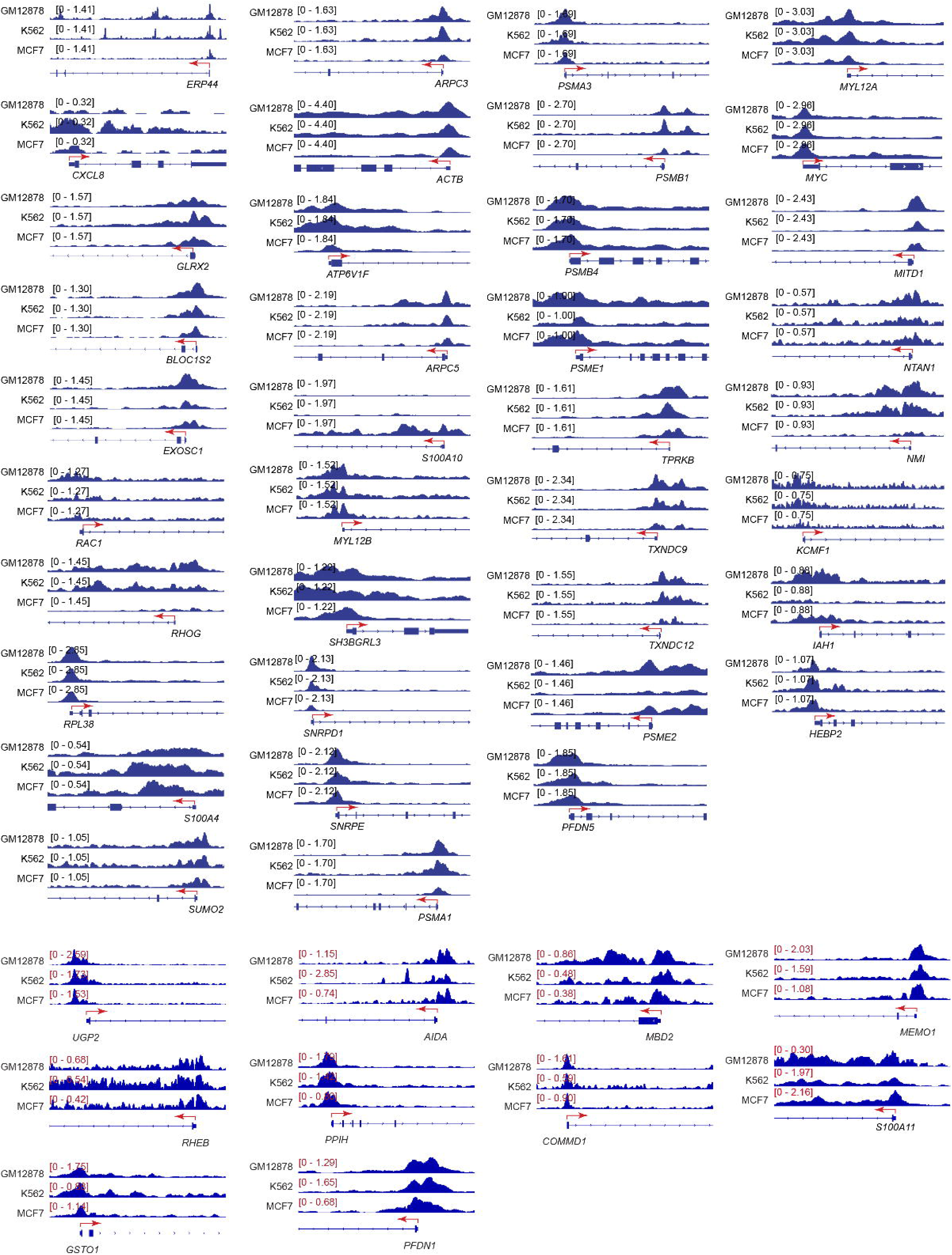
PML binding to signature gene promoters across multiple cell lines. Genome tracks showing PML binding sites on PML1 signature genes in GM12878, K562, and MCF-7 cells. Binding intensity normalized to the same scale for each gene, except *UGP2*, AIDA, *MBD2*, *MEMO1*, *RHEB*, *PPIH*, *COMMD1*, *S100A11*, *GSTO1*, and *PFDN1*.

**Fig. S7.**
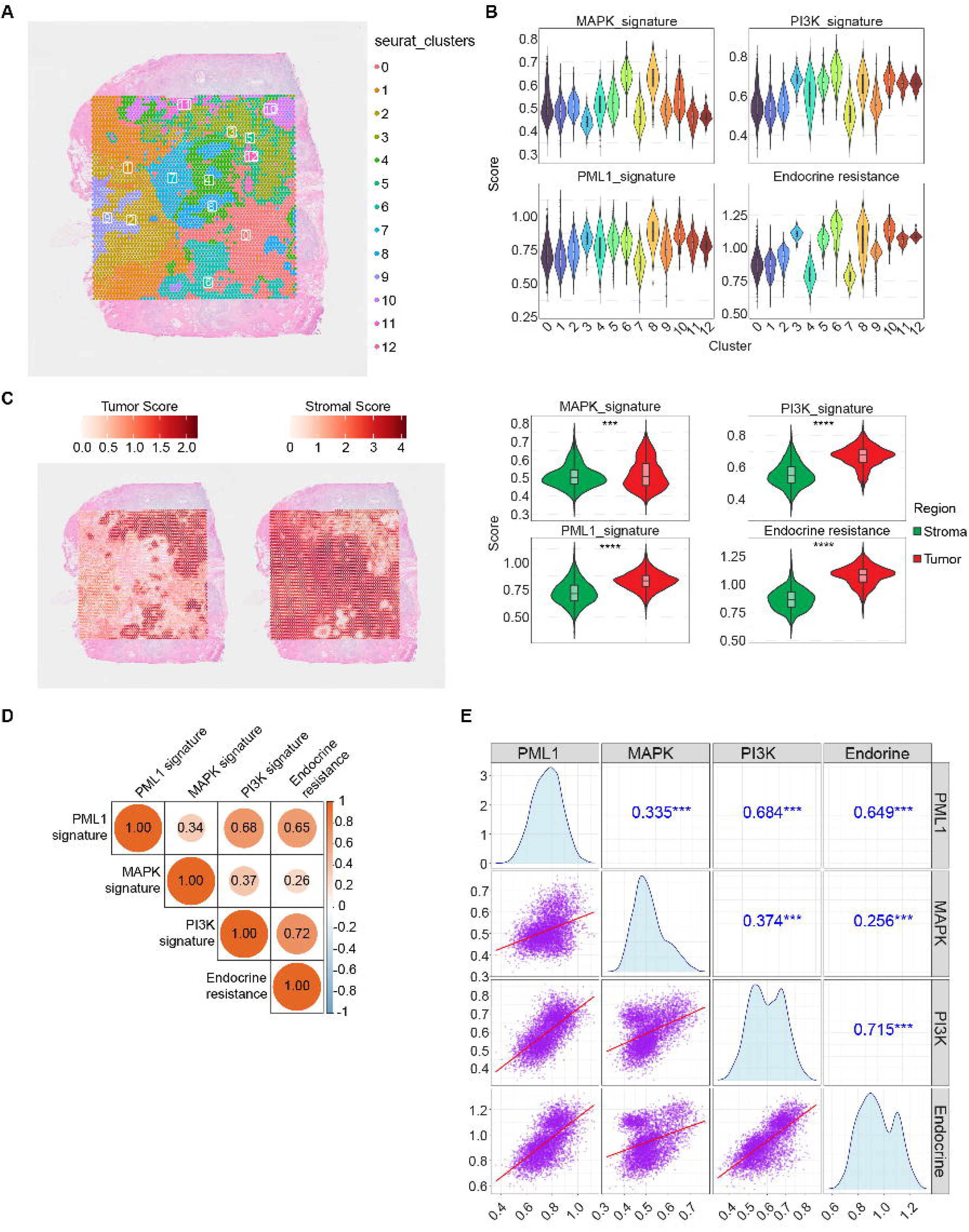
Spatial transcriptomics reveals strong correlations between PML1, PI3K, MAPK, and endocrine resistance gene signatures. ***A,*** Spatial transcriptomic analysis of tumor section showing H&E staining with Seurat clusters represented by color-coded hexagonal spots identifying 13 distinct cell clusters. ***B*,** Violin plots showing MAPK, PI3K, PML1, and endocrine resistance signatures across the 13 identified clusters. ***C,*** *Left*, Tumor and stroma cell scores across tumor tissue regions. *Right*, Violin plots demonstrating higher MAPK, PI3K, PML1, and endocrine resistance signature scores in tumor versus non-tumor sections. ***D,*** Correlation matrix showing relationships between PML1, MAPK, PI3K, and endocrine resistance gene signatures. ***E,*** Diagonal plots showing each gene signature’s distribution (histograms/density plots), with the upper triangle displaying correlation coefficients and *p*-values and the lower triangle showing pairwise scatterplots with regression lines (in red).

**Fig. S8.**
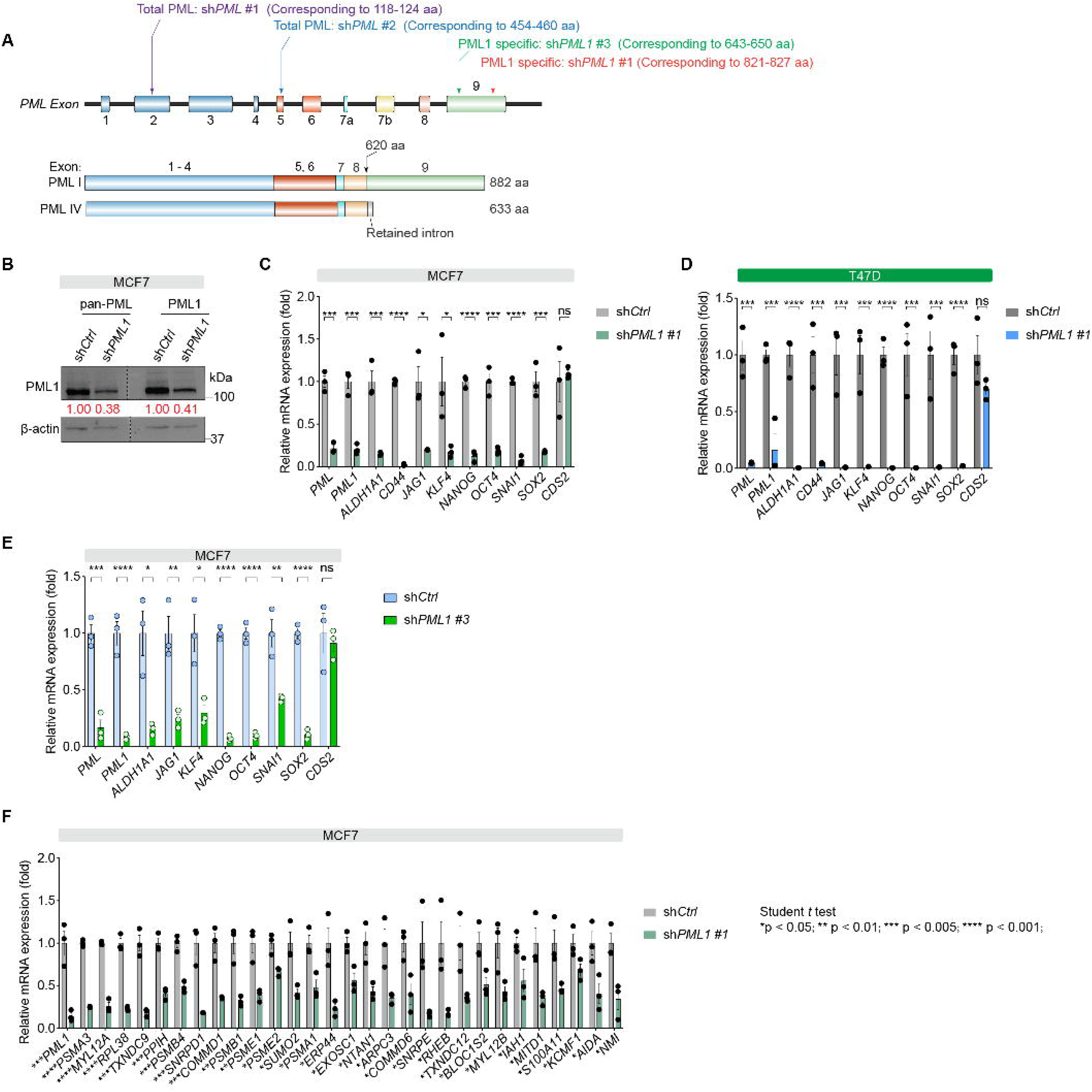
*PML1*-specific knockdown has similar effects to total *PML* knockdown on BCSC-associated and PML1 signature genes. ***A***, PML gene structure indicating target sites for total *PML* and *PML1*-specific shRNAs. ***B,*** Western blot validation of *PML1*-specific shRNA effects on total PML and PML1 protein levels. ***C-D***, Effects of *PML1*-specific shRNA (sh*PML1* #1) on the expression of BCSC-associated genes in MCF-7 (***C***) and T47D cells (***D***). ***E,*** Effects of *PML1*-specific shRNA (sh*PML1* #3) on the expression of BCSC-associated genes. ***F***, Effects of *PML1*-specific shRNA (sh*PML1* #1) on the expression of PML1 signature genes.

**Fig. S9.**
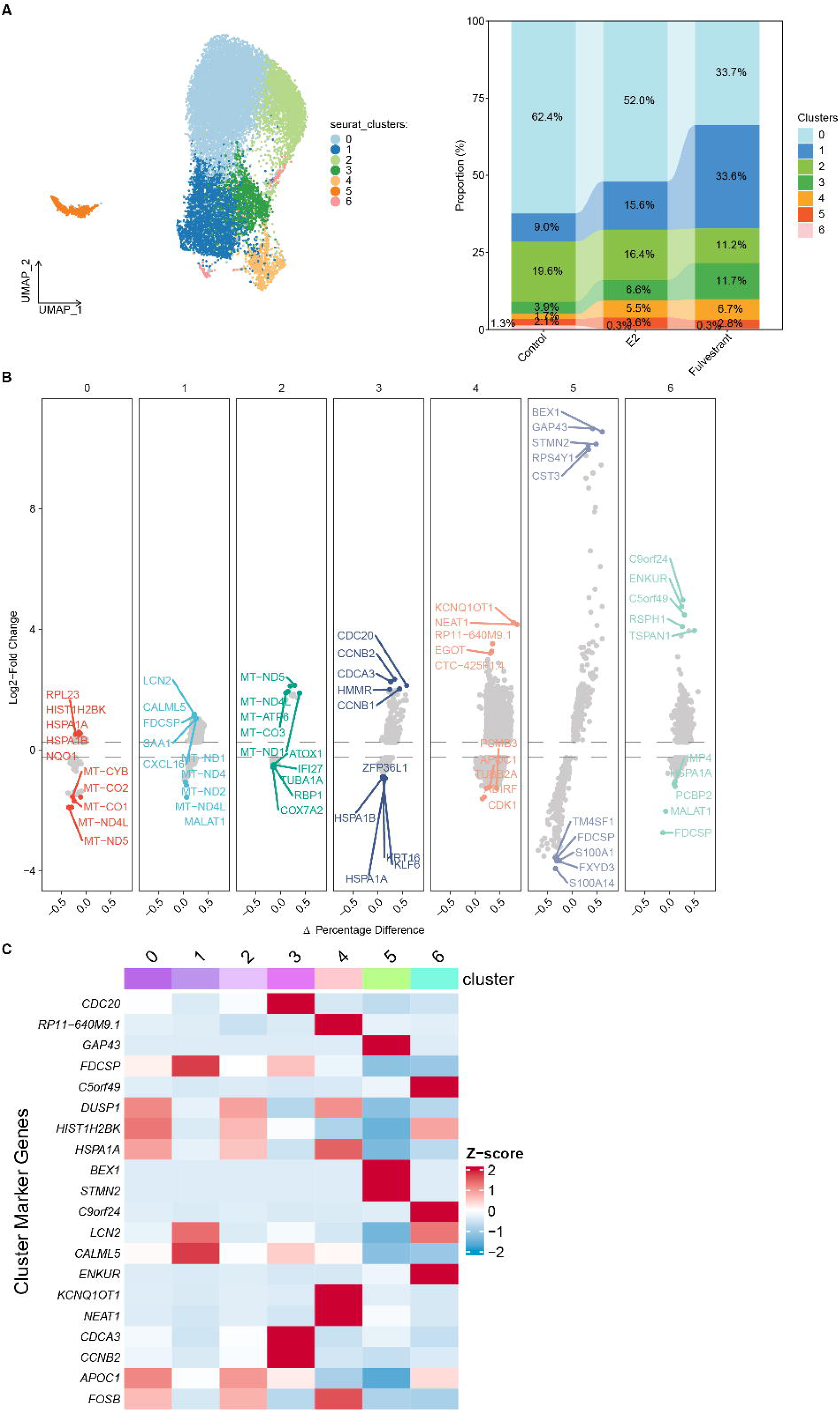
scRNA-seq analysis of E2-accelerating Pdx breast tumor. ***A,*** UMAP visualization of cell clusters identified by Seurat analysis showing distribution into 7 groups (*left*) with a stacked bar chart showing cluster proportions across three treatment groups (*right*). ***B***, Volcano plots displaying differential gene expression across 7 distinct cell clusters showing the relationship between expression change (Log2 Fold Change on y-axis) and Δ Percentage Difference (x-axis). ***C***, Heatmap of marker gene expression patterns across 7 cell clusters.

**Fig. S10.**
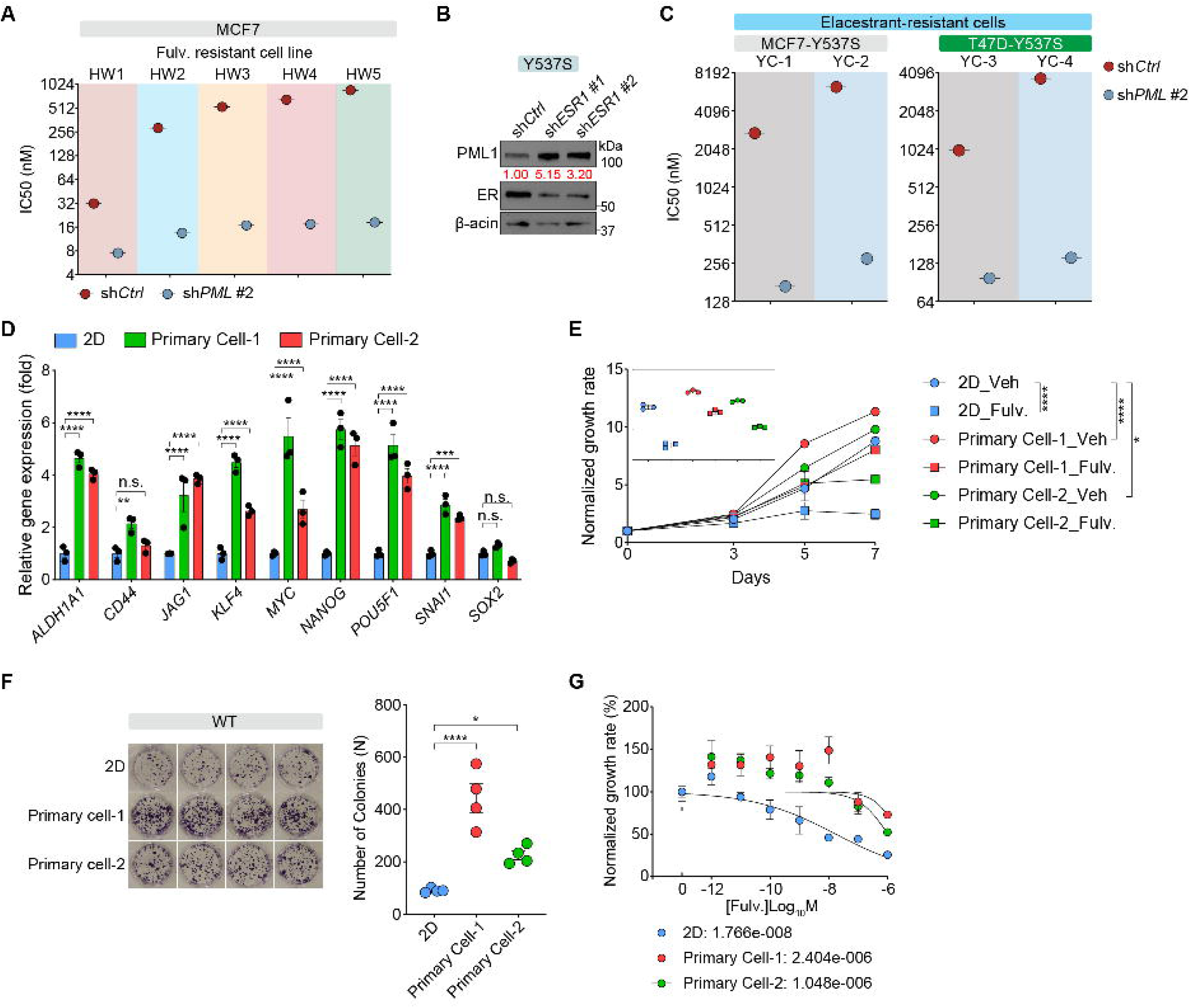
Primary tumor-derived cells exhibit enhanced tumorigenic properties compared to cultured MCF-7 cells. ***A***, *PML* knockdown significantly reduced Fulv IC50 in Fulv-resistant cells. ***B,*** Loss of *ESR1* by knockdown increases PML1 protein abundance in ^Y537S^ER-KI MCF-7 cells. ***C,*** *PML* knockdown significantly reduced ELA IC50 in ELA-resistant MCF-7 and T47D cells. ***D,*** Elevated expression of BCSC-associated genes in primary tumor cells compared to cultured MCF-7 cells. ***E,*** Enhanced proliferative potential in primary tumor cells compared to cultured MCF-7 cells. ***F***, Increased colony formation capacity in primary tumor cells compared to cultured MCF-7 cells. ***G***, Higher Fulv IC50 values in primary tumor-derived cells compared to MCF-7 cells

**Fig. S11.**
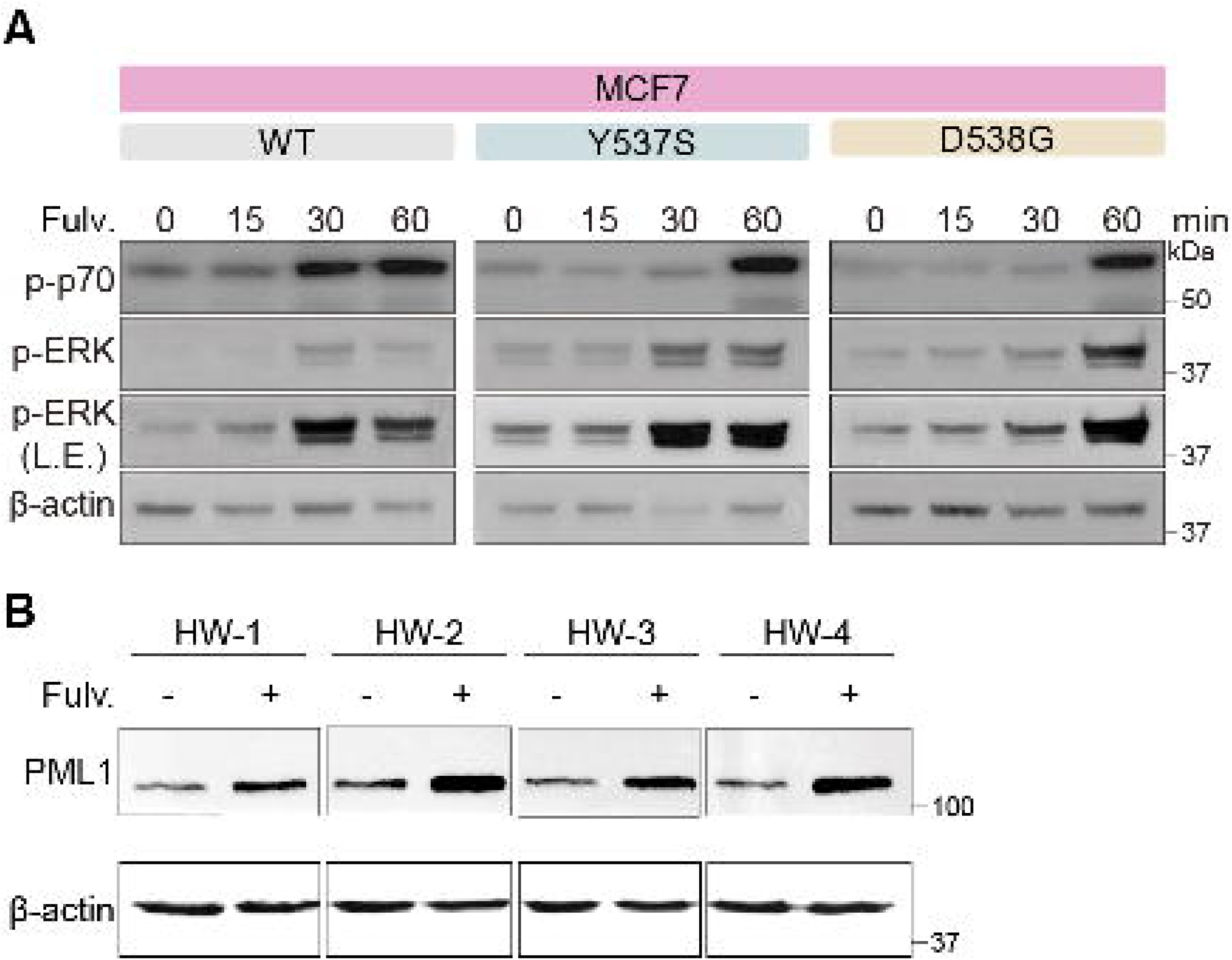
Fulv induces PML1 protein expression in Fulv-resistant cell variants. ***A***, Fulv triggers MEK-ERK and mTOR-p70 activation within one hour, preceding PML1 protein accumulation. ***B***, Increased PML1 protein levels in Fulv-resistant MCF-7 cell variants following Fulv treatment.

**Fig. S12.**
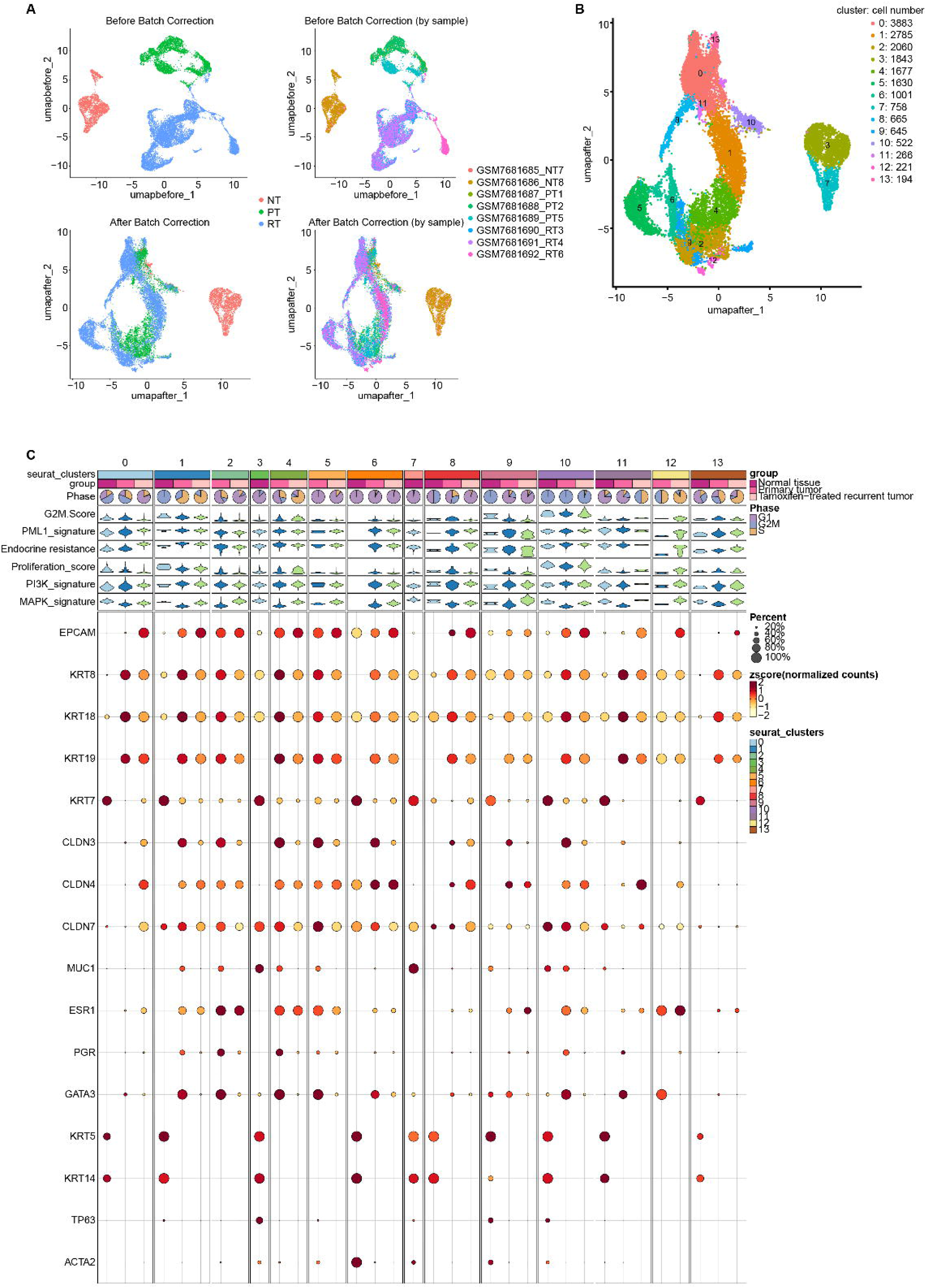
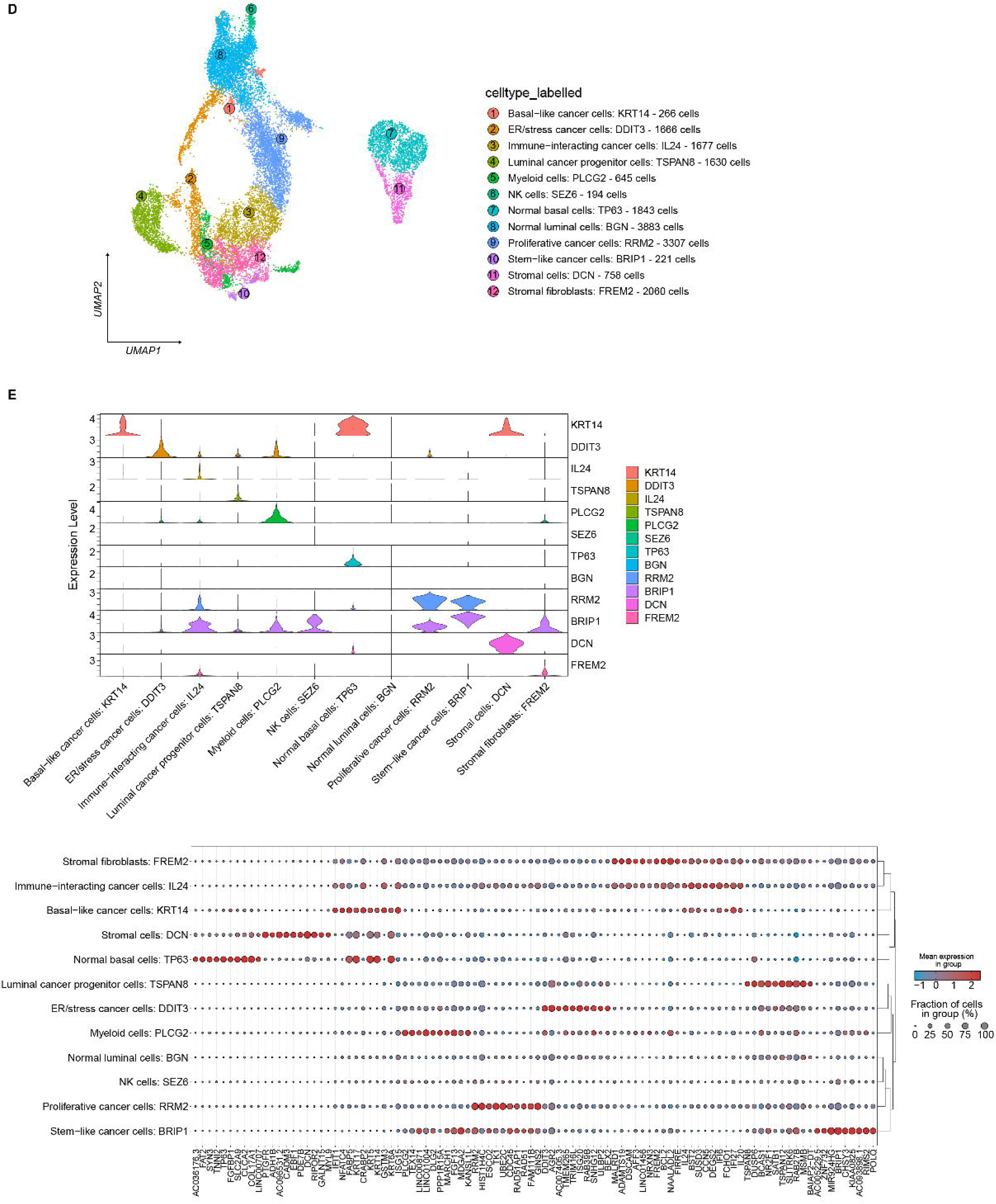
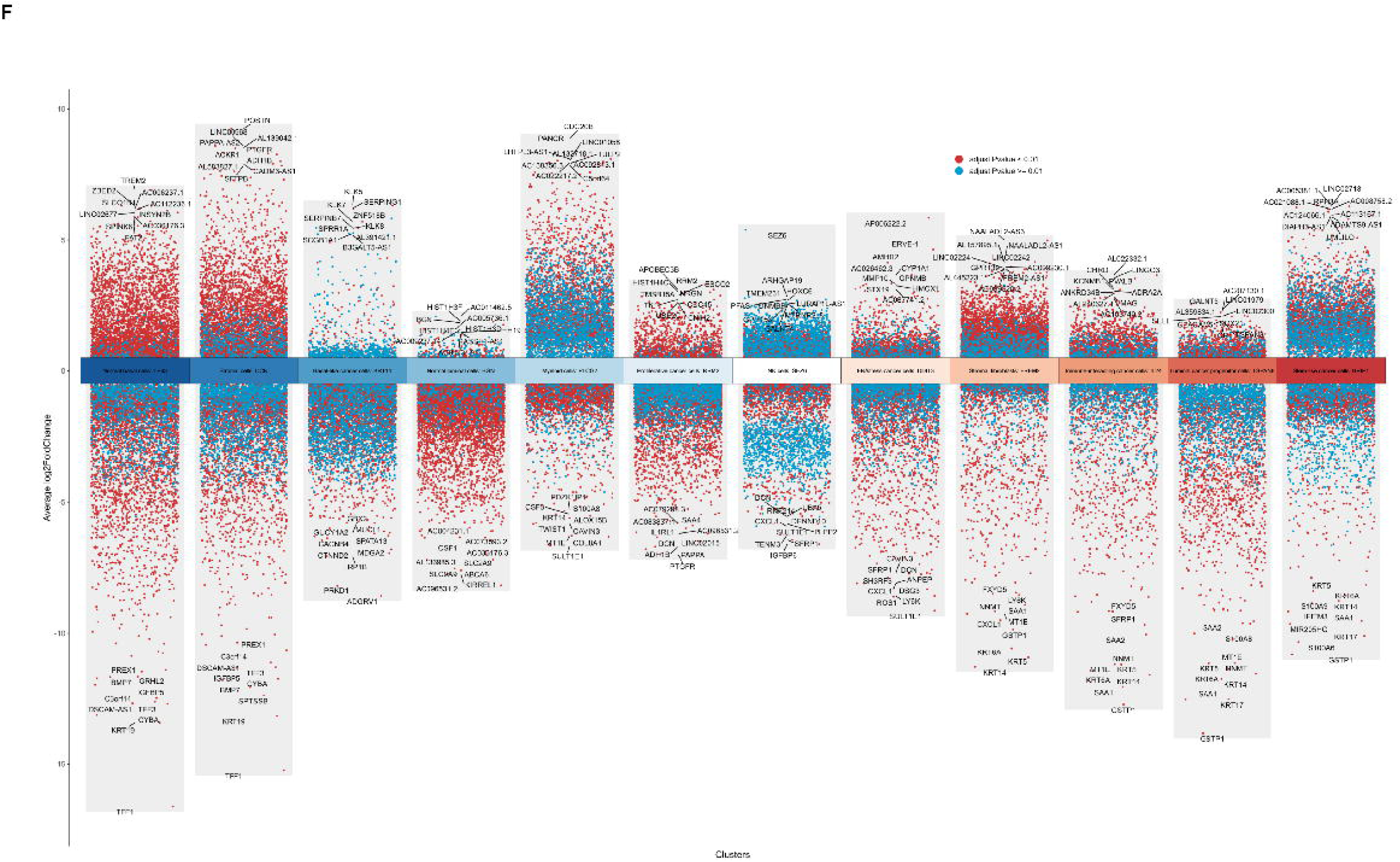
Comprehensive single-cell transcriptomic analysis of ER+ breast cancer samples. ***A***, UMAP demonstrates before-batched (*top*) and after-batched (*bottom*) batch correction, showing clustering by condition type (*left*, NT: normal tissue, PT: primary tumor, RT: TAM-resistant tumor) and individual patient samples (*right*). ***B***, UMAP visualization showing 14 different cell types. ***C***, Multi-layered visualization of cell clusters across different sample types. Of the 14 identified cell clusters, only 7 represent all three tissue types (normal, primary tumor, and TAM-resistant tumor). Segments 1 and 10 were similar and subsequently merged for downstream analysis. ***D***, UMAP plot displays 12 cell types in breast tissue and breast cancer, with each dot representing a single cell and colors indicating different cell populations. ***E*,** *Top*, A violin plot of gene expression levels across different cell types. *Bottom*, A gene expression heatmap showing different cell types in cancer and normal tissue. ***F***, Volcano plots reveal differential gene expression patterns specific to each cell type, highlighting key genes with significant expression changes.

**Fig. S13.**
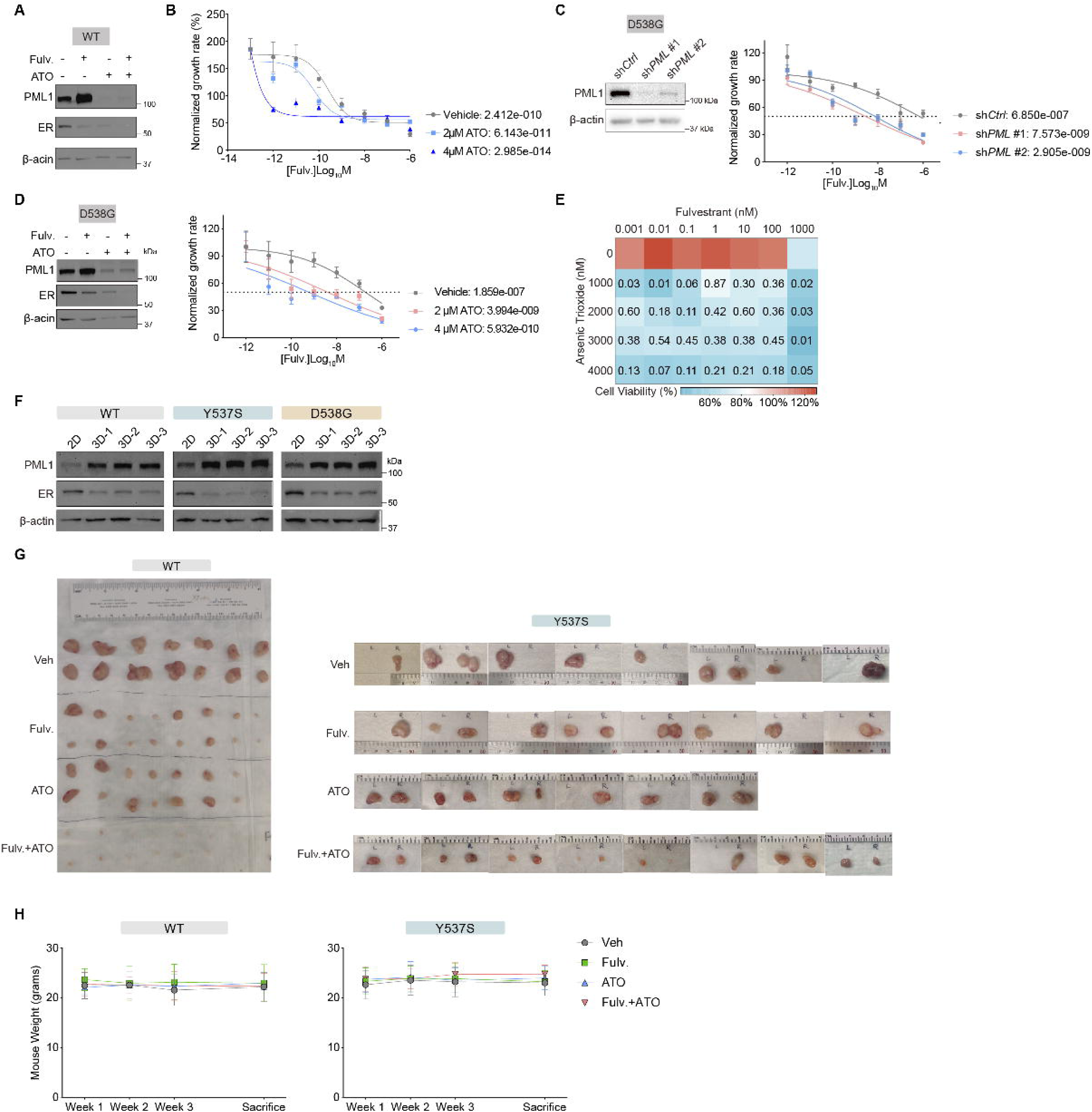
ATO enhances Fulv’s anti-cancer efficacy in ^WT^ER- and ^D538G^ER-KI MCF-7 cells. ***A-B***, ATO treatment significantly reduces PML1 protein abundance (***A***) and enhances Fulv’s anti-proliferative effects in ^WT^ER-KI cells (***B***). ***C-D****, PML* knockdown (***C***) or ATO treatment (***D***) potentiates Fulv’s inhibitory activity in ^D538G^ER-KI cells. ***E***, Drug synergy analysis of Fulv/ATO combinations in ^D538G^ER-KI cells using the Chou-Talalay method with Combination Index (CI) values. ***F***, Elevated PML1 protein levels in ^WT^ER-KI, ^Y537S^ER-KI, and ^D538G^ER-KI MCF-7 cells grown as 3D spheroids versus 2D cultures. ***G***, Representative images of tumor responses in mice bearing MCF-7 ^WT^ER-KI (*A*) or ^Y537S^ER-KI (*B*) xenografts following treatment with vehicle control, Fulv, ATO, or combination therapy. ***H***, Body weight measurements of mice throughout the experimental period.

**Fig. S14.**
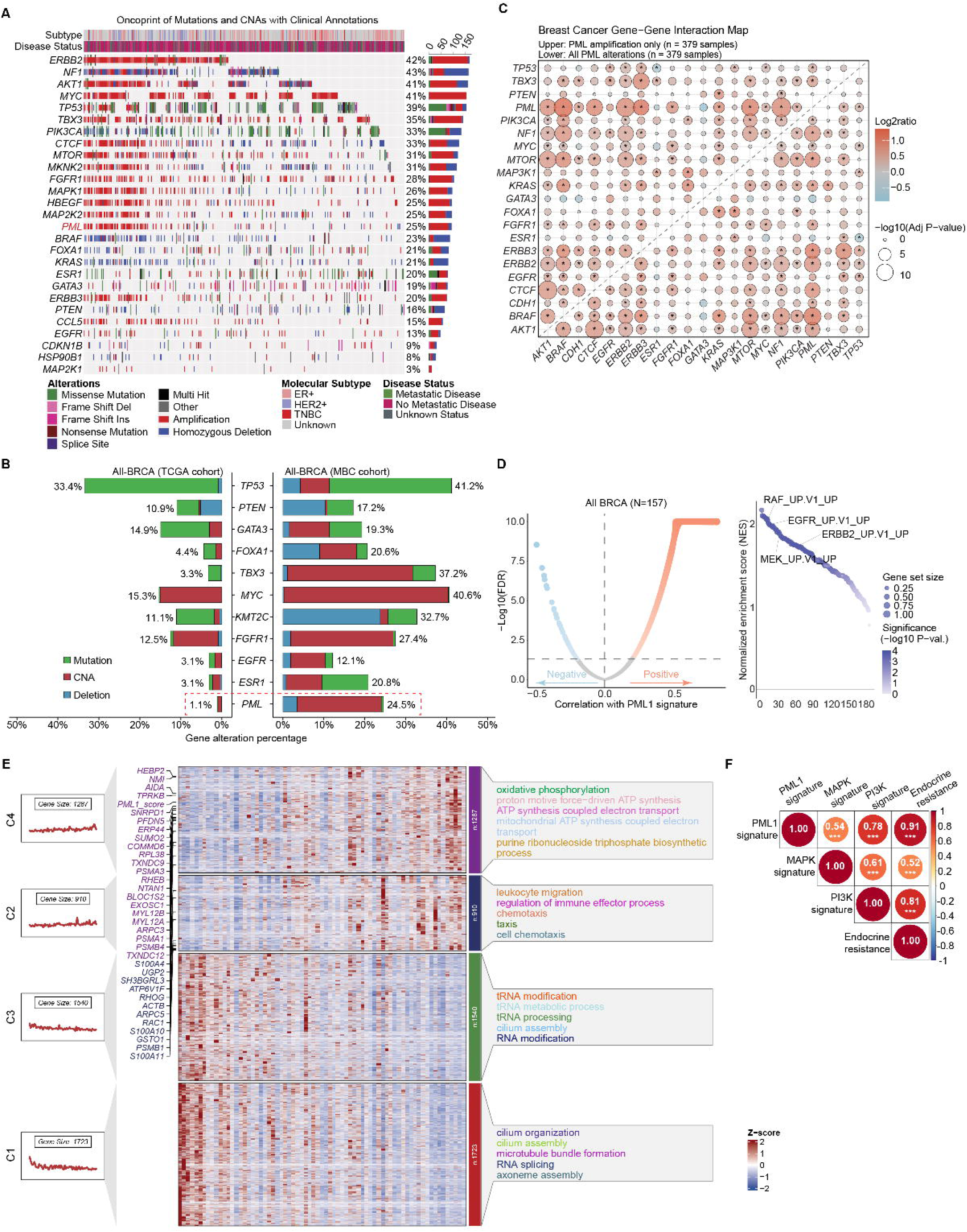
*PML* gene alterations in MBC. ***A,*** Oncoplots showing *PML* gene amplification in MBC tumors (N=379). ***B,*** Comparative analysis of gene mutation and amplification frequencies between MBC (N=379) and TCGA primary tumors (N=1084), showing significantly higher rates in all MBC. ***C,*** Gene-gene interaction map for MBC tumors reveals significant co-occurrence between *PML* amplification/alterations and *AKT1*, *BRAF*, *CTCF*, *ERBB2*, *ERBB3*, *FGFR1*, *KRAS*, *MTOR*, and *NF1*. ***D,*** *Left*, Volcano plot for a cohort of MBC patients. The x-axis shows correlation values between the PML1 gene signature and other mRNAs, and the y-axis shows statistical significance as -Log10(FDR). *Right*, Oncogenic gene set enrichment analyses show that the PML1 gene signature is most significantly enriched in RAF, EGFR, MEK, and ERBB2 signaling in ER+ MBC. *Right*, Significant associations between PML1 gene signature and RAF, EGFR, ERBB2, and MEK oncogenic signatures in MBC. ***E,*** Heatmap of gene expression data organized into four clusters (C1, C2, C3, and C4) with associated biological processes and PML1 signature genes indicated. ***F,*** Strong correlation of PML1 gene signature and PI3K, MAPK, and endocrine resistance signature in all MBC cases.

**Fig. S15.**
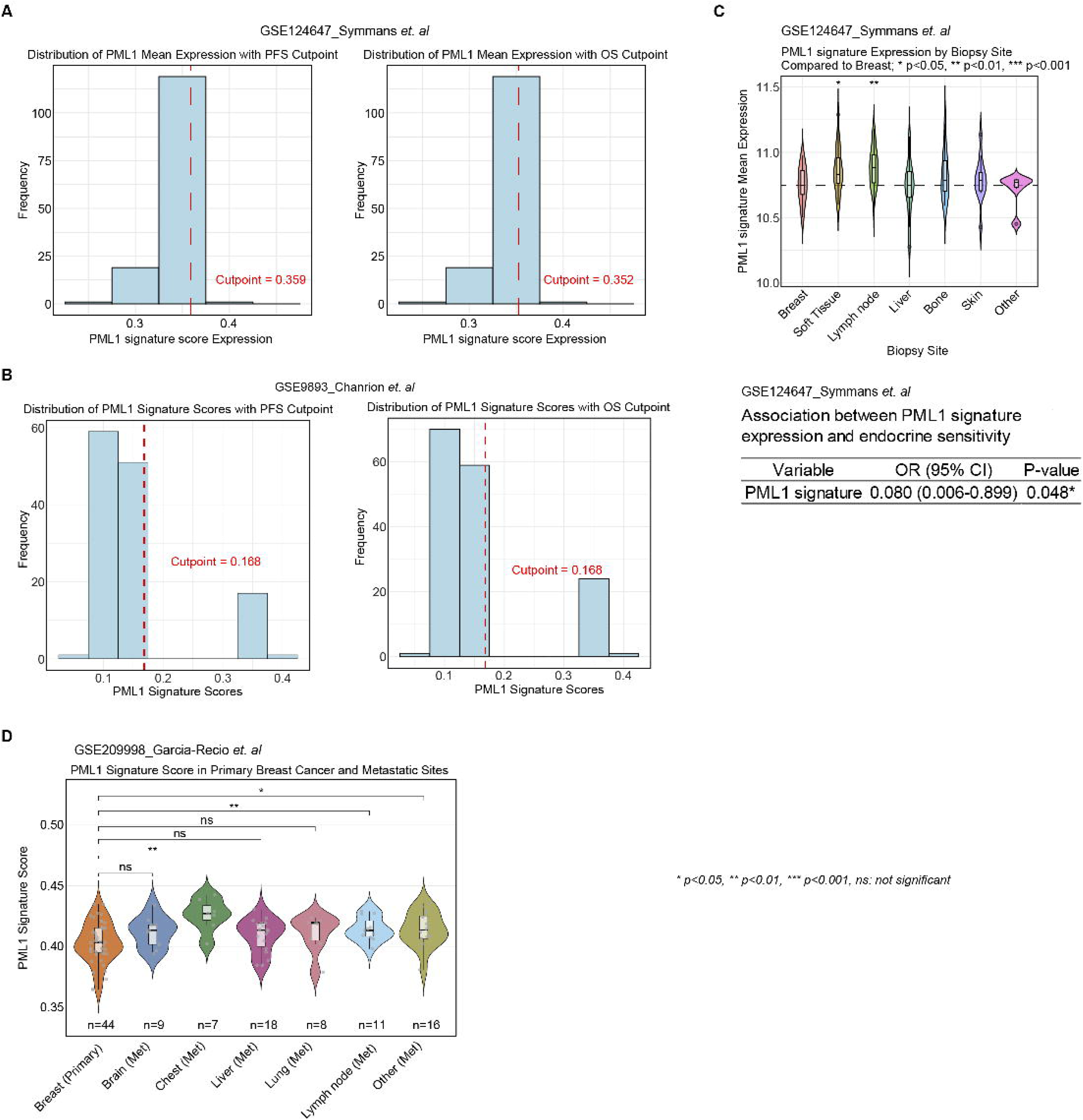
Clinical information for Fig. 5G-H. ***A,*** Histogram distribution of PML1 gene signature expression scores in the GSE124647 dataset with cutpoints for Progression-Free Survival (PFS, 0.359, *left*) and Overall Survival (OS, 0.352, *right*). ***B***, Histogram distribution of PML1 gene signature scores in GSE9893 dataset with cutpoints for PFS and OS (both 0.168). ***C***, Violin plots from the GSE124647 dataset showing PML1 signature expression across different metastatic sites compared to breast tissue. The association between PML1 gene signature scores and endocrine sensitivity is shown. ***D,*** Violin plots from the GSE209998 dataset comparing PML1 gene signature scores across primary breast cancer and various metastatic sites.

**Fig. S16.**
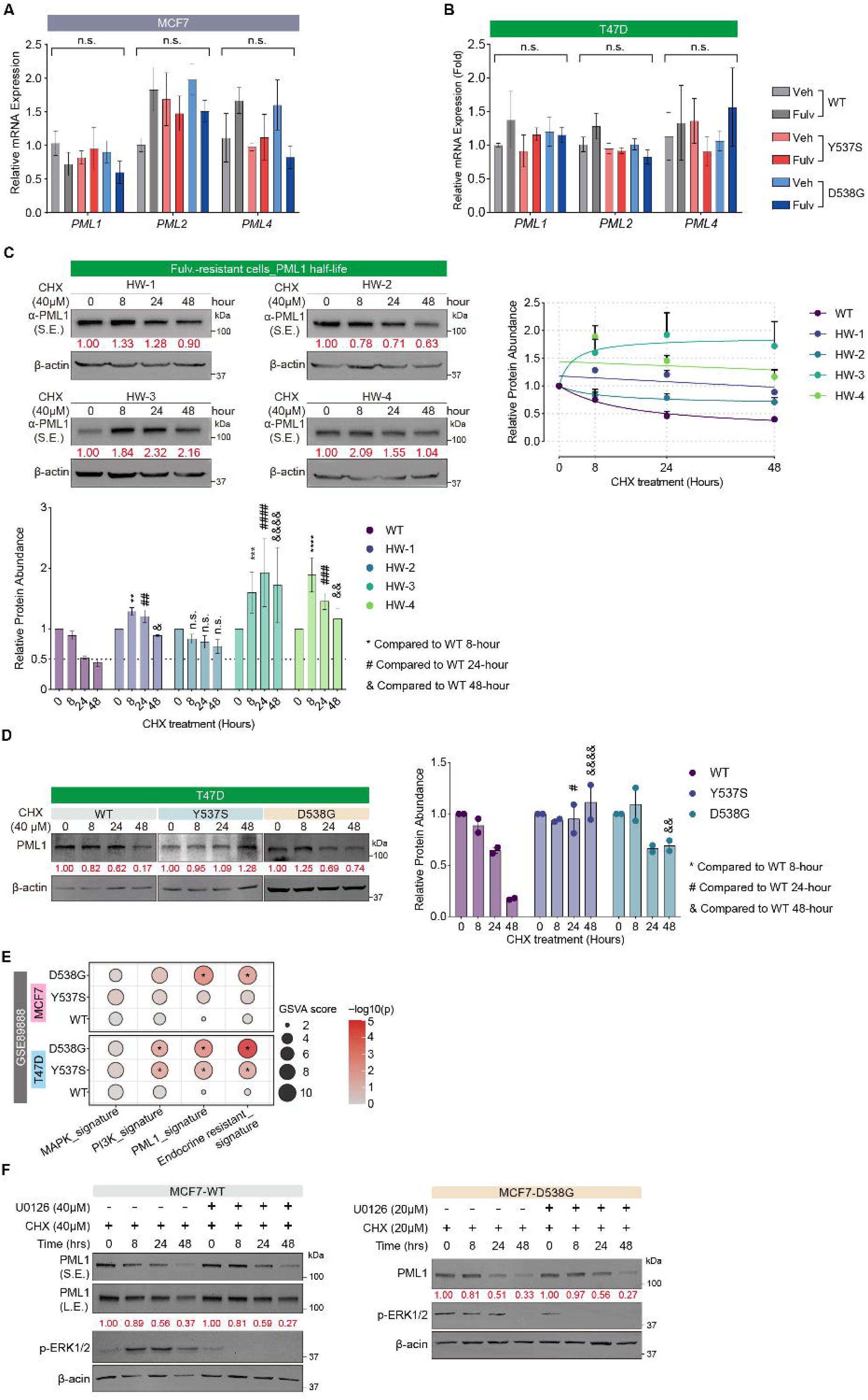
Fulvestrant extends PML1 protein half-life in resistant cells. ***A,*** RT-qPCR analysis showing comparable *PML* isoform (*PML1*, *PML2*, *PML4*) expression across all MCF-7 knock-in variants with or without Fulv treatment. ***B,*** Fulv induces PML1 protein levels in T47D KI variants without altering PML isoform mRNA expression. ***C-D***, Extended PML1 protein half-lives in Fulv-resistant MCF-7 variants (***C***) and T47D (***D***) cells. ***D,*** Extended half-lives of PML1 protein in T47D cell variants. ***E***, Dot plot visualization of gene signature enrichment analysis shows enhanced PI3K, MAPK, PML1, and endocrine resistance gene signatures in ^Y537S^ER-KI and ^D538G^ER-KI MCF-7 and T47D cell variants compared to ^WT^ER-KI cells. ***F,*** ERK inhibitor U0126 shows minimal effect on PML1 protein stability in ^WT^ER- or ^D538G^ER-KI MCF-7 cells.

**Fig. S17.**
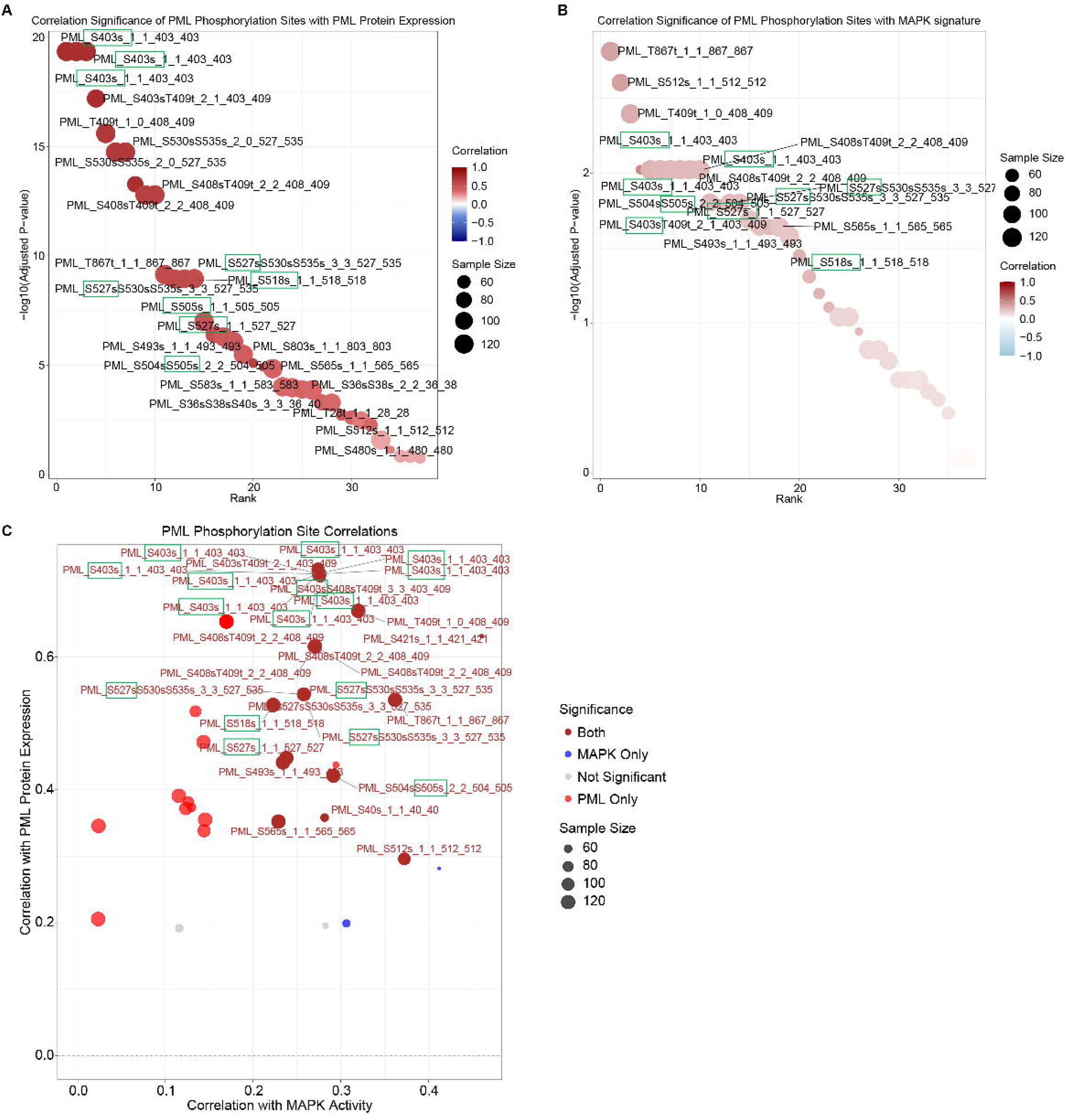
PML phosphorylation correlates with MAPK pathway activation in breast tumors. ***A***, Correlation analysis between PML protein abundance and site-specific phosphorylation. ***B***, Correlation between MAPK gene signature scores and phosphorylated PML peptides. ***C***, Scatter plot visualizing relationships between different PML phosphorylation sites with both PML protein expression (y-axis) and MAPK activity (x-axis) in breast tumors (N>90), with adjusted *p*-vale < 0.05.

**Fig. S18.**
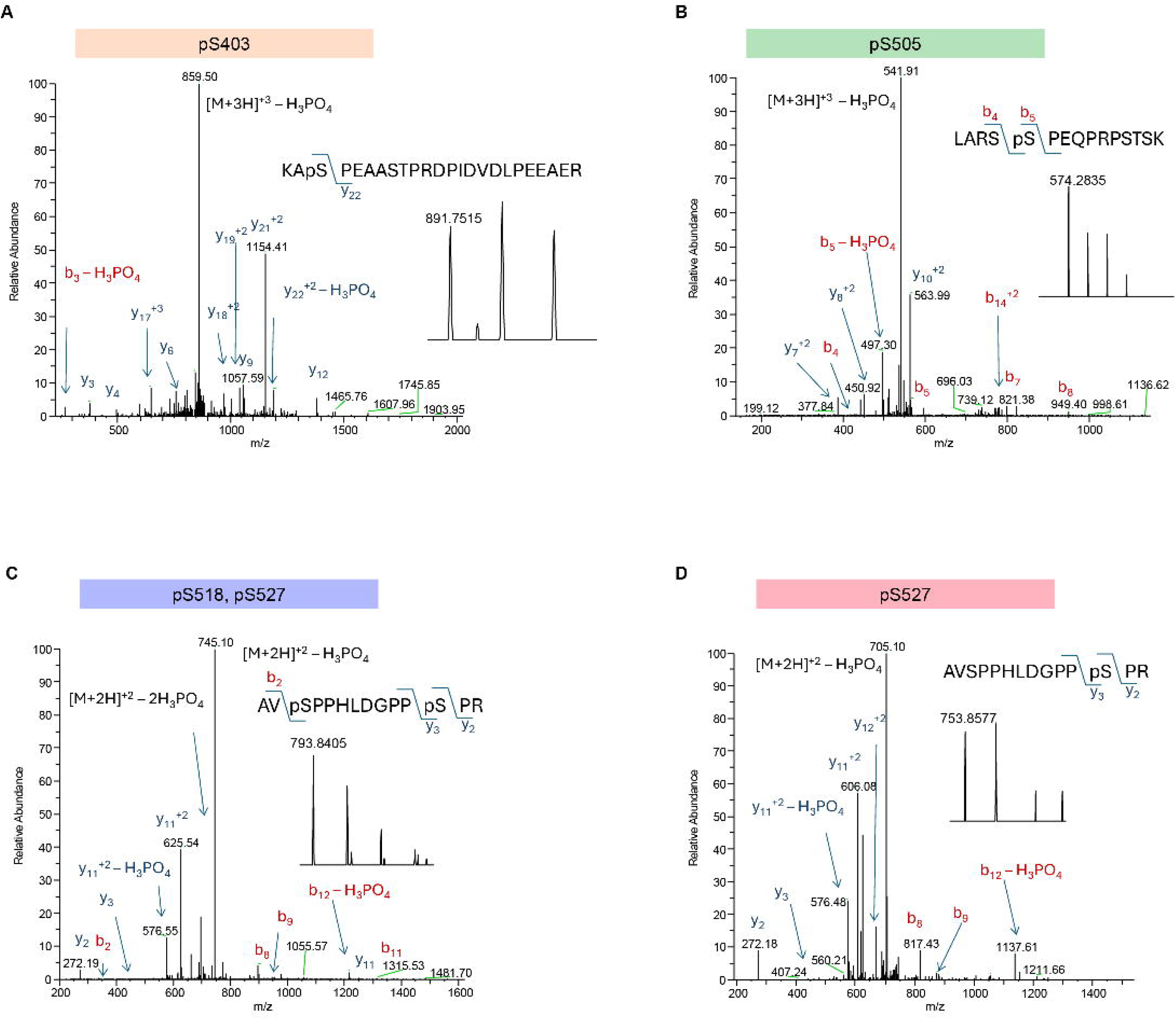
Mass spectrometry identification of PML1 phosphorylation sites. Proteins isolated from a PML IP experiment were digested with trypsin, and the digest was analyzed by LC-MS/MS. ***A***, Phosphorylation at S403 on peptide (401)KA**S**PEAASTPRDPIDVDLPEEAER(424). ***B***, Phosphorylation at S505 on peptide (501)LARS**S**PEQPRPSTSK(515) ***C,*** Dual phosphorylation at S518 and S527 on peptide (516)AV**S**PPHLDGPP**S**PR(529). ***D,*** Single phosphorylation at S527 on the same peptide region.

**Fig. S19.**
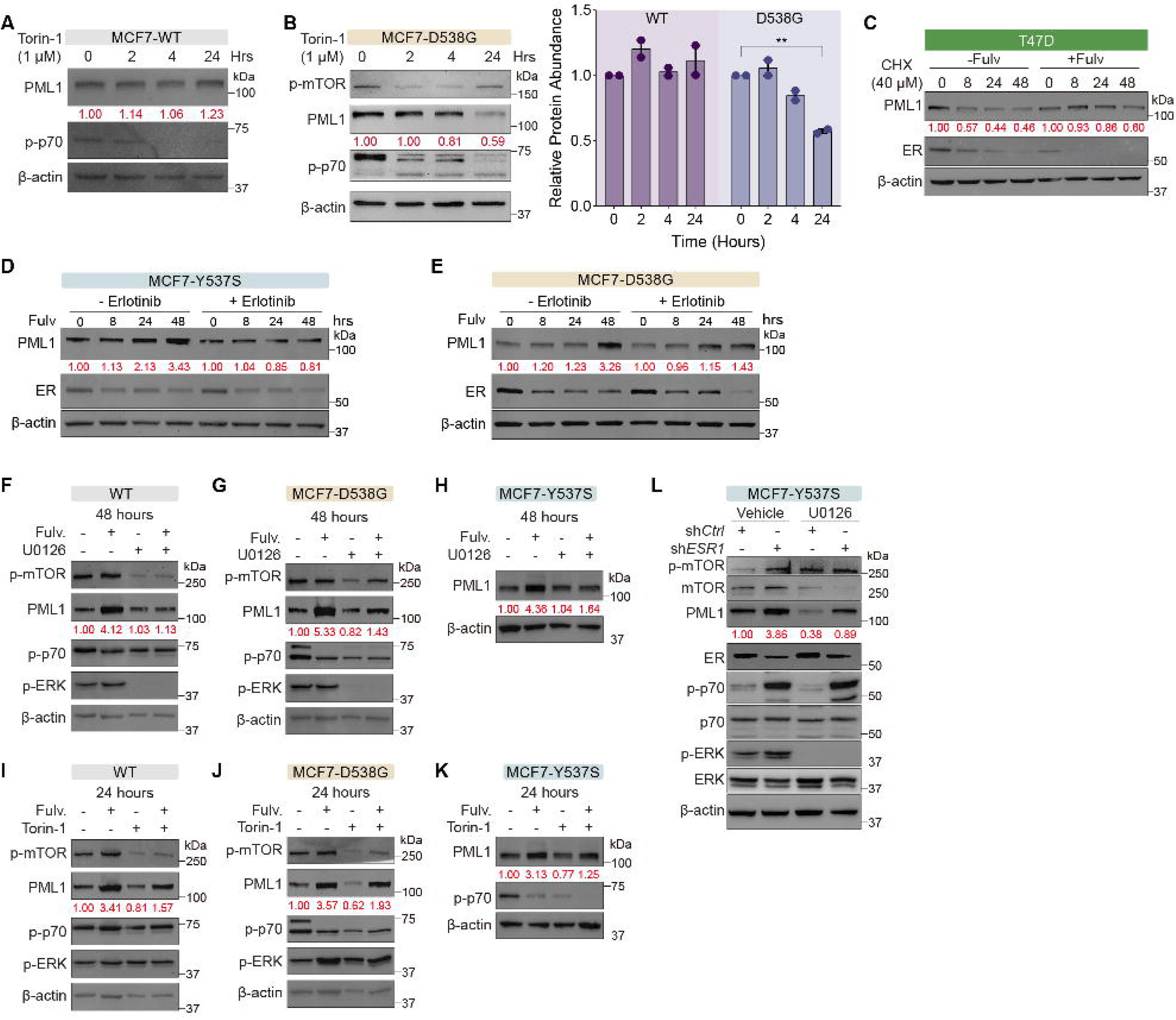
Differential regulation of PML1 protein by signaling pathway inhibitors. ***A-B***, mTOR inhibitor Torin-1 minimally affects PML1 levels in ^WT^ER-KI cells (***A***) but substantially reduces PML1 in ^D538G^ER-KI cells (***B***). ***C***, Fulv treatment prolongs PML1 protein half-life in T47D cells. ***D-K,*** Inhibition of Fulv-induced PML1 accumulation by: erlotinib in Y537SER-KI (***D***) and D538GER-KI cells (***E***); MEK inhibitor U0126 in ^WT^ER-KI (***F***), ^Y537S^ER-KI (***G***), and ^D538G^ER-KI cells (***H***); and Torin-1 in ^WT^ER-KI (***I***), ^Y537S^ER-KI (***J***), and ^D538G^ER-KI cells (***K***). ***L,*** U0126 blocks *ESR1* knockdown-induced PML1 protein accumulation in ^Y537S^ER-KI cells.

**Fig. S20.**
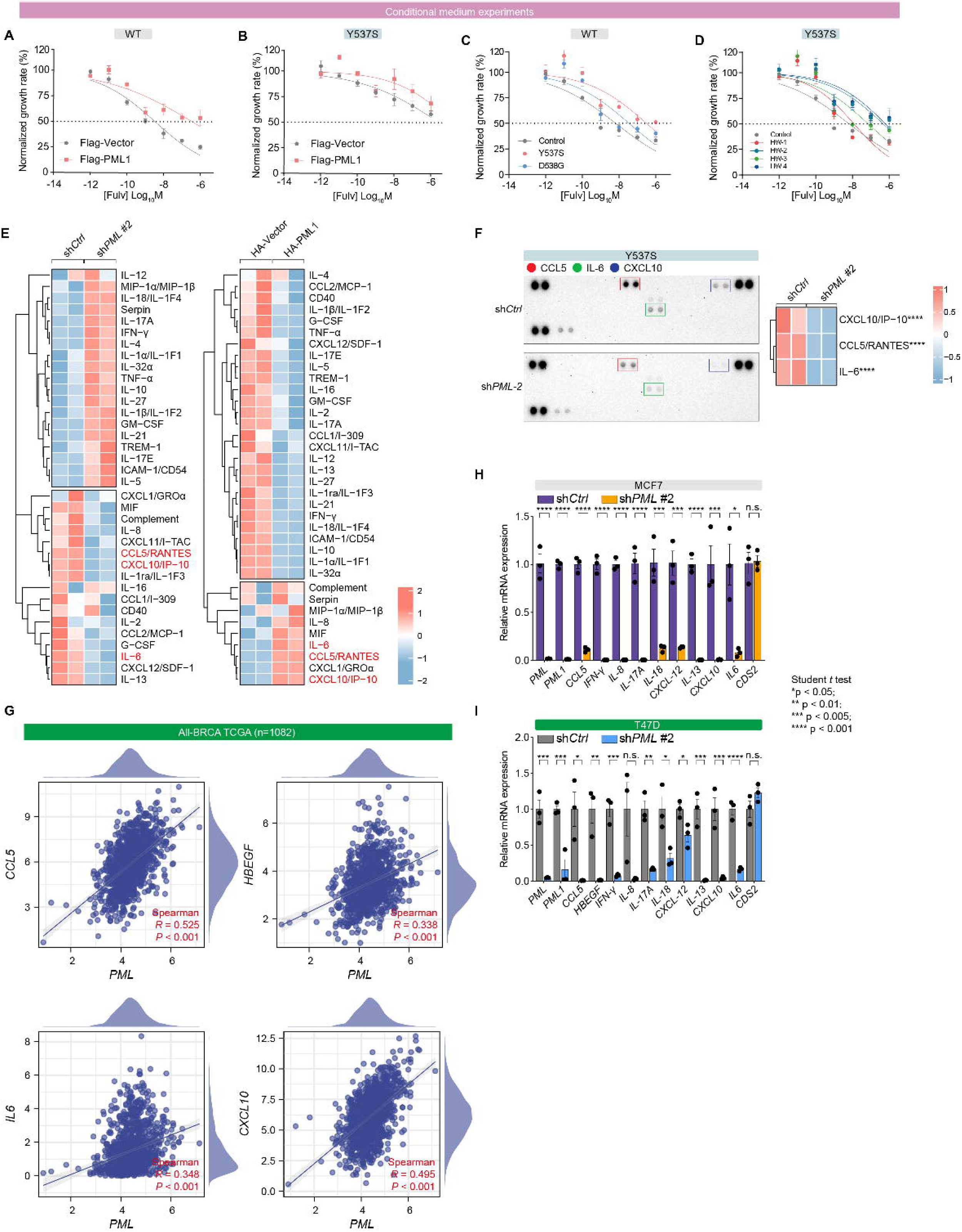
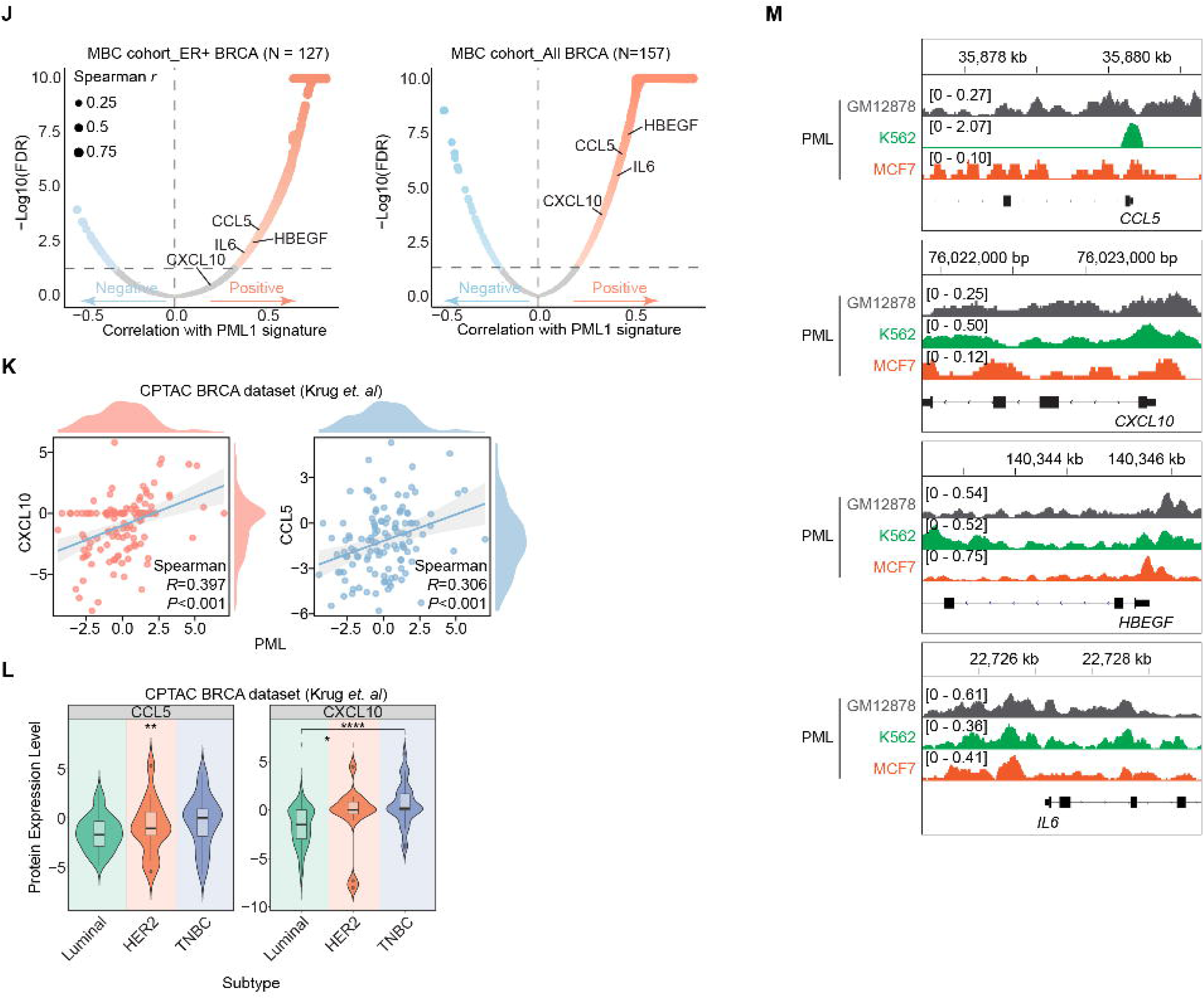
PML1 regulates inflammatory cytokine expression in endocrine-resistant breast cancer. **A-D**, Conditioned media (CM) from PML1-overexpressing cells confers Fulv resistance in ^WT^ER- (***A***) and ^Y537S^ER-KI MCF-7 (***B***), with CM from ^Y537S^ER- and ^D538G^ER-KI (***C***) and in-house Fulv-resistant MCF-7 variants (***D***) increasing Fulv IC_50_ compared to CM from ^WT^ER-KI controls. ***E***, Cytokine array heatmaps demonstrating PML’s regulatory role in cytokine expression profiles. ***F****, PML* knockdown in ^Y537S^ER-KI cells reduces secreted CCL5, IL-6, and CXCL10 in CM. ***G***, Positive correlation between *PML* mRNA levels and cytokines *CCL5*, *HBEGF*, *IL6*, and CXCL10 expression in breast tumors. ***H-I,*** *PML* knockdown significantly decreases the expression of inflammatory cytokines mRNAs in MCF-7 (***H***) and T47D (***I***) cells. ***J***, RNA-seq analysis shows a correlation between *HBEGF*, *CCL5*, *IL6*, and *CXCL10* expression and the PML1 gene signature in ER+ (*left*) and all MBC (*right*). ***K***, CPTAC analysis shows positive correlations between PML and CXCL10 and between PML and CCL5 protein levels in breast tumors. ***L***, CCL5 and CXCL10 protein levels are significantly higher in TNBC than in luminal breast cancer. ***M***, Genomic tracks showing binding of PML to the *CCL5*, *HBEGF*, *IL6*, and *CXCL10* promoter regions.

## Supplementary Tables

**Table S1.** List of PML1-bound gene promoters as well as genes and proteins whose expression correlate with PML1 mRNA and total PML protein levels, respectively.

**Table S2.** Annotation of signature genes related to PML1, PI3K, MAPK, and endocrine resistance.

**Table S3.** Primer sequences for RT-qPCR and ChIP-qPCR.

## Authors’ Disclosures

No disclosure to report.

## Acknowledgments

We thank Drs. Rinath Jeselsohn and Myles Brown (Dana-Farber Cancer Institute) for providing the ER mutant knock-in MCF-7 cell variants, Drs. Steffi Oesterreich and Adrian Lee (University of Pittsburgh) for the ER mutant knock-in T47D cell variants, and Dr. John Pink for MCF-7 2A and 5C cell variants. This work made use of the High Performance Computing Resource in the Core Facility for Advanced Research Computing at Case Western Reserve University. The Fusion Lumos instrument was purchased via an NIH shared instrument grant, 1S10OD023436-01.

## Funding

This work was supported by the American Cancer Society grant DBG-24-1314754 (HYK) and by National Institutes of Health, R01GM114056 (SY and HYK), R01CA257502 (RAK), (R01CA288626) JAD, the Case Comprehensive Cancer Center (P30CA043703). ZY, XL, and YC were supported as Hanson Research Summer Scholars.

## Authors’ contributions

**HYK** conceived and designed the study. **HYK** designed in-cell experiments executed by **HW, ZY, YC, CW, ZL, XL, CP, GL, JY, and JC**. **HW** and **HYK** designed xenograft studies using NOD/SCID mice, executed by **HW** and **YC**. **HW** conducted all bioinformatics and clinical data analyses. Toxicity studies using NSG mice were under the guidance of **RAK**. **HYK** wrote the original draft of the manuscript. All authors contributed to the editing and review of the manuscript and approved the final version.

## Competing interests

N/A.

## Notes

### Competing Interest Statement

The authors have declared no competing interest.

